# Dual-targeted NADK2 Links Mitochondrial Redox Homeostasis to Carbon Partitioning and Heterotrophic Growth in *Chlamydomonas reinhardtii*

**DOI:** 10.64898/2026.08.14.744979

**Authors:** Neda Fakhimi, Michelle Meagher, Justin Findinier, Ugo Cenci, Andrey Malkovskiy, Dimitri Tolleter, Masayuki Onishi, Adrien Burlacot, Nanette Boyle, Arthur Grossman

## Abstract

NAD kinases (NADKs) can modulate NADP(H) levels across shifting environmental conditions. In *Chlamydomonas reinhardtii*, we localized CreNADK1 to the chloroplast, CreNADK3 to the cytoplasm, and CreNADK2 to both the chloroplast and mitochondria. Importantly, during mixotrophic (light + acetate) and photoautotrophic (light) growth, most CreNADK2 is chloroplast localized whereas an increased proportion resides in mitochondria when the cells are grown heterotrophically (dark + acetate). In Cre*nadk2* mutants, a diminished ability to synthesize mitochondrial NADP(H) inhibits the TCA cycle and markedly retards heterotrophic growth. In the light, photosynthesis becomes the primary energy source, and the presence of chloroplastic CreNADK1 enables near-normal growth in mutant cells, with inhibition of TCA cycle activity partially ameliorated via rerouting of acetate metabolism through the glyoxylate shunt, bypassing the TCA cycle and diminishing mitochondrial ROS production. These findings demonstrate spatial plasticity of NAD(H)Ks and the implementation of ‘bypass’ metabolism to enable growth of Cre*nadk2* mutants.

## INTRODUCTION

Nicotinamide adenine dinucleotide (NAD(H)) and its phosphorylated form (NADP(H)) are essential coenzymes that serve as critical electron carriers in numerous metabolic processes. NAD(H) primarily functions in catabolic reactions whereas NADP(H) is predominantly involved in anabolic processes, particularly biosynthetic pathways and antioxidant defense systems (Agledal et al., 2010; Gakière et al., 2018; Lu et al., 2025; Outten & Culotta, 2003; Pollak et al., 2007; Ziegler, 2000). Interconversion between NAD(H) and NADP(H) is catalyzed by NAD kinases (NADKs), which help cells adjust the levels of these cofactors for driving cellular metabolism and ameliorating oxidative stress (Pollak et al., 2007). Regulating NAD(P)(H) pools is fundamental for controlling redox homeostasis, energy metabolism, and facilitating/controlling various biosynthetic pathways (Kawai & Murata, 2008).

NADKs among different organisms exhibit diverse substrate specificities. Based on substrate preferences, they can be classified as NAD^+^-specific (phosphorylating only NAD^+^), NADH-specific (phosphorylating only NADH), or dual-specific (phosphorylating both NAD^+^ and NADH) (Mori et al., 2005). Substrate specificity of NADKs may play a significant role in determining the metabolic flexibility of an organism, particularly in response to dynamic environmental conditions and cellular energy demands.

Prokaryotes and eukaryotes differ significantly in their NADK systems. Most prokaryotes, including heterotrophic bacteria, possess a single NADK that varies in its substrate specificities (Barbosa et al., 2023). Cyanobacteria, which are oxygenic photosynthetic prokaryotes, generally contain two distinct NADKs that reflect their complex metabolism and the need to integrate photosynthesis and carbon fixation into diverse biosynthetic pathways (Gao & Xu, 2012). In contrast, eukaryotes generally have more complex NADK systems. *Arabidopsis thaliana* (hereafter *Arabidopsis*) has four distinct NADKs with AtNADK1 in the cytosol, AtNADK2 in the chloroplasts, AtNADK3 in peroxisomes, and AtNADKc associated with the outer mitochondrial membrane (Dell’Aglio et al., 2019; Turner et al., 2005). The budding yeast *Saccharomyces cerevisiae* (hereafter *Saccharomyces*) contains three NADKs with distinct subcellular localizations: two in the cytosol (Utr1p and Yef1p) and one in the mitochondria (Pos5p) (Miyagi et al., 2009; Shi et al., 2005). Pos5p exhibits remarkably high NADH kinase activity and generates mitochondrial NADPH through either direct phosphorylation of NADH or phosphorylation of NAD^+^ with subsequent reduction by mitochondrial NADP^+^-dependent dehydrogenases, particularly acetaldehyde dehydrogenase (Ald4p) (Miyagi et al., 2009; Outten & Culotta, 2003). Human cells possess two NADKs, a cytosolic and a mitochondrial enzyme (Ohashi et al., 2012). The human mitochondrial enzyme can directly phosphorylate NADH to generate NADPH, which can alleviate mitochondrial oxidative stress (Ohashi et al., 2012). Compartmentalization of NADK enzymes in eukaryotic cells highlights the evolutionary importance of maintaining distinct NADP(H) pools that support organelle-specific metabolic requirements and redox defense mechanisms (Miyagi et al., 2009).

While our understanding of NADKs in land plants and heterotrophic organisms has advanced significantly, knowledge about these enzymes in algae remains limited. Microalgae are particularly intriguing subjects for exploring NADK function and regulation; despite being photosynthetic like land plants, algae demonstrate remarkable metabolic versatility and rapidly acclimate to environmental changes. These organisms also have considerable biotechnological value, with NADPH serving as a critical cofactor that controls numerous biosynthetic pathways relevant to the production of biofuel, pharmaceuticals, nutraceuticals and other high-value metabolites (Schneider et al., 2025; H. Zhang et al., 2025). The unicellular green microalga *Chlamydomonas reinhardtii* (hereafter *Chlamydomonas*) has served as a model organism for studying mechanisms associated with a diverse pallet of metabolic processes, including photosynthesis, pigment biosynthesis, central carbon metabolism, the carbon concentrating mechanism (CCM), and nutrient uptake and assimilation.

According to the latest annotation in Phytozome (https://phytozome-next.jgi.doe.gov/), the *Chlamydomonas* genome encodes three predicted NADKs that we have designated CreNADK1 (Cre07.g322950), CreNADK2 (Cre10.g431650) and CreNADK3 (Cre12.g560600). Among these, we focused this study on CreNADK2. Our phylogenetic analysis revealed that CreNADK2 belongs to a specialized clade of NAD kinases that is exclusive to microorganisms, a group for which no representative functional studies currently exist. Characterization of CreNADK2 and mutants disrupted for the gene encoding this enzyme have allowed us to identify its dynamic subcellular location and the role it plays in growth, redox balance, partitioning of metabolites, and metabolic control.

## RESULTS AND DISCUSSION

### Phylogenetic Divergence and Lineage-Specific Adaptation

Vickman and Erives (2019) demonstrated that the eukaryotic NADK repertoire is ancient and has undergone episodic evolution during major evolutionary transitions. Phylogenetic analyses have revealed that nearly all eukaryotic *NADK* genes belong to one of two ancient sister clades, cytosolic (cyto) and mitochondrial (mito) (Vickman & Erives, 2019). Our phylogenetic analysis of *NADK* genes encompasses a wide range of taxa including eukaryotic groups such as Chloroplastida, Rhizaria, Rhodophyta, Glaucophyta, Stramenopiles, Alveolata, and Opisthokonta (including *Homo sapiens*, *Mus musculus*, *Drosophila melanogaster, and Saccharomyces cerevisiae*), as well as prokaryotic groups including heterotrophic bacteria and photosynthetic cyanobacteria (**Fig. 1A**). Phylogenetic analyses of our study and the work of Vickman and Erives have demonstrated that the eukaryotic cytosolic (cyto) clade is comprised of three distinct subclades. *Chlamydomonas* possesses three *NADK* genes within the broad eukaryotic cyto-clade (bootstrap support = 62), with each gene falling into one of the three identified cyto-subclades. The phylogenetic patterns suggest that all the *CreNADKs* originated from the genome of the host organism associated with the primary endosymbiosis. This distinguishes them from the NADKs in heterotrophic bacteria and photoautotrophic cyanobacteria (**Fig. 1A**). Furthermore, these results show that CreNADK proteins align with those of other eukaryotes, including members of the fungal kingdom, unranked amoebozoans, and members of Chloroplastida, in which the ancestral mito-clade *NADK* gene has been lost. However, *Chlamydomonas* is distinctive in possessing an additional *NADK* gene, Cre*NADK2*, belonging to the third cytosolic subclade, a gene absent in plants. (Vickman & Erives, 2019).

**Fig. 1.**
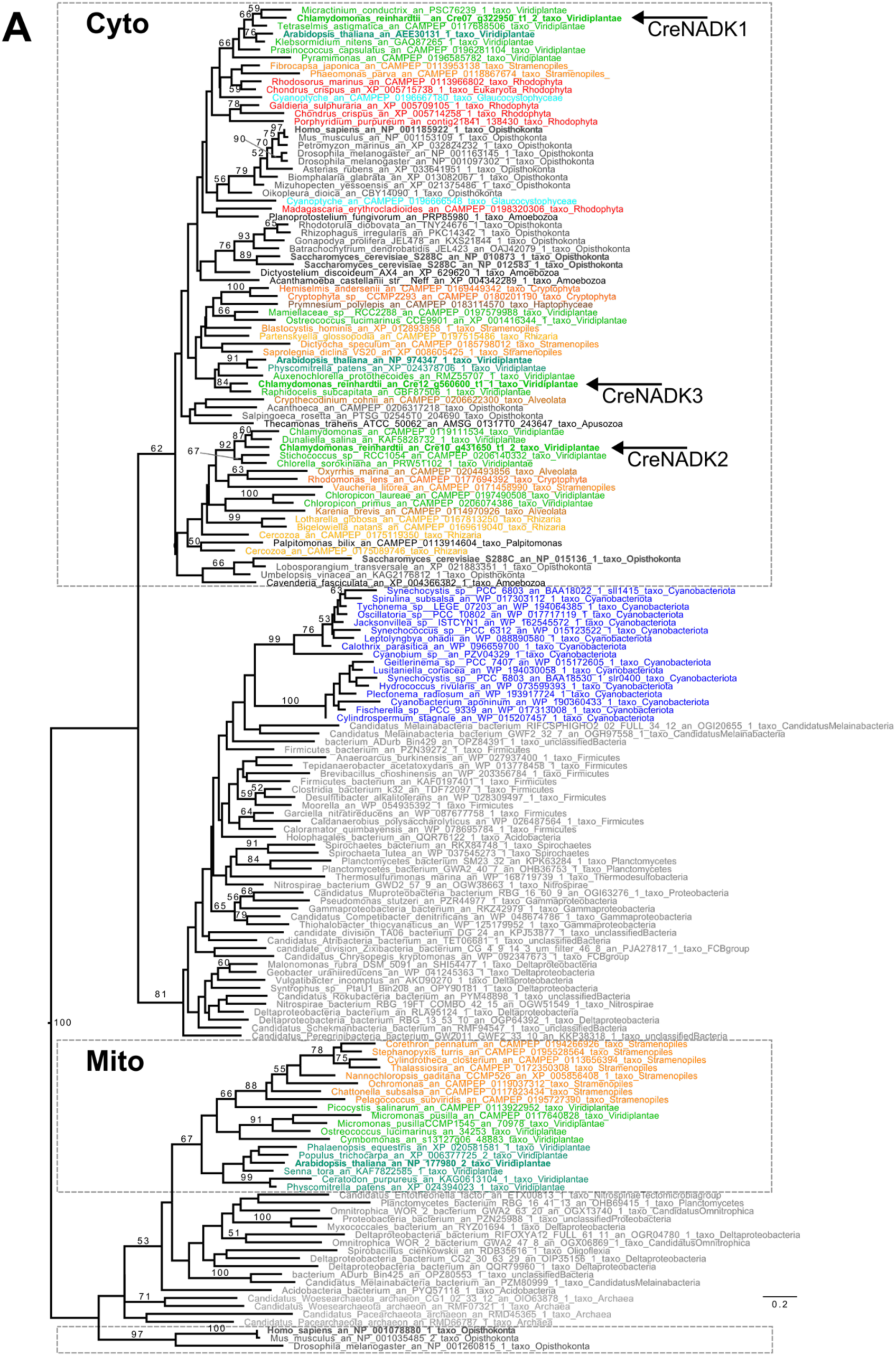
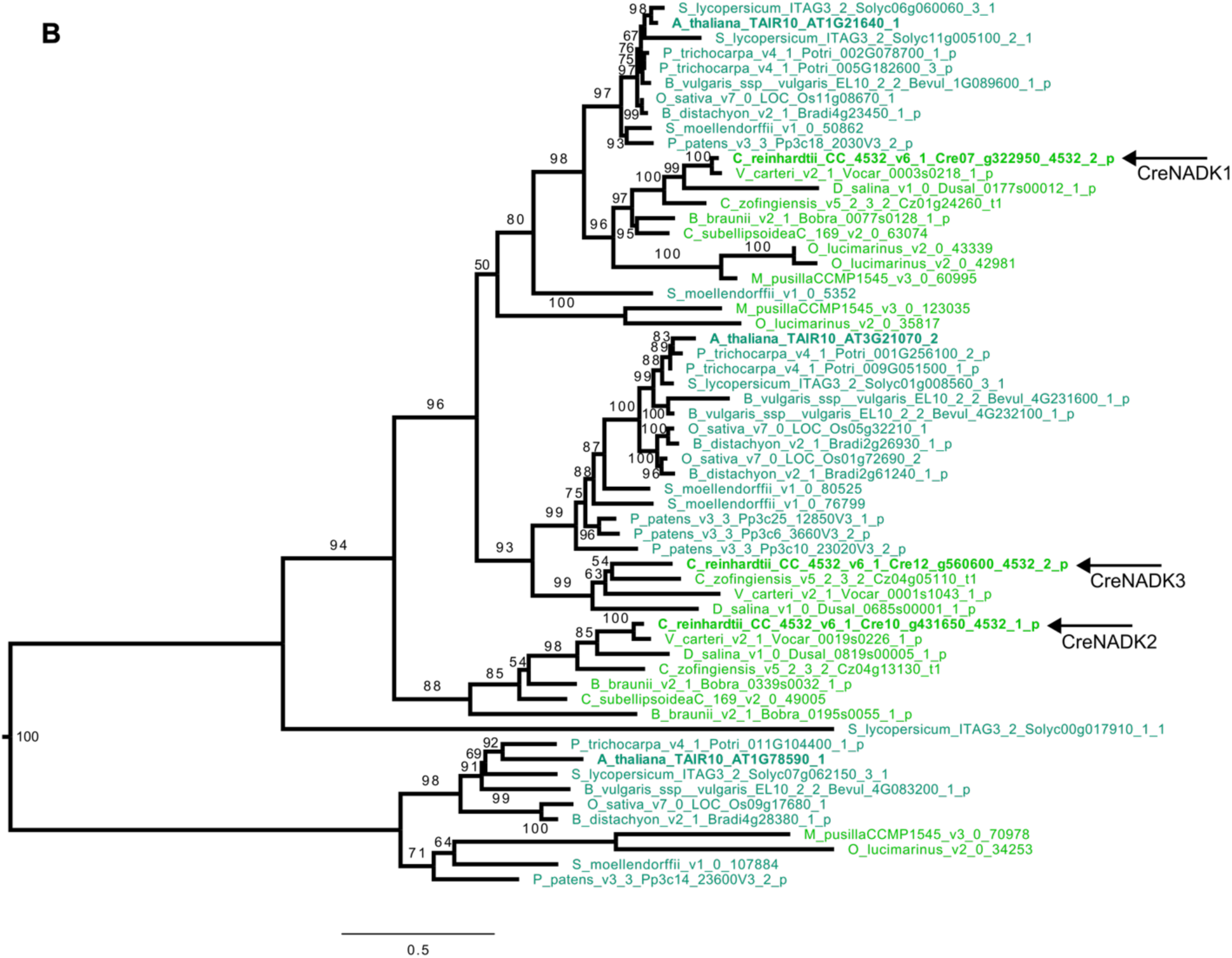
NADK phylogenies. (**A**) Phylogenetic tree of a large selection of NADKs. Sequences from land plants are in dark green whereas sequences from other Chloroplastida are in light green. Colors representing sequences from other organisms are: Rhizaria in yellow, Rhodophyta in red, Glaucophyta in light blue, Cyanobacteria in dark blue, Stramenopiles in orange, Alveolata in brown, Opisthokonta in dark grey, other Eukaryota in black, and Bacteria in light grey. The scale bar indicates the number of inferred substitutions per amino acid per site. Values for nodes are drawn from 100 bootstrap repetitions and only values greater than 50 are indicated at the nodes. NADKs from *Chlamydomonas*, indicated by arrows, belong to the cyto-clade [group with various eukaryotes; no prokaryotes]. The cytosolic (cyto) and mitochondrial (mito) clades are enclosed by gray dashed boxes. (**B**) Phylogenetic tree of NADK constructed from complete genome sequences available in Phytozome. The sequences were aligned with MUSCLE, block selection was applied using BMGE with a BLOSUM30 matrix and a block size of 4, and the tree was built using IQ-TREE. Values from 1,000 ultrafast bootstrap repetitions are indicated at each node. Land plants are shown in dark green, while other green lineages are depicted in light green. The global topology observed here is consistent with the tree in **Fig. 1A**. Note that the distinct protein identifiers utilized for *Arabidopsis* and other species between panels **A** and **B** reflect the source databases; identifiers in **Fig. 1A** are derived from the NCBI dataset, whereas those in **Fig. 1B** correspond to Phytozome accessions.

Phylogenetic analysis that specifically focused on the Chloroplastida reveals a similar organization; the NADKs group into major cyto- and mito-clades, with the cyto-clade further divided into three subclades. While all three CreNADKs cluster within the cyto-clade, land plants possess members of both the mito- and cyto-clades (**Fig. 1B**). A comparison of the different CreNADKs with those of *Arabidopsis* shows that Cyto-subclade 1 includes CreNADK1, which clusters with AtNADK2, and human NADK1. Given that AtNADK2 has been experimentally localized to chloroplasts (Chai et al., 2005), this group likely represents a chloroplast-targeted NADK within photosynthetic eukaryotes. Cyto-subclade 2 includes CreNADK3, which clusters with AtNADK1; the latter is localized to the cytosol. We have placed CreNADK2 into cyto-subclade 3, which is not represented by a direct ortholog in *Arabidopsis*. Additionally, the peroxisomal AtNADK3 groups with human mito-clade NADK2. This NADK has no homolog in *Chlamydomonas.* Overall, our phylogenetic analyses suggest that NADKs have undergone functional divergence and lineage-specific adaptations across different photosynthetic organisms.

Although the NADKs belonging to the mito-clade and the cyto-subclades 1 and 2 are well characterized in model organisms including *Saccharomyces*, humans, and *Arabidopsis*, cyto-subclade 3 proteins, including CreNADK2, appear to be specific to microorganisms and are largely understudied. Many microorganisms that harbor this NADK are unicellular protists or inhabit soil and aquatic environments that are subject to frequent stress or fluctuations in environmental factors, including light, oxygen, and organic carbon availability. Fluctuating conditions often require rapid acclimation of cellular metabolism, raising the possibility that NADKs in cyto-subclade 3 play a dynamic, underexplored role in acclimation.

### Divergence of Chlorophyte-Specific NADK Clades Supports Metabolic Flexibility

The phylogenetic analyses provide insights into the evolutionary relationship of the NADKs and their functional roles in *Chlamydomonas*. NADKs from the two main clades and most of the subclades are present in chlorophytes, including *Chlamydomonas*, *Ostreococcus lucimarinus*, and *Micromonas pusilla* (the latter two are members of the early diverging Prasinophyceae), as well as in the streptophytes, such as the land plant *Arabidopsis*. The clustering of the NADK sequences from both Chlorophyta and Streptophyta indicates a high degree of conservation, suggesting that these NADKs originated from a common ancestor predating the divergence of green algae and land plants. The separation of cyto-subclades 1 and 2, despite their similar taxonomic representation, may reflect subtle sequence variations or gene duplication events that led to protein neofunctionalization and distinct subfamilies within this conserved lineage.

Cyto-subclade 3 forms a clade exclusively composed of NADKs from the core green algae, including *Chlamydomonas*, but is notably absent in *Ostreococcus lucimarinus*, *Micromonas pusilla*, and land plants. This subclade likely represents a lineage of NADKs that evolved specifically within the core Chlorophyta (Chlorophyceae, Trebouxiophyceae, Ulvophyceae, and Chlorodendrophyceae) following divergence of the Prasinophyceae. Other green algae harboring this NADK, including *Dunaliella* and *Chlorella*, are well known for their robust ability to adapt to changing and/or harsh conditions. *Dunaliella salina* can survive in high-salinity environments (Barbosa et al., 2023) whereas *Chromochloris zofingiensis* and *Coccomyxa subellipsoidea* have been shown to be highly adaptable and grow under diverse environmental conditions (Bleisch et al., 2025). These findings raise the possibility that divergence of cyto-subclade 3 NADKs may be associated with the physiological flexibility required to thrive in variable environments, a feature common among these core green algae. The mito-clade includes NADKs solely in *Ostreococcus lucimarinus*, *Micromonas pusilla*, and land plants, with no representation in the core green algae such as *Chlamydomonas*. The clustering of NADKs from basal Prasinophyceae with land plants suggests that this group represents an ancestral NADK variant retained in these lineages and that has either been lost or significantly diverged in the core Chlorophyta.

The fourth *Arabidopsis* NADK, AtNADKc, has putative orthologs in various eukaryotic lineages (**Fig. S1**). Although it performs the same catalytic function as the other NADKs, the activity of this specific protein appears to be associated with particular stress responses and is linked to the oxidative burst and responses of plants to pathogens (Dell’Aglio et al., 2019). This specialization would provide metabolic flexibility by both maintaining NAD^+^/NADP^+^ balance and enabling more targeted responses to oxidative stress, thereby limiting tissue damage during wounding and pathogen attack. It is worth noting that the naming and numbering conventions associated with each of the NADK genes and proteins are not consistent among different organisms.

### Subcellular Localization of NADK Isoforms

To determine the spatial distribution of the three CreNADK proteins, we expressed each isoform in a *Chlamydomonas* mutant strain lacking the corresponding kinase, using a fluorescent reporter construct where mNeonGreen was fused to the C-terminus of the NADK protein. Confocal microscopy revealed distinct localization patterns for each enzyme (**Fig. 2A**). CreNADK1 was exclusively localized within the chloroplast whereas CreNADK3 displayed a fluorescent signal characteristic of cytosolic localization. CreNADK2, however, exhibited a unique dual-targeting profile, with fluorescent signals detected in both the chloroplast and mitochondrion. The mitochondrial association of CreNADK2 can be most clearly observed in cortical views of the cells (**Fig. 2A**, CreNADK2 cortex section image) where the reticulated mitochondrial network is more readily resolved. To further substantiate these observations, full Z-stack galleries of the mNG fluorescent signals were generated and are presented in **Fig. S2**, allowing for comprehensive visualization of 3D protein localization within the cell. To authenticate the observed mitochondrial localization, we utilized the mitochondrion-specific dye MitoTracker Red CMXRos with the cell lines expressing CreNADK2-mNeonGreen fusions. Colocalization analysis confirmed that CreNADK2 aligned with the mitochondrial marker (**Fig. 2B**). These localization patterns partially align with the phylogenetic relationships of NADKs across photosynthetic and non-photosynthetic eukaryotes. CreNADK1 is most similar to the chloroplastic AtNADK2, reflecting conserved plastidial function. Similarly, cytosolic CreNADK3 clusters with the cytosolic AtNADK1. Intriguingly, however, the cytosolic human NADK and the two cytosolic paralogs in *Saccharomyces* share higher sequence similarity with the chloroplastic isoforms in *Chlamydomonas* and *Arabidopsis* than with the cytosolic isoforms in these green lineage organisms. This suggests a complex evolutionary history in which the cytosolic NADKs in plants and algae may have a distinct origin from those in opistokonts.

**Fig. 2.**
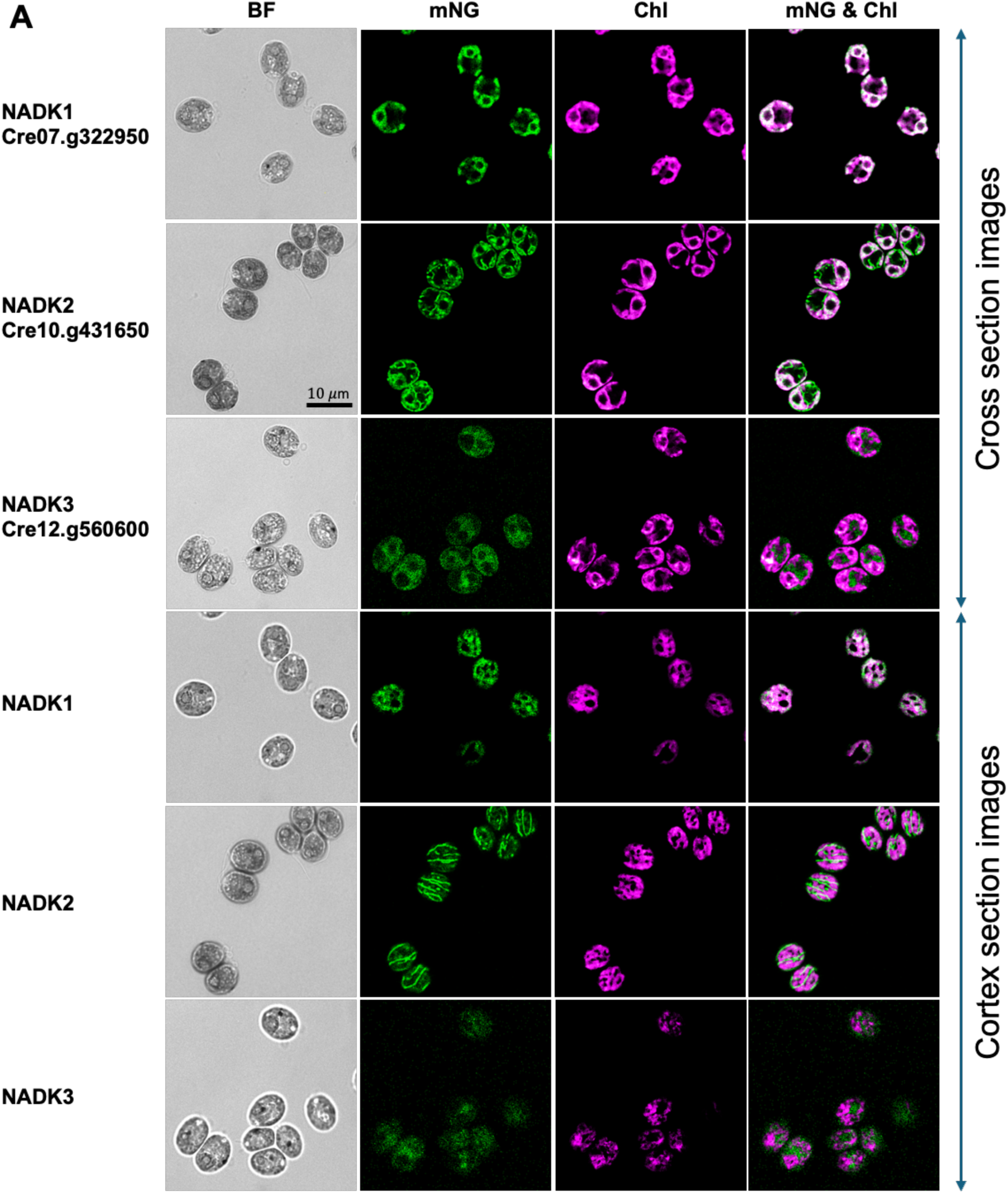
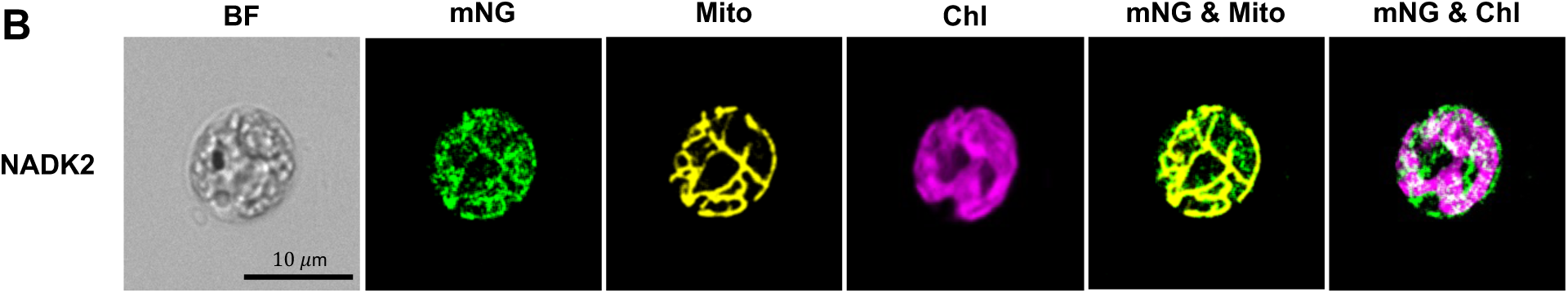
Subcellular localization of CreNADK1, CreNADK2, and CreNADK3. To overcome transgene silencing and ensure robust expression, a bicistronic strategy was implemented using the pRT067 vector which couples the *NADK* transgenes with a paromomycin resistance cassette (see Materials and Methods). All NADK isoforms were expressed with the mNeonGreen (mNG) tag fused to their C-termini. Localization patterns were verified in at least three independent transformants per each genetic background across multiple genetic backgrounds, including wild-type (WT), single *nadk* mutants, and double-mutant combinations. (**A**) Subcellular localization of the NADKs. Representative confocal images of CreNADK1, CreNADK2, and CreNADK3 are presented as longitudinal cross-sections and cortical sections (near cell surface). The cortical sections clearly reveal localization of CreNADK2 in mitochondria (in addition to the chloroplast), a feature not observed in either CreNADK1 or CreNADK3 transformants. Representative images are shown in cross-sections (top three rows) and cortical sections (bottom three rows). From left to right: brightfield (BF), mNG fluorescence indicating NADK localization (mNG), chlorophyll autofluorescence (Chl), and merged mNG and Chl signals (mNG & Chl). In the NADK3 localization panel, brightness was linearly enhanced to facilitate visual detection of the low-intensity signal. (**B**) Colocalization of the CreNADK2-mNG signal with the mitochondria-specific dye MitoTracker Red CMXRos in cross section. Columns from left to right: brightfield (BF), mNG fluorescence showing CreNADK2 protein localization (mNG), MitoTracker Red CMXRos indicating mitochondrial distribution (Mito), chlorophyll autofluorescence (Chl), merged CreNADK2 and mitochondrial signals (mNG & Mito), and merged CreNADK2 and chlorophyll signals (mNG & Chl). Full Z-stack galleries of the mNG fluorescent signals are provided in **Fig. S2**. Scale bar (10 μm) applies to all panels.

### Light-Dependent CreNADK2 Distribution

To our knowledge, no dual-targeted NADK has been previously described. The localization of CreNADK2 to both chloroplasts and mitochondria, principal hubs of energy transduction (processes linked to ROS production), suggests a pivotal role for this enzyme in orchestrating redox homeostasis and metabolic fluxes associated with varying trophic conditions. Additionally, this localization pattern combined with the finding that CreNADK2 resides within a distinct phylogenetic clade separated from the two established cytosolic clades, identifies this enzyme as a unique target for functional studies.

To determine the spatial distribution of CreNADK2 between the mitochondria and chloroplast, we employed a modified, quantitative 3D Z-stack confocal image analysis (Malkovskiy et al., 2025) shown in **Fig. S3**. Partitioning of CreNADK2-mNG was assessed under heterotrophic (dark), mixotrophic, and photoautotrophic conditions, at low (30 μmol photons m^−2^ s^−1^) and moderate (250 μmol photons m^−2^ s^−1^) light intensities. Our results reveal that CreNADK2 localization is highly dynamic and responsive to the energetic state of the cell (**Fig. 3A** and **B**). Under heterotrophic conditions, CreNADK2 is dually localized to the mitochondria and chloroplast, although it exhibits lower abundance in the latter. However, the transition to light triggers a robust redistribution of the enzyme. As irradiance increases, mitochondrial CreNADK2 levels are significantly attenuated, while its abundance in the chloroplast becomes dominant under both mixotrophic and autotrophic conditions (**Fig. 3B**). This shift suggests that while dark-state cells require elevated mitochondrial NAD kinase capacity, the onset of photosynthesis necessitates a redistribution of CreNADK2, with more enzyme associated with the chloroplast to support increased photosynthetic carbon fixation, anabolic metabolism, and detoxification of ROS generated by photosynthetic electron transport. To investigate whether the shifts in CreNADK2 distribution were driven by changes in organelle volume under the different growth conditions, we quantified changes in the volumes of both mitochondria and chloroplasts (**Fig. 3C** and **D**). Chloroplast volume was larger in cells grown in the light compared to those grown in the dark, reaching its maximum under mixotrophic moderate light conditions (**Fig. 3C**). Conversely, total mitochondrial volume was markedly reduced in the light, with the most pronounced reduction observed in photoautotrophic cells compared to cells grown mixotrophically at equivalent light intensities. Under both growth regimes, moderate light further diminished mitochondrial volume relative to low light (**Fig. 3D**). Despite relatively stable total cellular protein content across these conditions (**Fig. 3I**), the density of the CreNADK2 protein within the chloroplast (abundance per unit volume) increased significantly in accord with photosynthetic capacity, peaking in cells grown autotrophically in moderate light (**Fig. 3F**). In contrast, the mitochondrial protein density declined in the light (relative to heterotrophic growth), particularly at moderate intensities (**Fig. 3G**). Notably, in cells grown heterotrophically, mitochondrial protein density reached a maximum, significantly exceeding that of the chloroplast (**Fig. 3F** and **G**). These data indicate that the cellular demand for NAD kinase activity is prioritized toward the mitochondria in the dark and toward the chloroplast under moderate light conditions. Additionally, CreNADK2 exhibits a distinct pattern around the pyrenoid at higher light intensities, suggesting a specialized role for this enzyme in modulating pyrenoid function. Specifically, CreNADK2 may support the carbon-concentrating mechanism (CCM) under elevated photosynthetic rates, where accelerated CO_2_ fixation creates a localized low-CO_2_ environment.

**Fig. 3.**
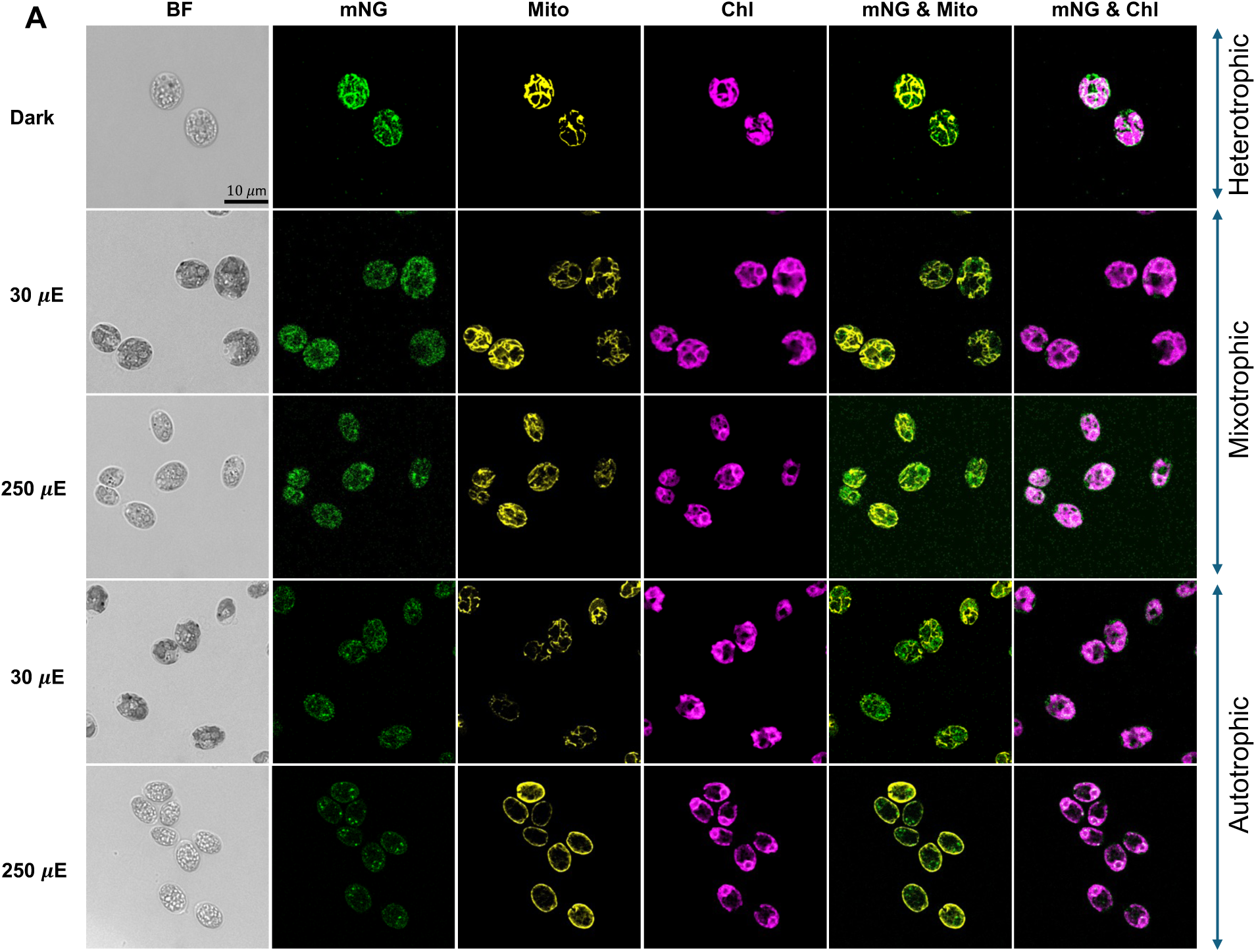
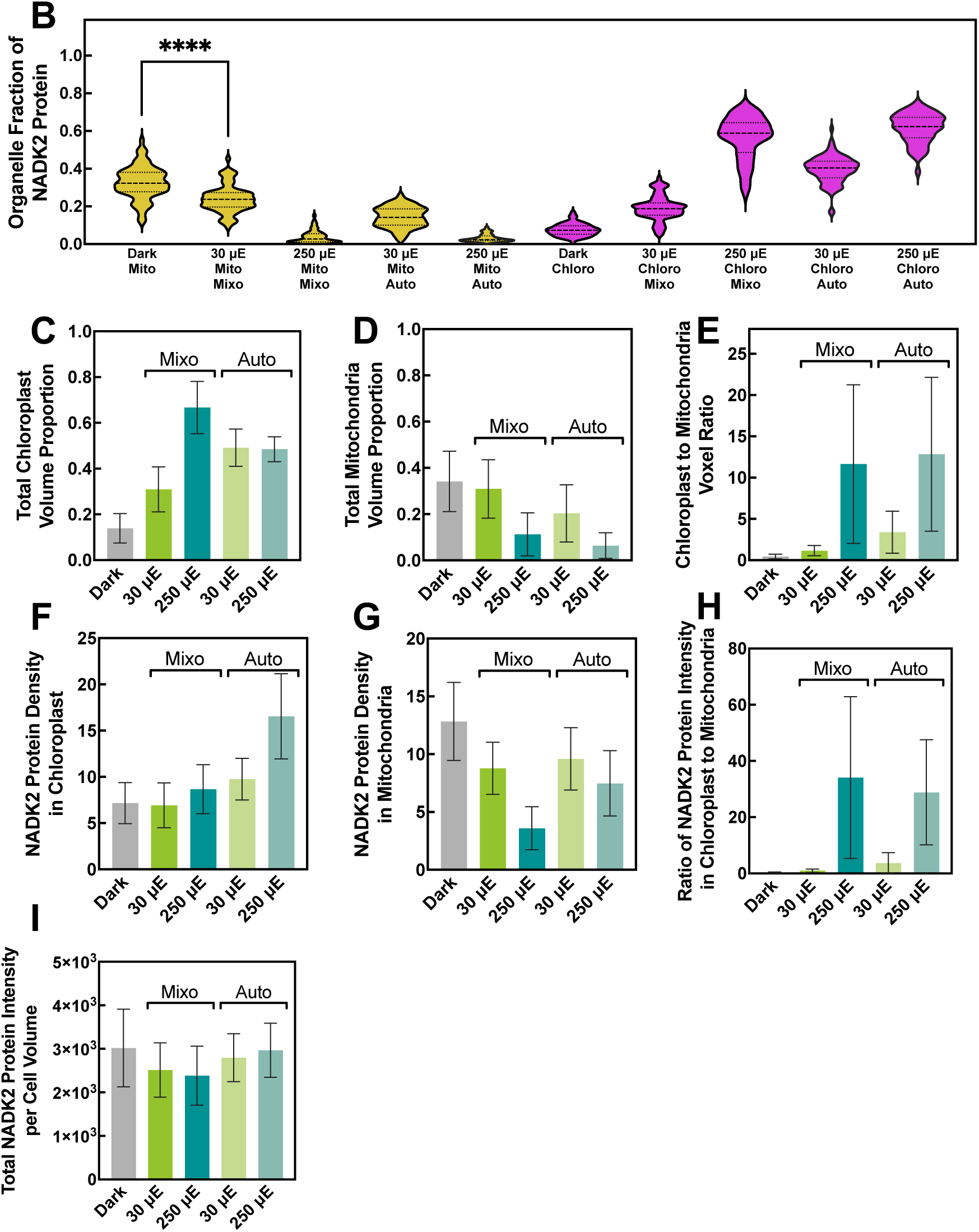
Light-dependent CreNADK2 partitioning between chloroplast and mitochondria. Transformants expressing bicistronic CreNADK2-mNeonGreen (mNG) fusion construct (detailed in **Fig. 2**) were cultured in TAP medium in the dark (heterotrophic) and both in TAP and TP (no acetate) media under low light (30 μE, mixotrophic and photoautotrophic, respectively) and moderate light (250 μE, mixotrophic/autotrophic) to drive different rates of photosynthesis. Subcellular localization was quantified across approximately 100 individual cells per condition. (**A**) Representative confocal micrographs illustrating trophic-dependent redistribution of CreNADK2. Columns from left to right: brightfield (BF), mNG fluorescence indicating CreNADK2 protein (mNG), MitoTracker Red (Mito), chlorophyll autofluorescence (Chl), merged CreNADK2 and mitochondria signals (mNG & Mito), and merged CreNADK2 and chlorophyll signals (mNG & Chl). For conditions in which NADK2 is expressed at low levels, the brightness was linearly enhanced to facilitate visual detection of the low-intensity signal. (**B**) Quantitative partitioning of CreNADK2 between mitochondria (yellow) and chloroplasts (pink), calculated as the fraction of organelle-specific protein intensity relative to total cellular protein intensity. (**C–D**) Volumetric proportions of the chloroplast (**C**) and mitochondria (**D**), represented as total organellar voxels normalized to cell volume in individual cells. (**E**) Ratio of chloroplastic to mitochondrial voxels per cell, indicating relative organellar scaling. (**F–G**) Relative CreNADK2 density within the chloroplast (**F**) and mitochondria (**G**), defined as mNG fluorescence intensity per unit organellar voxel. (**H**) Ratio of chloroplast-to-mitochondria protein intensity. (**I**) Total CreNADK2 abundance expressed as protein intensity normalized to total cell volume. Data represent mean ± SD (≈100 cells). Full Z-stack galleries for all fluorescence channels are provided in **Fig. S4**. The scale bar (10 μm) applies to all panels. μE is equivalent to μmol photons m^−2^ s^−1^. Mito, mitochondria; Chloro, chloroplast; Mixo, mixotrophic; Auto, photoautotrophic.

Our results suggest that CreNADK2 provides the cell with metabolic plasticity, shifting the capacity for NADP(H) production between the cell’s two primary bioenergetic hubs. This analysis further indicates that when the Calvin-Benson-Bassham (CBB) cycle is most active, under moderate light compared to both low light and heterotrophic conditions, both the total fraction and relative density of CreNADK2 in the chloroplast increase, reflecting heightened metabolic demand per unit volume. Finally, we observed a clear correlation between the chloroplast-to-mitochondria volume ratio (**Fig. 3E**) and the corresponding ratio of protein intensities (**Fig. 3H**). This correlation suggests that organellar spatial dominance and surface area availability, where the protein import machinery resides, may impact transport efficiency and steady-state distribution of dual-targeted proteins like CreNADK2.

### CreNADK2 is Essential for Heterotrophic but not Phototrophic Growth

In photosynthetic organisms, NAD kinases generate NADP^+^, a terminal electron acceptor of Photosystem I (PSI). The light-driven generation of NADPH and ATP fuel inorganic carbon assimilation via the CBB Cycle. While vascular plants such as *Arabidopsis*, rice, and tomato typically rely on a single plastid-localized NADK, *Chlamydomonas* encodes two chloroplast-localized isoforms. Cyanobacteria such as *Synechocystis* sp. PCC 6803 also have two NADKs that can modulate the ratio of NAD^+^:NADP^+^ and potentially support different metabolic functions as environmental conditions change (Ishikawa et al., 2016, 2019, 2021). Functional studies of the NADKs in *Arabidopsis* have established that the plastidial AtNADK2 is vital for energy transduction and that At*nadk2* null mutants exhibit severe growth inhibition, chlorosis, and reduced photosynthetic capacity (Chai et al., 2005; Takahashi et al., 2006). In contrast, our autotrophic growth assays of the Cre*nadk2* mutants revealed that the absence of this isoform (which is in a different subclade than the AtNADK2) does not significantly impair growth or chlorophyll accumulation under low (30 μmol photons m^−2^ s^−1^) or moderate (250 μmol photons m^−2^ s^−1^) light intensities (**Fig. 4B** and **C**). Further characterizations of photosynthetic performance based on light-response curves confirmed these observations. Maximum electron transport rates (ETR_max_) in the mutant remained comparable to those of WT cells grown across four light regimes, 30, 50, 100, and 250 μmol photons m^−2^ s^−1^ (**Fig. 4D**). This lack of a photoautotrophic phenotype mirrors observations in *Synechocystis* single *nadk* null mutants; viability during autotrophic growth was not compromised, although metabolite profiles were altered in the *slr0400* strain, which was null for one of the NADKs (Ishikawa et al., 2016). By retaining two plastidial NADKs, the *Chlamydomonas* chloroplast, like cyanobacteria, maintains a functional redundancy/overlap that may safeguard photosynthetic electron flux when the activity of one of the NADKs is compromised. This feature appears to have been lost in the lineage leading to vascular plants.

**Fig. 4.**
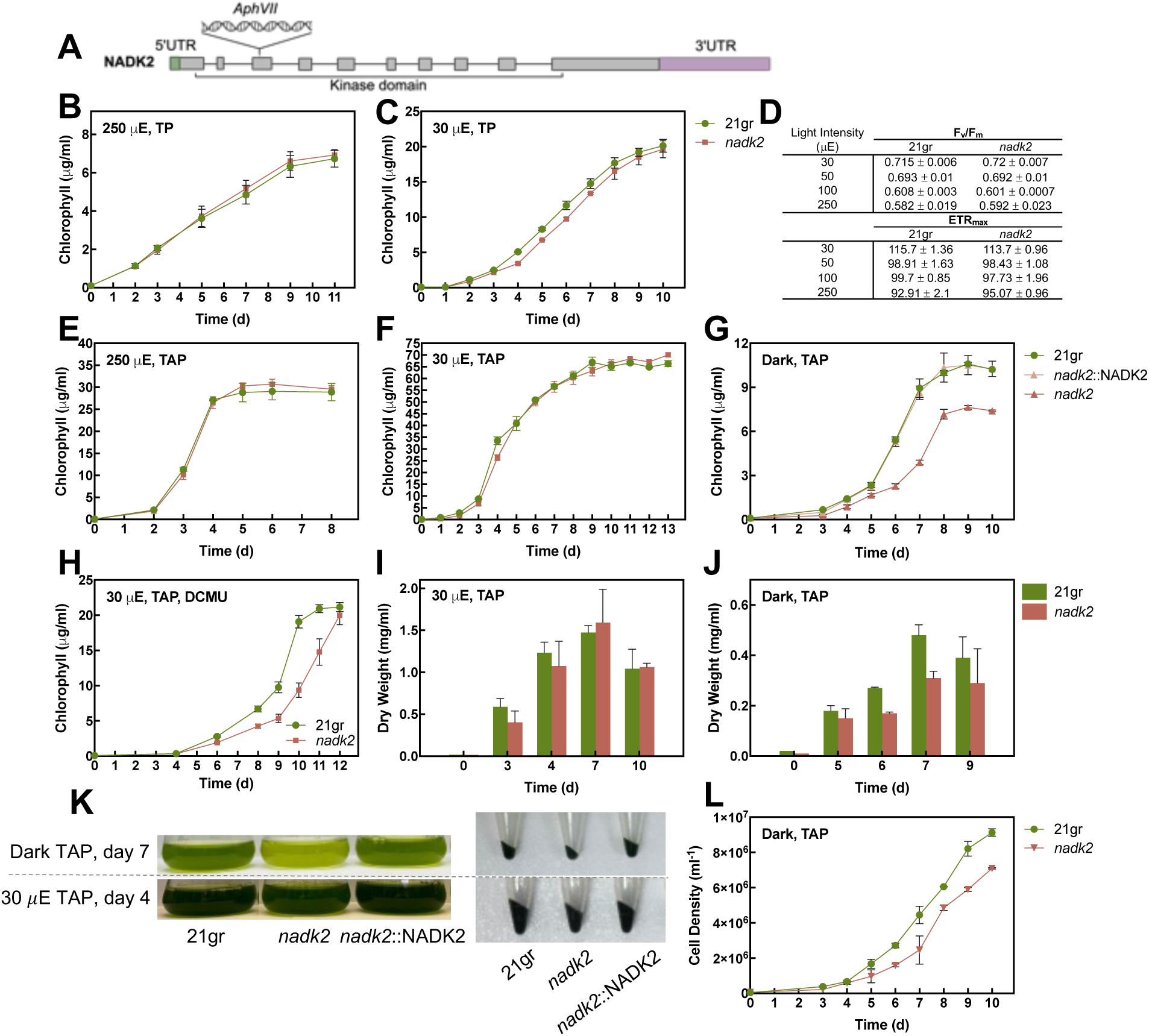
Growth phenotype and photosynthetic performance of Cre*nadk2* mutants. (**A**) Schematic representation of the CRISPR-Cas9-mediated knockout strategy showing insertion of the hygromycin resistance cassette into the third exon of the Cre*NADK2* kinase gene. (**B** and **C**) Photoautotrophic growth of WT and Cre*nadk2* strains based on chlorophyll concentration under moderate light (ML, 250 μE) and low light (LL, 30 μE) conditions. (**D**) Photosynthetic capacity of WT and Cre*nadk2* mutants measured using pulse-amplitude-modulated (PAM) fluorometry. The (F_v_/F_m_) represents the maximum quantum yield of PSII whereas ETRmax is the maximum electron transport rate (ETR_max_) based on light-response curves (**Fig. S5**). (**E** and **F**) Mixotrophic growth (TAP medium) of WT and Cre*nadk2* based on chlorophyll concentration under ML and LL. (**G** and **H**) Heterotrophic growth in the dark and under LL conditions in the presence of 10 μM 3-(3,4-dichlorophenyl)-1,1-dimethylurea (DCMU). (**I** and **J**) Comparative biomass accumulation of WT and Cre*nadk2* as determined by dry weight measurements in LL (mixotrophic) and the dark (heterotrophic). (**K**) Representative images of mid-logarithmic phase cultures and corresponding cell pellets derived from 100 mL cultures grown under mixotrophic (LL) and heterotrophic (dark) conditions. (**L**) Cell densities (cells/mL) of WT and Cre*nadk2* cultures during heterotrophic growth in the dark. Data represent means ± SD (n≥3). μE is equivalent to μmol photons m^−2^ s^-1^.

Mixotrophic growth assays (TAP medium) under low and moderate light intensities revealed no significant differences in chlorophyll accumulation in WT cells compared to Cre*nadk2* mutants (**Fig. 4E** and **F**). However, biomass analysis in low light indicated a transient growth lag in Cre*nadk2* at the onset of the logarithmic phase, although the mutant attained WT dry weight levels once the cells reached late log phase (**Fig. 4I**). The most pronounced phenotype of the mutant was observed under heterotrophic conditions (dark) where acetate serves as both the primary carbon and energy source. Under these conditions, Cre*nadk2* exhibited severe growth impairment based on all methods of normalization; chlorophyll concentration, dry weight, and cell density (**Figs. 4G**, **J**, and **L**).

A similar observation was documented for *Synechocystis* sp. PCC 6803, where individual NADK deletions do not significantly impact mixotrophic growth, and yet the loss of the *sll1415* NADK isoform (but not the *slr0400* isoform) drastically reduced photoheterotrophic growth (Gao & Xu, 2012). To determine if this phenotype in *Chlamydomonas* was strictly dependent on the absence of light or the absence of photosynthetic activity, we assessed growth in medium supplemented with acetate under low light conditions in the presence of DCMU, an inhibitor of Photosystem II. The growth inhibition of the mutant relative to WT cells under this condition confirmed that the loss of this kinase compromises heterotrophic metabolism whether in the dark, or in the light in the presence of DCMU (**Fig. 4H**). The dispensability of CreNADK2 may reflect functional redundancy among isoforms (e.g. CreNADK1 and CreNADK2 in chloroplast) and potential delivery of NADP^+^ synthesized in the chloroplast or cytoplasm to the mitochondria. Aspects of our findings are exemplified by the *Arabidopsis* mitochondrial RNA polymerases (RPOTmp); while RPOTmp is dual localized, its mitochondrial function is partially differentiated from the mitochondria-specific RPOTm (Kühn et al., 2007). This functional specialization suggests why dually targeted proteins are maintained despite the presence of organelle-resident homologs. Conversely, the lack of functional divergence between chloroplastic RPOTp and dually targeted RPOTmp likely renders the latter redundant in the plastid, and therefore it would be only essential for mitochondrial function (Tarasenko et al., 2016). A similar regulatory logic likely governs the *Chlamydomonas* NAD kinase system. The presence of plastid-localized CreNADK1 may provide sufficient activity to mask the loss of CreNADK2, which would be redundant under standard mixotrophic conditions. However, the unique pool of mitochondrial CreNADK2 lacks a redundant counterpart, rendering it critical when the cells are grown heterotrophically and mitochondrial energy transduction is the primary driver of cellular metabolism. Our results suggest that dual targeting of CreNADK2 represents a strategic distribution of NADK activity, ensuring adequate cofactor synthesis in mitochondria under dark conditions, while providing a supplementary boost to the chloroplastic NADP(H) pool in the light when photosynthetic carbon assimilation becomes highly active. Preservation of this enzyme partitioning suggests a regulated mechanism that provides the cells with metabolic resilience. Additionally, by maintaining a baseline presence of CreNADK2 in both bioenergetic compartments, the cell ensures a primed response to environmental variability, a hypothesis supported by dynamic density shifts observed under the different irradiances (**Fig. 3C**). However, the observation that cells can survive the loss of mitochondrial CreNADK2 activity in the dark, albeit with a significantly diminished growth rate, prompted us to investigate potential underlying metabolic compensations. To understand the cellular survival strategy in the absence of this mitochondrial NADP(H) source, we characterized the metabolic responses of these cells under both heterotrophic and mixotrophic conditions.

### Loss of CreNADK2 Triggers Inhibition of TCA Cycle and Metabolic Rerouting

To elucidate the metabolic consequences of CreNADK2 deficiency, we performed targeted LC-MS/MS-based quantification of metabolites associated with central carbon metabolism in cells grown heterotrophically in the dark and mixotrophically in low light. Heterotrophically grown Cre*nadk2* cells accumulated intermediates associated with the TCA cycle, glyoxylate shunt, gluconeogenesis, and both the oxidative and reductive branches of the pentose phosphate pathway (PPP) (**Fig. 5A**). Specifically, citrate, α-ketoglutarate, glyoxylate, malate, oxaloacetate, and multiple glycolytic/gluconeogenic intermediates including phosphoenolpyruvate, pyruvate, dihydroxyacetone phosphate, fructose 1,6-bisphosphate, and glucose 6-phosphate were significantly elevated in the mutant relative to WT cells. This metabolic congestion may be reflected by a mitochondrial oxidative burst stemming from the loss of CreNADK2-mediated NADP^+^ synthesis. In heterotrophic *Chlamydomonas*, mitochondria rely on NADPH-dependent antioxidant systems to scavenge ROS generated by electron leakage at Respiratory Complexes I and III. Specifically, superoxide radicals are converted to hydrogen peroxide by superoxide dismutase, which is subsequently detoxified via the glutathione peroxidase and peroxiredoxin cycles. These pathways require NADPH as the ultimate electron donor for both glutathione reductase and thioredoxin reductase (Dayer et al., 2008). In the Cre*nadk2* mutant, the mitochondrial NADPH pool is likely depleted, as the loss of CreNADK2 activity disrupts the primary biosynthetic route for the synthesis of mitochondrial NADP(H). Furthermore, the reductive capacity of key NADPH-generating enzymes, including malic enzyme, isocitrate dehydrogenase, aldehyde dehydrogenase, and nicotinamide nucleotide transhydrogenase, would be functionally impaired (**Fig. 6**). This deficiency could stem from a dual limitation: a reduced substrate pool (NADP^+^) and a restricted supply of reducing equivalents that results from the metabolic diversion of acetate toward the glyoxylate shunt rather than the TCA cycle (discussed in the next section). Collectively, these perturbations would compromise mitochondrial antioxidant defenses. The resulting oxidative stress would damage ROS-sensitive TCA cycle enzymes; aconitase, which contains a labile [4Fe4S] cluster (Fridavich, 1995; Yan et al., 1997), the α-ketoglutarate dehydrogenase complex, which utilizes a lipoic acid cofactor that is highly susceptible to oxidative modification (Tretter & Adam-Vizi, 2004), and pyruvate dehydrogenase which has cysteine residues that are susceptible to oxidation (Hurd et al., 2012). Oxidative poisoning of these enzymes creates metabolic constrictions that explain the simultaneous accumulation of citrate and α-ketoglutarate in the Cre*nadk2* mutant. Using organelle-targeted roGFP2 redox biosensors, we confirmed that the mitochondrial matrix is significantly more oxidized in the Cre*nadk2* mutant compared to WT cells during dark growth (**Fig. 5D**), providing intracellular spatial confirmation of this redox perturbation.

**Fig. 5.**
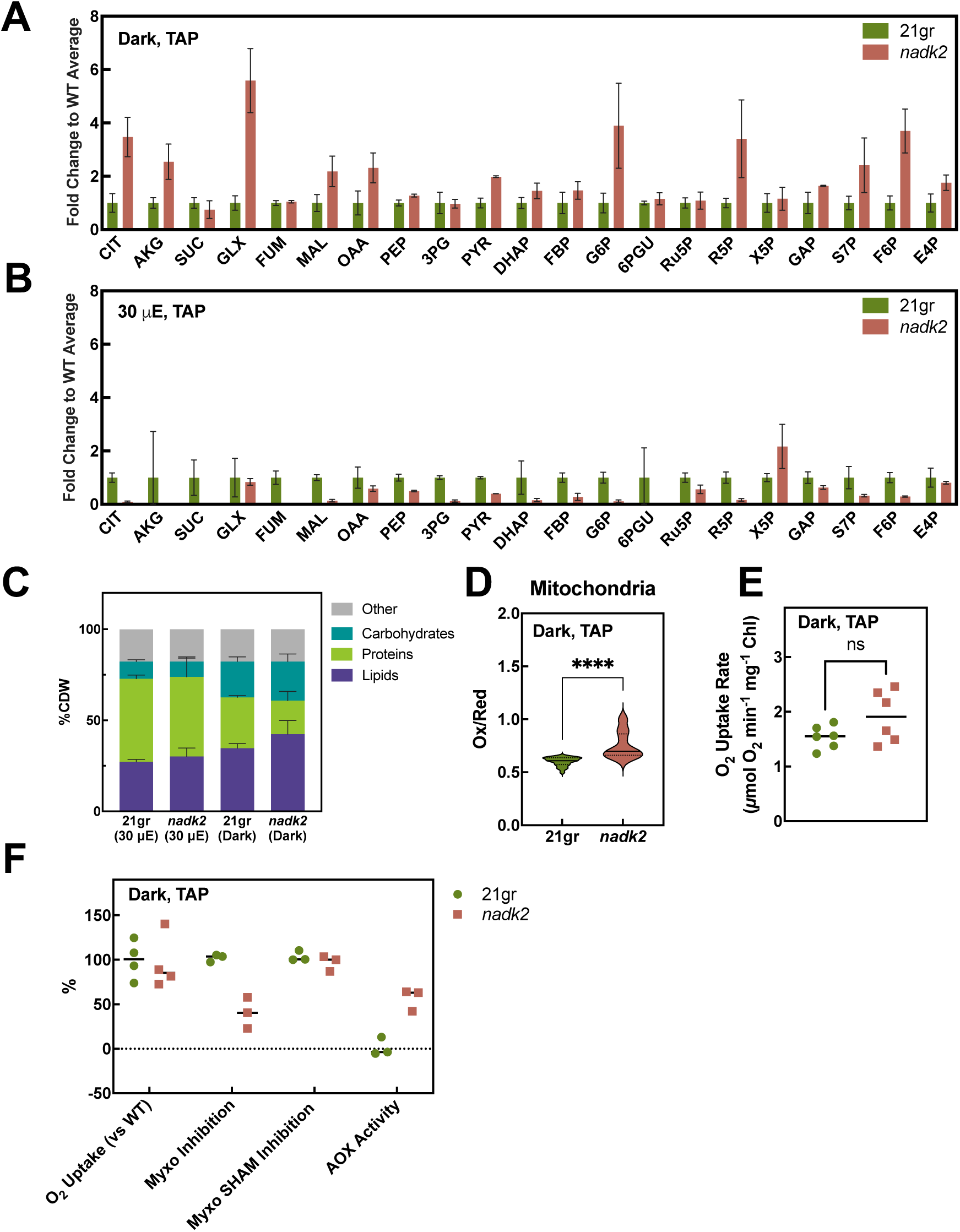
Metabolite and redox profiles of Cre*nadk2* mutants under different trophic conditions. (**A** and **B**) LC-MS-based metabolite profiles from WT (21gr) and Cre*nadk2* cells grown in TAP medium in (**A**) the dark and (**B**) under low light (30 μE) conditions. Values represent the fold change in the Cre*nadk2* mutants relative to the average in WT cells, which was set at 1. (**C**) Total biomass composition including carbohydrate, protein, lipid, and residual components in WT and Cre*nadk2* mutant cells grown in dark and low-light conditions. For all panels, the data represents the mean ± SD (n=3 biological replicates). **(D)** In vivo redox state of WT and Cre*nadk2* cells based on fluorescence from roGFP2 expressed in the cells and targeted to the mitochondrial matrix. Cells were grown in the dark, and the redox state was trapped using N-ethylmaleimide (NEM) prior to ratiometric fluorescence measurements (Gutscher et al., 2008). Data are means ± SD (n=3 independent experiments, ≥24 technical replicates). **(E)** Respiratory O_2_ uptake rates in WT and Cre*nadk2* mutant cells during heterotrophic growth. Data are a means ±SD (n=2 independent biological replicates with 3 technical replicates each, 6 total replicates). **(F)** Respiratory flux partitioning in Cre*nadk2* relative to WT. Oxygen uptake was measured in untreated cells, cells treated with myxothiazol (cytochrome pathway inhibitor), and cells treated with both myxothiazol and SHAM (alternative oxidase inhibitor). The alternative oxidase (AOX) capacity was calculated as the SHAM-sensitive fraction of respiration. Data are means ± SD (n≥3 biological replicates). Statistical significance was determined by Student’s t-test (P values provided for critical comparisons). μE is equivalent to μmol photons m^−2^ s^−1^. Metabolites: 3PG 3-phosphoglycerate; 6PGU, 6-phosphogluconate; AKG, α-ketoglutarate; CIT, citrate; DHAP, dihydroxyacetone phosphate; E4P, erythrose 4-phosphate; F6P, fructose 6-phosphate; FBP, fructose 1,6-bisphosphate; FUM, fumarate; G6P, glucose 6-phosphate; GAP, glyceraldehyde 3-phosphate; GLX, glyoxylate; ICT, isocitrate; MAL, malate; OAA, oxaloacetate; PEP, phosphoenolpyruvate; PYR, pyruvate; R5P, ribose 5-phosphate. Ru5P, ribulose 5-phosphate; S7P, sedoheptulose 7-phosphate; SUC, succinate; X5P, xylulose 5-phosphate.

**Fig. 6.**
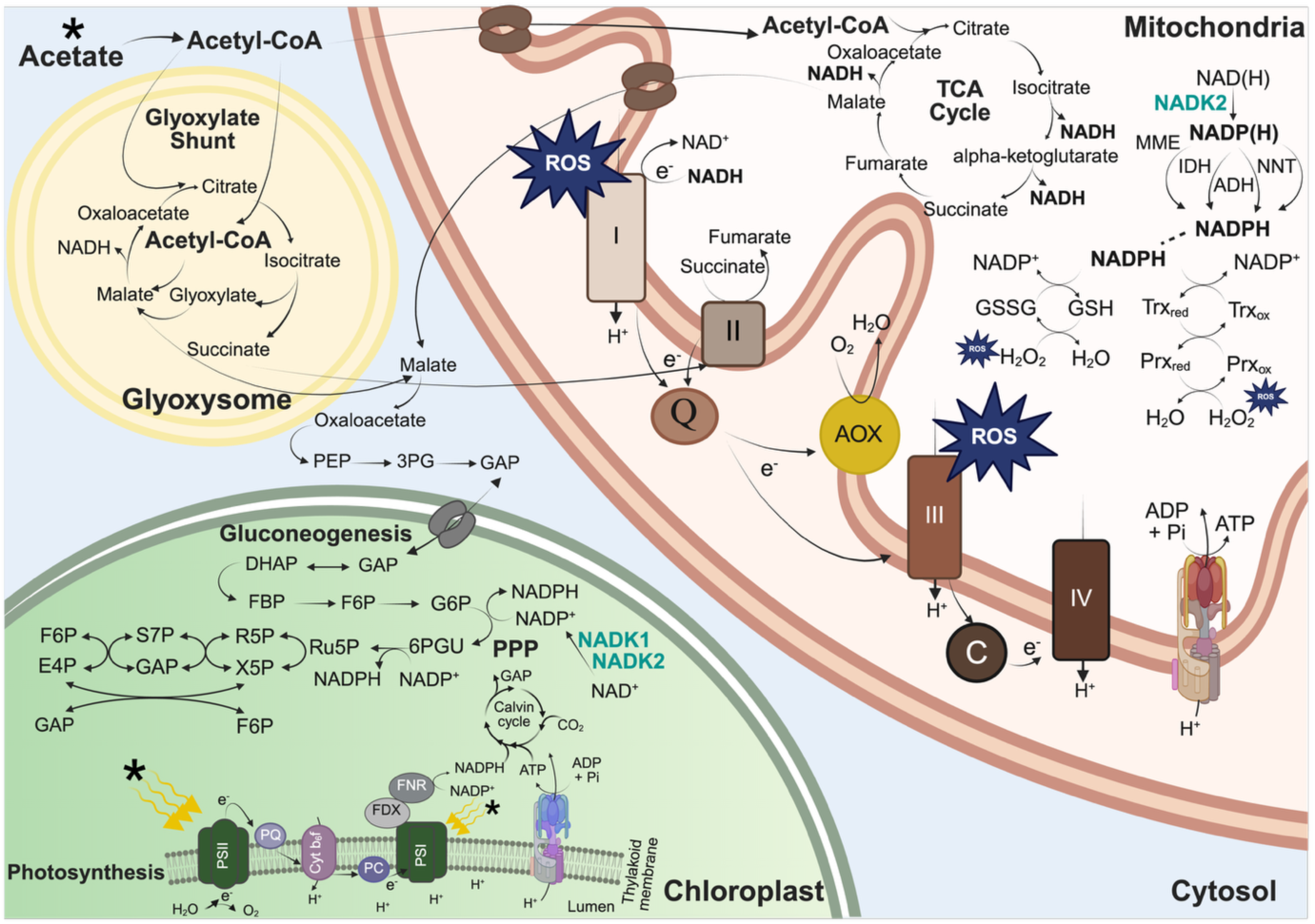
Schematic of integrated metabolic network in *Chlamydomonas* illustrating redox and carbon flux partitioning. The map highlights the primary metabolic intersections between four subcellular compartments. In the mitochondria, the conventional respiratory electron transport chain (Complexes I–IV) and alternative oxidase (AOX) pathways are depicted alongside the tricarboxylic acid (TCA) cycle and the NADPH-dependent ROS detoxification systems. The glyoxysome houses the glyoxylate shunt as a carbon-conserving bypass of the TCA cycle. Shown in the cytosol is the conversion of malate to glyceraldehyde 3-phosphate (GAP), which can provide fuel for gluconeogenesis. The chloroplast integrates photosynthetic electron transport (PET), the CBB cycle, gluconeogenesis, and the PPP. Two primary sources of energy are indicated by asterisks. Metabolites: 3PG, 3-phosphoglycerate; 6PGU, 6-phosphogluconate; ADP, adenosine diphosphate; ATP, adenosine triphosphate; CO_2_, carbon dioxide; DHAP, dihydroxyacetone phosphate; E4P, erythrose 4-phosphate; F1,6P, fructose 1,6-bisphosphate; F6P, fructose 6-phosphate; GAP, glyceraldehyde 3-phosphate; G6P, glucose 6-phosphate; GSH, glutathione; GSSG, glutathione disulfide; H^+^, proton; H_2_O, water; H_2_O_2_, hydrogen peroxide; NAD^+^, nicotinamide adenine dinucleotide; NADH, reduced nicotinamide adenine dinucleotide; NADP^+^, nicotinamide adenine dinucleotide phosphate; NADPH, reduced nicotinamide adenine dinucleotide phosphate; O_2_, molecular oxygen; PEP, phosphoenolpyruvate; Prx_ox_, oxidized peroxiredoxin; Prx_red_, reduced peroxiredoxin; R5P, ribose 5-phosphate; ROS, reactive oxygen species; Ru5P, ribulose 5-phosphate; S7P, sedoheptulose 7-phosphate; Trx_ox_, oxidized thioredoxin; Trx_red_, reduced thioredoxin; X5P, xylulose 5-phosphate. Proteins: ADH, aldehyde dehydrogenase; C, mitochondrial plastoquinone; FDX, ferredoxin; FNR, ferredoxin-NADPH reductase; IDH, isocitrate dehydrogenase; MME, malic enzyme; CreNADK1, NAD kinase 1; CreNADK2, NAD kinase 2; NNT, nicotinamide nucleotide transhydrogenase; PC, lumen plastoquinone. Protein Complexes: AOX, alternative oxidase; Cyt b_6_f, cytochrome b_6_f; I, respiratory complex I; II, respiratory complex II; III, respiratory complex III; IV, respiratory complex IV; PSI, photosystem I; PSII, photosystem II. Pathways: PPP, pentose phosphate pathway; TCA, tricarboxylic acid cycle. PQ, chloroplastic plastoquinone pool; Q, mitochondrial quinone pool. Created with BioRender.com.

### AOX and the Glyoxylate Bypass Enable Survival of Cre*nadk*2 Mutants

Despite a compromised mitochondrial antioxidant capacity, Cre*nadk2* cells remain viable under heterotrophic conditions, indicating the occurrence of compensatory survival strategies. While total oxygen uptake in the dark was similar in the mutants and WT cells (**Fig. 5E**), the respiratory pathway bias in the Cre*nadk2* strain was significantly different. In WT cells, a large majority of respiratory electrons used cytochrome oxidase as the terminal acceptor while in the Cre*nadk2* mutants approximately 50% of total respired electrons were redirected through the alternative oxidase (AOX) pathway (**Fig. 5F**). This pathway bypasses proton-pumping Complexes III (ubiquinol-cytochrome c oxidoreductase) and IV (cytochrome c oxidase), thereby preventing over-reduction of the ubiquinol pool and diminishing superoxide formation and ROS accumulation, albeit at the cost of a reduced ATP yield (Li et al., 2023; Maxwell et al., 1999).

The bioenergetic shift described above is coupled with a strategic redirection of the carbon flux. The accumulation of glyoxylate in mutant cells aligns with the finding that a greater proportion of the organic carbon used by the cell is a consequence of acetate metabolism, which requires the activity of the glyoxylate shunt. This shunt allows the cell to conserve/assimilate organic carbon because it bypasses the decarboxylation steps of the TCA cycle (**Fig. 6**). To quantitatively evaluate the activity of the pathways involved in central carbon metabolism, we performed ^13^C metabolic flux analysis. Our results demonstrate that in Cre*nadk2* cells grown heterotrophically, the flux of acetate into the TCA cycle decreased by ∼82%, whereas the acetate flux supporting the glyoxylate shunt increased by ∼210% relative to WT cells (**Fig. 7A** and **B**). Consistent with this metabolic redirection, the overall flux through TCA cycle intermediates plummeted by 35% in the mutant. Crucially, this bypass diminishes the reducing pressure on Complex I (NADH:ubiquinone oxidoreductase) by eliminating two major NADH-generating reactions. Because Complex I is both a primary source of ROS production and a highly sensitive target for oxidative damage (Murphy, 2008) associated with its labile [Fe-S] clusters, its function is likely compromised in the Cre*nadk2* mutant. This putative oxidative poisoning of Complex I impairs the cell’s ability to oxidize NADH, creating a significant bottleneck in the recycling of reducing equivalents and directly contributing to an elevated mitochondrial NADH/NAD^+^ ratio. By utilizing the glyoxylate bypass, the cell minimizes production of NADH, which can no longer be efficiently processed by Complex I, thereby preventing further electron leakage and ROS generation. The resulting succinate generated by the glyoxylate shunt along with diminished TCA cycle activity likely favors electron input to Complex II (succinate dehydrogenase). As Complex II is structurally insulated against ROS generation and does not contribute directly to the proton motive force, its preferential use lowers the mitochondrial membrane potential. Additionally, since ROS production exponentially changes with respect to the membrane potential, this shift serves as a biophysical shield against further superoxide generation (Ahmed Selim & Wojtovich, 2025; Starkov & Fiskum, 2003). However, these protective measures result in the generation of secondary metabolic backpressure. Oxidative inhibition of Complex I creates a persistent NADH surplus. A high level of NADH acts as a potent allosteric inhibitor of citrate synthase, effectively closing the gate to the TCA cycle (Harford & Weitzman, 1975; Srere & Matsuoka, 1972). The observed accumulation of malate and oxaloacetate in the Cre*nadk2* mutants may be driven by two primary mechanisms. First, inhibition of mitochondrial citrate synthase would restrict metabolite flux into the TCA cycle, resulting in the localized accumulation of these metabolites at the cycle’s entry point. Second, there may be an imbalance between the rate of production of metabolites in the glyoxylate shunt and consumption of these metabolites in downstream pathways, including gluconeogenesis and the oxidative pentose phosphate pathway (OPPP), resulting in production levels that exceed the cell’s assimilatory capacity.

**Fig. 7.**
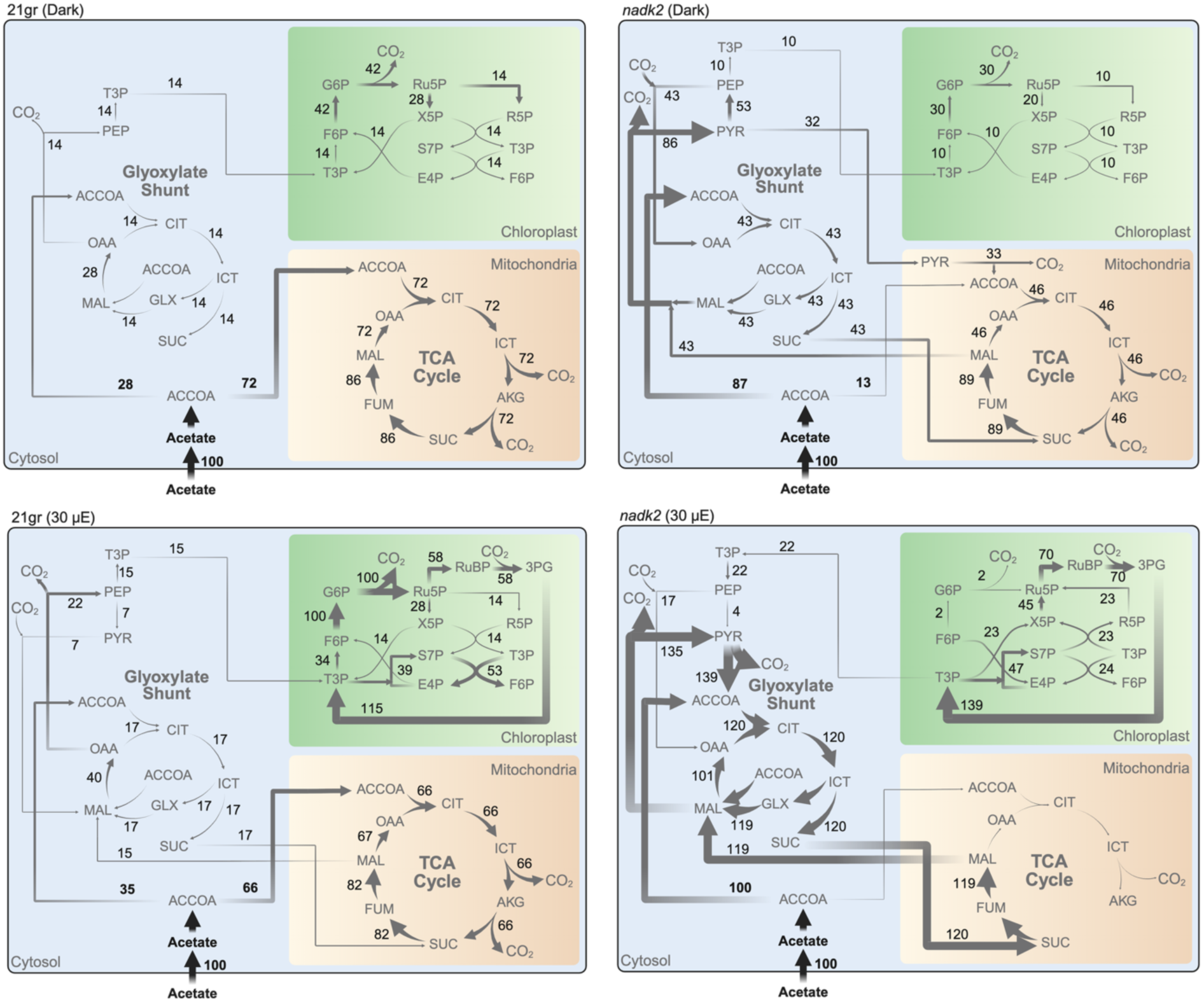
Metabolic flux of central carbon metabolism in WT and Cre*nadk2* mutants. Quantitative carbon flux maps of cells exposed to dark heterotrophic (**A** and **B**) and low light (30 μE) mixotrophic (**C** and **D**) growth conditions. Results were generated via INCA using steady state ^13^C labeling. The metabolic flux analysis focuses on the TCA cycle, glyoxylate shunt, and pentose phosphate pathway. Net flux values (expressed in mmol g^−1^ DW h^−1^) correspond to the optimal value within the 95% flux confidence intervals (**Supplemental Data 1**). All represented fluxes are normalized to a constant acetate uptake flux of 100 mmol g^−1^ DW h^−1^. To minimize visual complexity, relative fluxes with a magnitude of less than 1 mmol g^−1^ DW h^−1^ are omitted from the map, and remaining arrow thicknesses are scaled to represent relative metabolic fluxes. μE is equivalent to μmol photons m^−2^ s^−1^. Metabolites: 3PG, 3-phosphoglycerate; ACCOA, acetyl-CoA; AKG, α-ketoglutarate; CIT, citrate; CO_2_, carbon dioxide; E4P, erythrose 4-phosphate; F6P, fructose 6-phosphate; FUM, fumarate; G6P, glucose 6-phosphate; GLX, glyoxylate; ICT, isocitrate; MAL, malate; OAA, oxaloacetate; PEP, phosphoenolpyruvate; PYR, pyruvate; R5P, ribose 5-phosphate; Ru5P, ribulose 5-phosphate; RuBP, ribulose 1,5-bisphosphate; S7P, sedoheptulose 7-phosphate; SUC, succinate; T3P, triose 3-phosphate; X5P, xylulose 5-phosphate. Created with BioRender.com.

Our data also suggests that the cell maintains fumarate at basal levels, potentially to avoid the toxic effects of fumarate accumulation. At high concentrations, fumarate can cause protein succination, a deleterious non-enzymatic modification of cysteine residues. To mitigate this risk, excess fumarate is converted into malate and oxaloacetate. These metabolites serve as versatile metabolic currencies that can readily be exported to the cytosol to alleviate excess accumulation of metabolites and redox equivalents in the mitochondria and support auxiliary pathways. Once in the cytosol, the high concentration of oxaloacetate serves as the fuel for gluconeogenesis via phosphoenolpyruvate carboxykinase (PEPCK), forcing the redirection of carbon into the OPPP (**Fig. 6**). Additionally, the preservation of carbon skeletons combined with the inability to rapidly metabolize them through a poisoned TCA cycle, necessitates a marked redirection of organic carbon toward storage. Consequently, Cre*nadk2* mutants exhibited elevated carbon storage relative to WT cells; under heterotrophic conditions, total lipid production increased by 22% and total carbohydrate production by 9.5% (**Fig. 5C**). The synthesis of these macromolecules can serve as metabolic safety valves that diminish mitochondrial redox pressure and allow for fixed carbon storage, which can be mobilized when the supply of energy is diminished. However, this survival strategy comes at a high biosynthetic cost: total protein content in dark-grown mutant cells plummeted by 34% (**Fig. 5C**). This drastic reduction suggests a state of suboptimal metabolism that creates a severe energy deficit for the cells because of redirection of electron flow to AOX, which generates low levels of ATP relative to cytochrome oxidase. Hence, to sustain viability, the energetically expensive processes of protein synthesis and assembly appear to be deprioritized.

### Gluconeogenic Overload and OPPP Bottlenecks

The redirection of carbon toward gluconeogenesis in the Cre*nadk2* mutants results in the intracellular accumulation of glucose 6-phosphate. Although the OPPP, present in both the cytosol and chloroplast of *Chlamydomonas*, would be sustained by plastidial and cytosolic NADP^+^ pools (generated by CreNADK1 and CreNADK3, respectively), the reduced production of NADP^+^ in the Cre*nadk2* mutant might cause a kinetic bottleneck in this pathway. The conversion of glucose 6-phosphate to 6-phosphogluconate is the rate-limiting step of the OPPP. As carbon is diverted away from the throttled TCA cycle and into gluconeogenesis, the generation of glucose 6-phosphate exceeds the pathway’s oxidative capacity. This accumulation of glucose 6-phosphate is further exacerbated by the cell’s diminished energy status as mitochondrial electron flow is diverted through the ‘non-phosphorylating’ AOX, which mitigates ROS production and restricts ATP synthesis. The resulting energy deficit would slow downstream anabolic pathways, such as nucleotide and cell wall synthesis, which normally consume pentose intermediates. Consequently, carbon accumulates in the pentose pool (**Fig. 6**). Intriguingly, we observed the specific accumulation of ribose 5-phosphate, while the levels of its isomers, ribulose 5-phosphate and xylulose 5-phosphate, remained low. This suggests a thermodynamic shift within the non-oxidative branch of the PPP. In a state of restricted metabolism that slows consumption of metabolic intermediates (reduced anabolic processes), the equilibrium appears to favor the aldose form (ribose 5-phosphate) over the ketose forms (ribulose 5-phosphate and xylulose 5-phosphate) of the 5 carbon intermediates.

Together, our data suggests that *Chlamydomonas* mitigates mitochondrial oxidative stress when grown heterotrophically in the absence of CreNADK2 by diverting the flux of carbon from catabolic energy production and the synthesis of key metabolites that would be used for growth toward anabolic storage carbon. Such an adjustment in the mutant minimizes mitochondrial ROS generation, allows for maintenance of essential mitochondrial metabolism and baseline ATP production, and stimulates the preservation of fixed carbon in storage forms that can be accessed when needed, at the cost of a diminished rate of heterotrophic growth.

### Photosynthesis-Driven Metabolic Reprogramming Alleviates the Impact of Mitochondrial Redox Stress in Cre*nadk2*

In contrast to the metabolic congestion observed in the dark, the metabolomic profile of Cre*nadk2* mutants grown under low light intensity revealed a distinct, opposite metabolic landscape. Generally, net accumulation of metabolites was significantly lower in low light cultures compared to dark-grown cells (**Table S1**), which likely reflects a higher metabolic flux and metabolite turnover rate in the presence of light. Furthermore, unlike the accumulation observed in the dark, the Cre*nadk2* mutants displayed lower levels of almost all accumulated metabolites compared to WT cells (**Fig. 5B**). This light-dependent, marked shift in the metabolite profile can be attributed to the ‘unloading’ of bioenergetic burdens from the mitochondria to the chloroplast. In the dark, mitochondrial respiration is the sole energy producer, requiring maximum TCA cycle capacity to generate NADH for the synthesis of ATP and NADPH to fuel biosynthetic activities. Mitochondrial activity would likely hit a redox wall in the absence of CreNADK2 and diminished levels of NADP^+^, leading to mitochondrial oxidation and damage to cellular proteins. In low light, however, the burden shifts: the chloroplast would act as the major power plant, providing ATP and NADPH derived from photosynthetic electron transport, with the chloroplast localized CreNADK1 supplying high levels of NADP^+^ which is the primary electron acceptor for Photosystem I.

Our results indicate that the respiratory rate of WT cells declines in low light, TAP-maintained cultures compared to dark maintained cultures (**Fig. S6A**). Following transfer of the cells to the dark, the TCA cycle transitions from a high-output energy generator to a basal-level biosynthetic hub. It transitions to a primary function in the dark of providing α-ketoglutarate for glutamate and purine synthesis, citrate for fatty acid production, succinyl-CoA for chlorophyll synthesis, oxaloacetate for aspartate synthesis and gluconeogenesis, with the flexibility of also providing citrate and oxaloacetate from the glyoxylate shunt. This metabolic demand for carbon skeletons, coupled with low steady-state pools, likely explains the reason for the near-total depletion (undetectable levels) of TCA cycle intermediates, including citrate, α-ketoglutarate, succinate, and fumarate, and the very low levels of malate. Our flux analysis provides physical evidence for this reprogramming, whereas acetate partitioning between the TCA cycle and the glyoxylate shunt was comparable in WT cells under both dark and light conditions. Interestingly, the Cre*nadk2* mutants underwent a near-total metabolic pivot in the light (relative to the dark). In low light, almost all acetate flux in the mutant was channeled into the glyoxylate shunt, with only minimal allocation to the TCA cycle (**Fig. 7A**, **C**, and **D**). This shift confers two significant advantages to Cre*nadk2* mutants. First, it mitigates the risk of ROS generation through diminished TCA cycle activity, which limits NADH production and reduces the respiratory rate. Crucially, this redirection also bypasses the use of the α-ketoglutarate dehydrogenase complex, which can be a significant non-respiratory source of ROS within the mitochondrial matrix (Tretter & Adam-Vizi, 2004). Second, while dark-grown cells rely on the glyoxylate shunt for survival, light-grown cells primarily utilize the CBB cycle for *de novo* carbon fixation. Intriguingly, this suggests that the availability of photosynthetically-derived energy removes the oxidative pressure generated by mitochondrial respiration. By maintaining a reduced but functional TCA cycle to provide essential metabolites, the mutant bypasses the vulnerability of an oxidized mitochondrial matrix. Comparable levels of glyoxylate observed in Cre*nadk2* and WT cells further suggest that the shunting of acetate and the subsequent blockage of gluconeogenesis and the OPPP observed in the dark do not occur when the cells are exposed to light. These physiological adjustments observed in cells grown in low light effectively balance catabolism and anabolism, allowing Cre*nadk2* mutants to maintain growth rates comparable to those of WT cells. The impact of this photosynthetic/light rescue is most evident in the recovery of the cellular macromolecular profile. Under mixotrophic conditions, the severe protein deficit observed in the dark was almost entirely rescued, with only a marginal 4% reduction in the Cre*nadk2* mutants compared to WT cells. Furthermore, the excess carbohydrate that accumulates in the dark was eliminated, showing no significant difference between genotypes in the light. While total lipid production remained elevated in the mutant by 11%, this represents a 50% reduction in accumulation compared to levels observed during heterotrophic growth (**Fig. 5C**). These results demonstrate that light-driven energetics mostly alleviate the mitochondrial metabolic bottleneck, providing the necessary ATP to restore protein homeostasis and normalize carbon partitioning.

### CreNADK2 Deficiency Alters the Amino Acid Composition

Beyond reducing total protein quantity, the loss of CreNADK2 fundamentally altered the qualitative amino acid composition of the cells. These shifts suggest that Cre*nadk2* mutants synthesize or maintain a distinct set of proteins compared to WT cells, with the amino acid profile being strongly modulated by light availability. During heterotrophic dark growth, the amino acid profile of the Cre*nadk2* protein fraction exhibited a marked change (**Fig. S6B**). We observed an increased proportion of alanine, valine, leucine, proline, glutamate, lysine, arginine, and histidine residues within the total protein fraction. Conversely, the relative content of serine, methionine, and tyrosine was significantly reduced. This shift suggests that in the absence of mitochondrial CreNADK2, the cell’s translational landscape is reconfigured, favoring accumulation of proteins richer in the former group of amino acids while limiting those with a high proportion of the latter group. During mixotrophic growth in low light, an even more pronounced change in the amino acid profile was observed relative to WT cells (**Fig. S6C**). While the proportions of glycine, proline, and aspartate increased, the most striking change was a 3- to 4-fold increase in the lysine and histidine content in the biomass protein fraction of mutants relative to WT cells. In contrast, the proportions of isoleucine, serine, threonine, phenylalanine, arginine, and tyrosine were significantly lower in the mutants. Most notably, cysteine levels in the Cre*nadk2* protein fraction dropped below the limit of detection under mixotrophic conditions. These data demonstrate that the loss of CreNADK2 leads to an amino acid profile that is severely depleted of sulfur-containing and aromatic amino acids, especially upon illumination. However, it is important to note that the recovery of cysteine and tryptophan from proteins is incomplete because of the inherent susceptibility of these amino acids to degradation during acid hydrolysis. Therefore, these residues are likely systematically underestimated across all samples, although the relative comparison between mutant and WT cells remains qualitatively robust. The significant reduction of cysteine within the protein fraction, coupled with the robust enrichment of lysine and histidine, reflects a profound, light-dependent shift in the mutant amino acid composition.

### Evolutionary Divergence in Mitochondrial Bioenergetics and ROS Management

The metabolic fate of cells lacking mitochondrial NADK is dictated by the bioenergetic features and respiratory and photosynthetic electron transport chains of the cell. One aspect of divergence between the bioenergetics of *Chlamydomonas/Saccharomyces* (yeast) and mammalian cells is the presence of a mitochondrial AOX, a non-proton-pumping (or poorly proton-pumping) terminal oxidase that occurs in *Chlamydomonas* and yeast but not in mammals. This allows fungal and algal cells to bypass the standard cytochrome oxidase pathway when stressed, creating a safety mechanism that dissipates excess cellular energy that might otherwise cause cell damage. *Chlamydomonas*, however, also faces a unique challenge compared to *Saccharomyces*; unlike yeast, it possesses a classic, proton-pumping Complex I (NADH:ubiquinone oxidoreductase), similar to that of human cells. While efficient for ATP generation, Complex I is a primary site of superoxide leakage, particularly when the ubiquinone pool is over-reduced. Yeast, which lacks Complex I entirely, relies on internal and external NADH dehydrogenases (Ndi1, Nde1/2) that do not pump protons. Consequently, while both yeast and *Chlamydomonas* have the AOX safety valve, *Chlamydomonas* must also manage the ROS liability associated with Complex I. These respiratory differences shape the physiological and metabolic phenotypes of the mitochondrial NAD kinase mutants in mammalian cells, yeast, and *Chlamydomonas*.

The functional rigidity of mammalian mitochondria, specifically the absence of AOX and the reliance on Complex I, might be one of the reasons why there are severe physiological consequences in vertebrates deficient for a mitochondrial NADK. In mice, homozygous *Nadk2* knockouts result in embryonic and preweaning lethality (Dickinson et al., 2016). Recent characterization confirms that results from studies with murine models recapitulate key features of mitochondrial *nadk* mutants in humans, including the occurrence of severe metabolic and neuronal abnormalities (R. Zhang & Zhang, 2023). The pronounced neuronal vulnerability of the mutant reflects its diminished mitochondrial activity, especially given the high energetic demands of maintaining ion gradients and synaptic transmission. Consequently, neuronal cells are the first to fail when mitochondrial NADK is deficient and the redox homeostasis is perturbed. In humans, the severity of pathologies correlates directly with residual enzymatic activity. Patients with hypomorphic mutations (e.g., splice site or alternative start codon usage) in *NADK* genes may survive longer than those in which there is a lesion that causes a complete loss of function, but their bioenergetic capacity remains critically compromised. For instance, a patient with a hypomorphic mutation in *NADK2* who survived for 10 years displayed clear signs of mitochondrial failure. Beyond the expected impact of NADPH deficiency on fatty acid oxidation, respiratory chain assays revealed activities at the very bottom or slightly below the bottom of the normal range, particularly for Complexes I+III and IV. This profile suggests a generalized mitochondrial dysfunction where cells do not produce sufficient ATP to meet their physiological demands, likely contributing to observed neurological impairments such as ataxia and microcephaly (Tort et al., 2016). Conversely, severe loss-of-function mutations invariably caused fatal encephalopathy in early childhood (Houten et al., 2014; Pomerantz et al., 2018). Collectively, these findings raise the possibility that the evolutionary absence of metabolic bypasses, such as those found in algal and fungal systems, renders the mammalian mitochondrial matrix, and specifically the energetically demanding neuronal environment, uniquely vulnerable to redox collapse precipitated by NADPH depletion. Shianna et al. (2006) observed that the *pos5* deletion in yeast triggers downregulation of respiratory complex genes and key antioxidant enzymes such as SOD2 and CCP1. They also attributed the observed oxidative stress primarily to disrupted iron–sulfur cluster assembly and subsequent iron overload, which drives Fenton chemistry and mitochondrial mutagenesis (Shianna et al., 2006). However, viewing these data through the lens of our *Chlamydomonas* findings suggests an alternative, preventative interpretation. We propose that downregulation of respiratory genes observed in both systems is not merely a symptom of failure, but a regulated strategy to minimize mitochondrial ROS generation through diminished metabolic activity.

Although *Chlamydomonas* possesses the ‘high-risk’ ROS generating Complex I, similar to that of humans, it also exhibits the metabolic plasticity of plants allowing for preventative metabolic rerouting which helps preserve redox homeostasis. In heterotrophic conditions, the cell deploys a two-pronged defense to mitigate Complex I liability. First, the glyoxylate shunt acts as a carbon valve, diverting flux away from the NADH-generating reactions of the TCA cycle toward the production of succinate. Succinate can enter the respiratory chain via Complex II (FADH_2_), a pathway that bypasses the high-risk electron leakage site of Complex I. Second, this altered flux is coupled to AOX activity, which bypasses Complexes III and IV, and serves as a safety valve that does not generate a proton gradient and prevents over-reduction of the ubiquinone pool. Thus, while human cells hit a redox wall due to the rigidity of their electron transport system, *Chlamydomonas* survives by reconfiguring the flow of both electrons and fixed carbon to diminish the inherent risks associated with Complex I.

## CONCLUDING REMARKS

*Chlamydomonas* has three NAD kinases, with CreNADK1 specific to the chloroplast and CreNADK3 specific to the cytoplasm. In contrast, CreNADK2 represents a member of a previously uncharacterized clade of NAD kinases that is dually targeted to both the mitochondrion and chloroplast, the primary bioenergetic organelles of *Chlamydomonas*. Importantly, the sub-cellular distribution of CreNADK2 is governed by trophic conditions and the energy metabolism associated with those conditions. Under heterotrophic conditions (dark + acetate), CreNADK2 partitions to both organelles, with its level in the chloroplast relatively low. Upon exposure to light, this kinase undergoes a robust spatial redistribution; as irradiance levels increase, mitochondria-localized CreNADK2 abundance is markedly attenuated, becoming more abundant in the chloroplast where it supports both mixotrophic and autotrophic growth. This dynamic situation suggests that the distribution of CreNADK2 reflects the energetic ‘work-load’ of the two organelles under the different growth conditions.

The physiological significance of the spatial dynamics of CreNADK2 is reflected by the phenotypes of Cre*nadk2* mutants. The mutants are severely impaired for growth under heterotrophic conditions where acetate is the sole carbon source and most of the cell’s energy is derived from mitochondrial respiration, with much less NADPH likely available for detoxification of respiratory-generated ROS. However, the growth phenotype is rescued under phototrophic conditions, where photosynthesis provides the necessary energetic landscape to compensate for diminished involvement of mitochondria in energy metabolism and circumvents the oxidative crisis triggered by a depleted mitochondrial NADPH pool. Dark-grown Cre*nadk2* mutants employ a two-pronged amelioration strategy for their survival: first, they reroute acetate flux through the glyoxylate shunt rather than the TCA cycle and later toward carbon storage (polysaccharides and lipids), thereby bypassing two NAD-reducing reactions of the TCA cycle and decreasing electron pressure on respiratory complex I, which generates ROS; second, they direct electrons toward AOX, preventing over-reduction of the ubiquinol pool and minimizing superoxide production. These findings reveal sophisticated mechanisms that govern organellar NAD metabolism, enabling microbial eukaryotes to maintain redox homeostasis and thrive in fluctuating environmental landscapes.

## MATERIALS AND METHODS

### Culture Conditions

The WT *Chlamydomonas reinhardtii* strain 21gr (mt^+^) (parental strain) was used for generating knockout mutants. All *Chlamydomonas* strains were grown photo-mixotrophically from stock plates in 50-ml flasks and then diluted into 125-ml flasks in Tris-Acetate-Phosphate (TAP) medium (Harris et al., 2009) and grown at 23 °C under constant illumination (80 µmol photons m^−2^ s^−1^) provided by LED panels. Cells were collected during exponential growth. All mutants were generated using CRISPR/Cas9 editing. Experiments were predominantly performed with cells grown in TAP medium; in some cases, cultures were grown in photoautotrophic Tris-phosphate (TP) medium at 23 °C (indicated in text). All growth experiments were initiated at a chlorophyll concentration of 0.1 µg ml^-1^ which corresponds to a concentration of ∼5-7 × 10^4^ cells mL^−1^. Low light (LL) and moderate light (ML) conditions were 30 ± 2 and 250 ± 10 μmol photons m^−2^ s^−1^, respectively.

### Mutant Generation and CRISPR/Cas9 Mutagenesis

WT *Chlamydomonas* strain 21gr (mt+) was used for generating knockout single mutants of Cre*NADK2* (Cre10.g431650). As detailed in the previous section, WT cells were cultured under continuous illumination at 70-80 μmol photons m^−2^ s^−1^ until the cells attained a density of 3 to 5 × 10^6^ cells mL^−1^. These cells were concentrated to a density of 2 × 10^8^ cells mL^−1^ in TAP medium supplemented with 40 mM sucrose. A single-guide RNA (sgRNA) complementary to the target sequence was designed using CHOPCHOP (https://chopchop.cbu.uib.no/) and synthesized by Integrated DNA Technologies (IDT). The sequence of the sgRNA and its target site is provided in **Supplemental Data 2**.

The protocol for inactivating the *NADK* genes was adapted from (Findinier et al., 2019). Prior to electroporation, Cas9 protein (IDT) and sgRNA were incubated together at 37 °C for 30 min. Approximately 500 ng of PCR product containing the *AphVII* cassette, which confers resistance to hygromycin, was added to the RNP (ribonucleoprotein) mixture. A 250-μL mixture that included cells, the RNP, and the cassette, was electroporated using the GenePulser II Electroporator (Bio-Rad). After a 16-h recovery period in TAP medium supplemented with 40 mM sucrose under very dim light (∼5-6 μmol photons m^−2^ s^−1^), the cells were plated onto solid TAP medium containing 20 μg mL^−1^ hygromycin. The orientation of the *AphVII* insertions, either sense or antisense, was determined by PCR amplification using primer pairs that anneal to the targeted gene on both sides of the inserted locus (**Supplemental Data 2**). The amplified fragments were sequenced to verify the insertion sites (ELIM BIOPHARM, Hayward, USA).

### Plasmid Construction for Mutant Rescue and Determination of Protein Subcellular Localization

The pRT067-PsaD=CrmNeonGreen_V2-3FLAG=APHVIII plasmid (**Fig. S7**) includes the codon-optimized *mNeonGreen* gene and the *AphVIII* ORF for paromomycin resistance. pRT067 was designed to express both a transgene and the resistance gene as a bicistronic mRNA, which enhances the success rate of expressing challenging genes. This plasmid was used as the backbone to individually express *CreNADK1*, Cre*NADK2*, and *CreNADK3* fused to *mNeonGreen*. Genomic DNAs encoding CreNADK1, CreNADK2, and CreNADK3 proteins were cloned into the pRT067 backbone plasmid using Gibson assembly (Gibson et al., 2009) and the individual plasmids were used to complement *nadk1*, *nadk2* and *nadk3* mutants to determine the subcellular localization of the three NADK proteins. To transform these plasmids into *nadk* mutant strains, they were linearized and approximately 700 ng of the linearized plasmids was transformed into the respective mutant backgrounds at a cell density of 2 × 10⁸ cells mL⁻¹. The GeneArt MAX Efficiency Transformation Reagent for algae (Invitrogen) was utilized to introduce the plasmid into algal cells through electroporation using the GenePulser II Electroporator (Bio-Rad) following the manufacturer’s instructions. Putative transformants were selected on solid TAP medium containing 10 µg mL⁻¹ paromomycin. Approximately 1000 of these potential transformants were transferred to individual wells of a 96-well, flat-bottom microtiter plate (Greiner Bio-One 655101), with each well containing 100 μL of TAP medium. These cells were then grown for 3 d while shaking at 200 rpm under 30 μmol photons m^−2^ s^−1^ of cool-white fluorescent light and then diluted twice into the same type of plate with wells containing 100 μL of TAP and 5 μg mL⁻¹ paromomycin and grown for an additional day. These cultures were then screened for mNeonGreen expression using a fluorescent microplate reader (Infinite M1000, TECAN). Excitation and emission settings for mNeonGreen were as follows: excitation at 504 nm with a bandwidth of 12 nm, and emission at 530 nm with a bandwidth of 12 nm. Chlorophyll excitation was performed at 440 nm with a bandwidth of 9 nm, and emission at 680 nm with a bandwidth of 20 nm. For each gene, we selected at least 100 independent transformants that showed the highest mNeonGreen:chlorophyll fluorescence ratio and grew them to a concentration of 2–4 × 10⁶ cells mL⁻¹.

### Confocal Microscopy

Protein localization was visualized using a Leica TCS SP8 confocal laser-scanning microscope equipped with LAS X software. For mitochondrial staining, cells were incubated with 1 μM MitoTracker Red CMXRos (Thermo Fisher Scientific) for 15-30 min prior to imaging. Samples were maintained at room temperature and imaged using a 63X1.4 numerical aperture (NA) oil-immersion objective. Excitation was provided by a white-light laser (WLL) at 505 nm for mNeonGreen and 580 nm for MitoTracker Red CMXRos. To maximize detection specificity and minimize crosstalk, signals were acquired sequentially. Emission was collected using the following windows: 515–570 nm for mNeonGreen (HyD SMD hybrid detector), 590–660 nm for MitoTracker Red (PMT detector), and 680–720 nm for chlorophyll autofluorescence (HyD SMD hybrid detector).

To further isolate signals from the fast-decaying chlorophyll autofluorescence (average lifetime ∼0.1 ns), time-gating was applied: a 2–6 ns gate was used to image mNeonGreen fluorescence, and a 0.3–6 ns gate was used to image chlorophyll fluorescence. Z-stacks consisting of 20–30 slices at 0.3-m intervals were obtained for each field of view. The images presented are projections of 4–6 longitudinal slices. Data were collected from a minimum of 10 independent biological replicates per strain and image analysis, including the generation of maximum intensity projections, was performed using Fiji software (Schindelin et al., 2012).

### Cell Segmentation and Volume Determination

Individual cells were segmented and the images extracted using an improved version of our previously described procedure (Malkovskiy et al., 2025). The mitochondrial and protein content per cell was defined as the relative volume fraction occupied by each signal. This was calculated by multiplying the number of voxels exceeding the background noise threshold by a factor of 2.5 to account for anisotropic Z-axis scaling and dividing by the total cell volume (V). Total cell volume was determined using an ellipsoid fit:

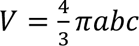

Where a, b, and c represent the semi-axes of the fitted ellipsoid.

### Colocalization Analysis

Colocalization of fluorescent signals was performed via direct pairwise comparisons of voxels across three channels (chlorophyll, mitochondria, and mNeonGreen-protein) for each individual cell. Due to the diffraction-limited resolution of confocal imaging, the signal from a point object spreads according to the system’s Point Spread Function (PSF). The full width at half maximum (FWHM) of this PSF is 0.6λ/NA (≈ 300 nm in our system), which is approximately three-fold greater than our lateral (XY) resolution. To account for the diffraction-limited point PSF and mitigate signal blooming beyond the nominal XY resolution (0.12 μm), a fixed threshold of 100 was applied to the 8-bit signal intensity for the chlorophyll autofluorescence channel. This threshold was empirically determined to capture most of the chloroplast volume while strictly constraining signal spread into adjacent voxels, thereby minimizing false-positive colocalization artifacts. To exclude non-specific background from deteriorating or less healthy cells, a dynamic threshold was applied to the mitochondrial (PMT) signal. This threshold was calculated as a linear function of the average voxel intensity for each individual cell. This robust approach was validated across multiple cell populations; cells exhibiting excessively high background levels were discarded from the final analysis. The spatial partitioning of CreNADK2-mNG was then quantified by calculating the volumetric overlap between the protein signal and the masked organellar channels (mitochondria and chloroplast). Representative 3D reconstructions and the corresponding voxel-based overlap distributions across heterotrophic and phototrophic conditions are detailed in **Fig. S8**.

### roGFP2 Redox Analysis

A roGFP2 sequence, originally developed by Gutscher et al. (2008) and utilized by Staudacher et al. (2018) (Gutscher et al., 2008; Staudacher et al., 2018), was codon-optimized for expression in *Chlamydomonas* and converted to a B5 MoClo part (Crozet et al., 2018). To ensure mitochondrial targeting and robust nuclear expression, the sequence was fused to the *Chlamydomonas* HSP70C transit peptide and interrupted by β-tubulin introns. Expression of this reporter was driven by the AR1 (*HSP70A*-*RBCS2*) promoter and the *PSAD* terminator. For transgene selection, a spectinomycin resistance cassette, under the transcriptional control of the β-tubulin promoter and a *PSAD* terminator, was integrated into the vector. The annotated sequence of the construct is presented in **Supplemental Data 3**. The finalized construct was linearized and transformed into both WT and Cre*nadk2* strains. Primary transformants were selected on TAP agar plates supplemented with spectinomycin. Surviving colonies were initially screened for fluorescence using a microplate reader (Infinite M1000, TECAN) with excitation at 488 nm (bandwidth 9 nm) and emission at 525 nm (bandwidth 10 nm). Subsequently, a TCS SP8 confocal laser-scanning microscope (Leica) was used to validate both the homogeneity of the fluorescence signal across the population and the accuracy of the mitochondrial subcellular localization. Ratiometric analysis of roGFP2 was performed using a microplate reader following the general principles described by Gutscher et al. (2008). To preserve the intracellular redox state during measurements, cells were treated with the membrane-permeable alkylating agent N-ethylmaleimide (NEM) to block reduced thiols. The degree of roGFP2 oxidation was determined by calculating the ratio of emission signals after excitation at 407 nm and 488 nm.

### Phylogenetic Analysis

NADK or NADKc proteins from *Chlamydomonas* and/or *Arabidopsis thaliana* were used as BLAST queries against sequences obtained from several databases (e.g., MMETSP, NCBI, JGI) without applying any filters or clade preference. The first 2,000 BLAST hits obtained with an e-value lower than 1e-10 were selected and aligned using MAFFT (Katoh & Standley, 2013). A preliminary phylogenetic tree was generated with FastTree (Price et al., 2010) after applying block selection with BMGE (Criscuolo & Gribaldo, 2010) using a block size of 3 and the BLOSUM30 similarity matrix. A dereplication step using TreeTrimmer (Maruyama et al., 2013) was applied, which reduced redundancy by sizing down monophyletic clades. The resulting set of sequences was realigned using MUSCLE (Edgar, 2004), and block selection was performed with BMGE using a block size of 4 and the BLOSUM30 matrix. The phylogenetic tree was built with IQ-TREE and the ultrafast bootstrap method (Nguyen et al., 2015). Several rounds of this procedure were conducted to progressively reduce diversity and obtain the final sequence set. For the final phylogenetic analysis, sequences were aligned with MUSCLE and block selection with BMGE was performed as described above. The final phylogenetic tree was constructed using IQ-TREE with the LG4X matrix and 100 bootstrap replicates.

### Dry Weight

A revised version of the protocol described by Mortensen and Gislerod (2014) was used to measure dry weight (Mortensen & Gislerød, 2014). Specifically, 10-15 mL of algal culture, depending on the culture concentration, was filtered through a 0.7-μm pore size, 7 cm-diameter Glass Microfiber Filter Paper (Grade 151, Ahlstrom) to collect the biomass. Filters containing the algae were washed with 20 mL of distilled water to remove adhering salts and then dried overnight at 104 °C. After drying, the filters were cooled to room temperature for 30 min in a desiccator before being weighed. The dry weight was calculated by subtracting the dry weight of the clean filter from the dry weight of the filter with the algae and then was normalized to the volume of the sample collected.

### Chlorophyll Concentration

*Chlamydomonas* cells were harvested by centrifugation at 5,000 × g for 5 min in 1.7-mL tubes, and chlorophyll concentrations measured following methanol extraction as described by (Porra et al., 1989).

### Chlorophyll Fluorescence

Chlorophyll fluorescence was measured using a PAM fluorometer (Dual-PAM 100, Walz GmbH, Germany) equipped with the red light measuring head to determine the electron transport rate through photosystem II. Blue detection pulses (10 µmol photons m⁻² s⁻¹) were applied at 20 Hz to record fluorescence (F). Red saturating flashes (5233 µmol photons m⁻² s⁻¹, 300 ms, 620 nm) were used to determine the maximal fluorescence yield in dark-acclimated samples (Fm) and in actinic light exposed samples (Fm′). The steady-state fluorescence yield measured under actinic light immediately prior to each saturating flash was recorded as Fs. Fluorescence emission was detected through a long-pass filter (>700 nm). The effective quantum yield of photosystem II [Y(II)] was calculated as (Fm′ – Fs)/Fm′. The electron transport rate (ETR) was determined as Y(II) × I × a, where I is the actinic light intensity and a is the correction factor for non-absorbed incident light (default instrument value: 0.84). Samples containing ∼5–8 µg chlorophyll ml⁻¹ were continuously stirred with a magnetic stirrer during measurements. Prior to recording the fluorescence values, the cells were dark-adapted for 10 min. Light-response curves were generated by increasing actinic light intensity stepwise at 1-min intervals. To prevent carbon limitation, bicarbonate was added to the sample at a final concentration of 5 mM.

### Membrane Inlet Mass Spectrometry (MIMS)

Cells were grown heterotrophically in TAP medium in the dark and harvested during exponential growth phase by centrifugation (450 × g, 3 min). Pellets were resuspended in 1.5 mL of fresh TAP medium to a final chlorophyll concentration of 10 μg mL^−1^. Gas exchange rates were monitored using a MIMS reaction chamber equipped with a Teflon membrane. To ensure inorganic carbon saturation, sodium bicarbonate (NaHCO_3_) was added to the suspension at a final concentration of 10 mM. Net O_2_ uptake was recorded in the dark to quantify the respiratory flux. The relative contributions of the respiratory pathways were determined based on the sequential addition of the respiratory inhibitors myxothiazol (2.5 μM) and salicylhydroxamic acid (SHAM; 400 μM); the former selectively inhibit the cytochrome bc1 complex (von Jagow et al., 1984) and the latter inhibits the alternative oxidase (AOX) (Schonbaum,’ et al., 1971).

### LC-MS/MS Analysis

All chemicals used for metabolite extraction and LC-MS/MS analysis were Optima grade reagents, and analytical grade reference standards for each metabolite were obtained from Sigma-Aldrich. Metabolite extraction was performed using the modified procedure of (Sake et al., 2020), incorporating grinding in a cryogenic ball mill to ensure complete cell lysis. Quenched cell pellets were ground with 5 mm diameter stainless steel balls in a Retch cryomill with liquid nitrogen cooling for 2.5 min at 25 Hz. Following grinding, 500 µL methanol was added to the samples along with internal standards, ribitol, and PIPES to a final concentration of 2 µM each. Samples were then subjected to three freeze-thaw cycles by plunging them into liquid nitrogen and thawing them at 1 °C, with vortexing. They were then centrifuged at 10,000 × g at 1 °C for 5 min. The supernatant was collected, and the remaining pellets were extracted two additional times using 500 µL of a 50:50 methanol–water mixture. Supernatants from each extraction step were pooled and dried using a Thermo Fisher SpeedVac concentrator. To remove residual cell debris, dried extracts were resuspended in 500 µL water and filtered sequentially through a 0.22 µm pore size Spin-X centrifuge tube filter, and a filter with a 10 K molecular weight cutoff. Each filter was rinsed with 50 µL water, which was collected and added to the total sample volume for subsequent steps. Filtered extracts were dried once more and resuspended in 100 µL water for LC-MS/MS analysis.

Metabolite extracts were analyzed by LC-MS/MS, following a method adapted from Young et al. (2011) (Young et al., 2011). Chromatographic separation was achieved using a Phenomenex 150 mm × 2 mm Synergi Hydro-RP column on an Agilent 1200 Series HPLC system, coupled to an AB Sciex 5500 QTRAP system. LC was performed with a 20 µL injection volume and gradient elution with 10 mM tributylamine and 15 mM acetic acid (aqueous phase, A) and acetonitrile (organic phase, B), at a constant flow rate of 0.3 mL min^-1^ and column temperature of 40 °C. The organic phase gradient profile was as follows: 0% B (0 min), 8% B (10 min), 16% B (15 min), 30% B (16.5 min), 30% B (19 min), 90% B (21.5 min), 90% B (26.5 min), 0% B (26.6 min), and 0% B (30.5 min). MS analysis was conducted in negative mode using multiple reaction monitoring (MRM) acquisition. Data were acquired using ABSciex Analyst 1.7 software. LC-MS/MS data files (.wiff and .wiff.scan) were converted to .mzML format using OpenChrom (Wenig & Odermatt, 2010). Peak detection, integration, window sizing, and noise assessment were performed using a combination of pyOpenMS (Röst et al., 2014) and SciPy (Virtanen et al., 2020) packages in Python.

### GC-MS Analysis of Amino Acids

For amino acid analysis, the residual pellet from metabolite extraction was hydrolyzed in 6 M HCl under vacuum for 20 h. The resulting amino acids were derivatized to their tert-butyldimethylsilyl forms and analyzed by GC-MS using an Agilent 6890 GC equipped with a DB-1701 column connected to an Agilent 5973 single quadrupole mass spectrometer. GC method parameters followed the description in (Antoniewicz et al., 2007). Data files were converted to .mzML format using OpenChrom (Wenig & Odermatt, 2010) and peak integration was performed using pyOpenMS (Röst et al., 2014) and SciPy (Virtanen et al., 2020) packages in Python.

### Biomass Composition Analysis

For determination of macromolecular biomass composition, harvested cell pellets were pulverized using a Retsch cryomill equipped with 5 mm stainless steel beads under cryogenic conditions to ensure exhaustive cellular disruption. Total lipid content was quantified gravimetrically following a chloroform:methanol extraction of the homogenized biomass, as previously described by Breuer et al. (2013) (Breuer et al., 2013). For quantification of proteins and carbohydrates, aliquots of the pulverized cell lysate were resuspended and diluted in ultrapure water (Milli-Q). Total soluble protein content was measured using the Pierce BCA Protein Assay Kit (Thermo Fisher Scientific) according to the manufacturer’s instructions. Total carbohydrate content was determined colorimetrically using the anthrone-sulfuric acid assay. All biomass components were normalized to dry cell weight. The percent dry weight of nucleotides, pigments, and all other components was calculated using biomass equations presented in the *Chlamydomonas reinhardtii* genome-scale metabolic model iCre1355 (Imam et al., 2015).

### Acetic Acid Measurement and External Flux Analysis

Acetic acid in the spent medium was quantitatively measured using a Cedex Bio Analyzer (Roche). The analysis employed the Acetate V2 Bio Roche kit, an enzymatic photometric assay based on phosphorylation by acetate kinase. Prior to measurement, 1 ml of sample was filtered through a 0.2 µm filter. For external flux analysis, acetic acid concentrations (Caa) and dry weight (DW) at two time points during the logarithmic growth phase were used, and the acetic acid uptake flux was calculated as (Caa₁ − Caa₂) / (DW₂ − DW₁).

### Fluxomics

#### Cultivation and Isotope Labeling

Cultures were grown in a mixture of unlabeled (U-^12^C), partially labeled (1-^13^C), and uniformly labeled (U-^13^C) acetic acid in a 0.4 (U-^12^C): 0.3 (1-^13^C): 0.3 (U-^13^C) ratio. This labeling mixture was selected based on previous metabolic flux analyses performed with *Chlamydomonas* (Boyle et al., 2017). The selection was validated for estimating fluxes in the network model through tracer simulations performed using INCA (Young, 2014). Initial seed flasks with the labeled acetate mixture were grown for each experiment to ensure isotopic steady state for each experiment. These flasks were then used to inoculate experimental cultures grown on the same label.

#### Central Metabolic Network Construction

A compartmentalized central metabolic network model was constructed based on enzyme localization data presented in the literature (Johnson & Alric, 2013). Atom transitions were assigned for each reaction (Mu et al., 2007). Linear reaction pathways were collapsed to ease the computational load, and biomass sink reactions were added. For measured metabolites existing in more than one compartment in the model, an artificial sampling sink reaction was added to account for contributions from these metabolites in different compartments towards the final labeling state. A full list of reactions and associated atom transitions can be found in **Table S2**.

#### Sample Collection and Quenching

Culture samples were collected in mid-exponential growth phase and rapidly quenched to stop metabolic activity and preserve in vivo metabolite concentrations and isotopic labeling patterns. The quenching procedure was performed as described by (Sake et al., 2020). 20 mL aliquots of normal saline (0.9%wt) in 50 mL conical vials were pre-chilled in a −27 °C freezer for approximately 45 min to supercool the solution to −4 °C. Once cooled, 10 mL of the culture samples were rapidly cooled in 10 mL aliquotes by plunging them into the cold saline solution; the quenched mixture was centrifuged at 2°C and 3,000 × g for 5 min. After centrifugation, 28 mL of supernatant was removed and discarded, and the remaining 2 mL of liquid was transferred to microcentrifuge tubes. The remaining supernatant was removed from the samples by centrifugation again at 2 °C and 3,000 × g for 3 min and the quenched cell pellets were stored at −28 °C until analysis.

#### Amino Acids Sample Preparation

For analysis of isotopic labeling of amino acids, proteins were hydrolyzed in 6 M HCl under a vacuum for 20 h and the resulting mixture of amino acids was converted to their tert-butyldimethylsilyl derivatives as described in work by (Antoniewicz et al., 2007). After derivatization, the mixture was analyzed by GC-MS.

#### Organic Acid Sample Preparation

Organic acids were extracted from quenched cell pellets prior to analysis. Metabolite extraction was performed using the extraction procedure of Young et al. (2011) but modified to incorporate grinding in a cryogenic ball mill to lyse the thick cell walls of *Chlamydomonas* (Young et al., 2011). Quenched cell pellets were ground with 5 mm diameter stainless steel balls in a Retch cryomill under liquid nitrogen cooling for 2.5 min at 25 Hz. After grinding, 500 μL of Optima grade methanol was added, and samples were resuspended by pipetting them up and down before transferring them to a fresh tube. Samples were then subjected to three freeze-thaw cycles consisting of dipping the samples into liquid nitrogen and thawing them on a shaker plate at 4 °C before collecting the supernatant after centrifugation at 10,000 × g at 1 °C for 5 min. The supernatant was transferred to a fresh tube, and the remaining pellet was extracted 2X more, as before, using a 50:50 mixture of Optima water and methanol. Supernatants from each extraction were pooled and dried on a Thermo Fisher SpeedVac Concentrator. Dried samples were derivatized and analyzed via GC-MS (Young et al., 2014).

#### Carbohydrates Sample Preparation

Cell wall sugars, synthesized in the cytosol, and starch, synthesized in the plastid, were analyzed to gain compartment-specific information concerning glucose levels in the cytosol and plastid. To analyze cell wall sugars, biomass was hydrolyzed with HCl as described by (McConnell & Antoniewicz, 2016). Cellular starch was hydrolyzed as previously described (Young et al., 2014). Dried samples containing sugar monomers from the above procedures were derivatized with hydroxylamine hydrochloride (2% w/v) in pyridine and propionic anhydride (Young et al., 2014). After derivatization, the mixture was analyzed via GC-MS.

#### Fatty Acids Analysis

To obtain compartment-specific labeling information for acetyl-CoA in the chloroplast, fatty acids were analyzed by GC-MS. Fatty acids from quenched cell pellets were converted to their fatty acid methyl ester (FAME) derivatives and analyzed as previously described (Christie, 1998).

#### GC-MS Analysis

All work was performed on an Agilent 6890 GC using helium as the carrier gas and equipped with an Agilent 5973 single quadrupole mass spectrometer detector. Amino acids, organic acids, and sugars were analyzed on an Agilent DB-1701 column, and fatty acids were analyzed on an Agilent DB-WAX column. GC-MS operating conditions were as described in the literature for amino acids (Antoniewicz et al., 2007), organic acids, sugars (Young et al., 2014), and fatty acids (Christie, 1998). Peak integration was performed using a python script developed in-house using pyOpenMS (Röst et al., 2014). Isotopomer peak areas were corrected for natural isotope abundance in unlabeled atoms using IsoCor v2 (Millard et al., 2019).

#### Flux Modeling

The computational platform INCA (Young, 2014) was used to model intracellular metabolite fluxes for each experiment under steady-state conditions. Experimentally determined acetate uptake rates and mass isotopomer distributions (MIDs) were integrated into the central metabolic network model to calculate internal flux distributions. The INCA software estimates fluxes by minimizing the sum-of-squared residuals (SSR) between simulated and experimental MIDs. A data fit is considered accepted if the total SSR falls within the expected SSR range, which is calculated assuming that the minimized variance-weighted SSR follows a chi-square distribution with *n* − *p* degrees of freedom, where *n* is the number of independent measurements and *p* is the number of fitted parameters. The expected SSR range is defined as 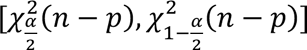, where *α* represents the threshold p-value for fit rejection.

Simulations were performed with a default value of *α* = 0.001. To ensure that results reflected a global SSR minimum, flux estimations were repeated 1,000 times from randomized initial guesses for each condition. Total SSR values and corresponding expected ranges for all simulations are provided in **Supplementary Data 2.**

#### Accession numbers

Genes characterized in this study can be found on https://phytozome-next.jgi.doe.gov/ under loci designations Cre07.g322950 (Cre*NADK1*), Cre10.g431650 (Cre*NADK2*), and Cre12.g560600 (Cre*NADK3*).

## Acknowledgement

We thank Dr. Ryutaro Tokutsu for providing the mNeonGreen_V2 sequence, Dr. Bijie Ren for synthesizing the codon-optimized roGFP2 gene, and Dr. Sai Kiran Madireddi for his invaluable technical expertise and advice regarding photosynthetic characterization.

## Author Contributions

Conceptualization, N.F. and A.G.; Investigation and Data Curation, N.F.; Methodology and Formal Analysis, N.F., M.M., J.F., U.C., A.M., and D.T.; Writing – Original Draft, N.F.; Writing – Review & Editing, A.G., N.F., N.B., and A.B.; Funding Acquisition, A.G. and N.F.; Resources, A.G., N.B., and M.O.; Supervision, A.G. and N.B.. N.F. designed the research and performed the majority of the experiments. M.M. performed the LC-MS/MS metabolite profiling and metabolic flux analysis. J.F. generated the mitochondria-targeting roGFP2 construct and assisted with time-sensitive experiments, including metabolomics sampling. U.C. conducted the phylogenetic analysis. A.M. performed the mitochondrial and chloroplast NADK2 protein quantification. D.T. conducted the respiration measurements. M.O. provided the bicistronic construct.

