## Supplemental Table 1 for "Dual-targeted NADK2 Links Mitochondrial Redox Homeostasis to Carbon Partitioning and Heterotrophic Growth in *Chlamydomonas reinhardtii*"

**Table S1.** **LC-MS-based metabolite profiling of wild type and *nadk2* cells.**

Metabolite concentrations in wild-type (21gr) and *nadk2* mutant cells grown in TAP medium under darkness and low light (LL, 30 μmol photons m^−2^ s^−1^). Samples were harvested during the logarithmic phase of growth for each condition. Values are reported as concentrations (μg mg^-1^CDW). Data represent the mean ± SD of three biological replicates (n=3).

|  | **Dark** | | | | | | **Low Light (30 μmol photons m^−2^ s^−1^)** | | | | | |
| --- | --- | --- | --- | --- | --- | --- | --- | --- | --- | --- | --- | --- |
|  | 21gr 1 | 21gr 2 | 21gr 3 | nadk2 1 | nadk2 2 | nadk2 3 | 21gr 1 | 21gr 2 | 21gr 3 | nadk2 1 | nadk2 2 | nadk2 3 |
| **CIT** | 0.320 | 0.619 | 0.399 | 1.419 | 1.923 | 1.306 | 0.112 | 0.094 | 0.135 | 0.015 | 0.009 | 0.007 |
| **AKG** | 0.016 | 0.011 | 0.013 | 0.024 | 0.039 | 0.039 | 0.000 | 0.001 | 0.000 | 0.000 | 0.000 | 0.000 |
| **SUC** | 0.102 | 0.069 | 0.095 | 0.048 | 0.101 | 0.051 | 0.005 | 0.001 | 0.006 | 0.000 | 0.000 | 0.000 |
| **GLX** | 0.044 | 0.064 | 0.076 | 0.285 | 0.316 | 0.424 | 0.004 | 0.004 | 0.000 | 0.002 | 0.003 | 0.002 |
| **FUM** | 0.095 | 0.097 | 0.081 | 0.093 | 0.100 | 0.093 | 0.009 | 0.005 | 0.009 | 0.000 | 0.000 | 0.000 |
| **MAL** | 0.061 | 0.038 | 0.074 | 0.095 | 0.161 | 0.123 | 0.012 | 0.014 | 0.014 | 0.002 | 0.001 | 0.002 |
| **OAA** | 0.117 | 0.047 | 0.128 | 0.164 | 0.269 | 0.243 | 0.001 | 0.001 | 0.001 | 0.000 | 0.001 | 0.001 |
| **PEP** | 0.019 | 0.016 | 0.020 | 0.023 | 0.023 | 0.024 | 0.001 | 0.001 | 0.001 | 0.000 | 0.000 | 0.000 |
| **3PG** | 0.276 | 0.671 | 0.571 | 0.396 | 0.543 | 0.539 | 0.051 | 0.051 | 0.045 | 0.008 | 0.005 | 0.005 |
| **PYR** | 0.127 | 0.088 | 0.115 | 0.216 | 0.222 | 0.218 | 0.005 | 0.005 | 0.005 | 0.002 | 0.002 | 0.002 |
| **DHAP** | 0.011 | 0.007 | 0.010 | 0.010 | 0.015 | 0.015 | 0.000 | 0.000 | 0.001 | 0.000 | 0.000 | 0.000 |
| **FBP** | 4.893 | 9.683 | 5.234 | 7.212 | 10.695 | 11.179 | 0.629 | 0.750 | 0.892 | 0.325 | 0.168 | 0.134 |
| **G6P** | 0.386 | 0.260 | 0.185 | 1.578 | 0.925 | 0.735 | 0.069 | 0.084 | 0.104 | 0.014 | 0.005 | 0.010 |
| **6PGU** | 0.249 | 0.283 | 0.251 | 0.234 | 0.338 | 0.332 | 0.001 | 0.005 | 0.001 | 0.000 | 0.000 | 0.000 |
| **Ru5P** | 0.030 | 0.023 | 0.021 | 0.036 | 0.026 | 0.020 | 0.001 | 0.001 | 0.001 | 0.001 | 0.001 | 0.001 |
| **R5P** | 0.009 | 0.010 | 0.007 | 0.043 | 0.024 | 0.020 | 0.001 | 0.001 | 0.002 | 0.000 | 0.000 | 0.000 |
| **X5P** | 0.035 | 0.020 | 0.042 | 0.022 | 0.047 | 0.045 | 0.001 | 0.001 | 0.001 | 0.001 | 0.002 | 0.002 |
| **GAP** | 0.040 | 0.018 | 0.042 | 0.054 | 0.055 | 0.055 | 0.001 | 0.001 | 0.002 | 0.001 | 0.001 | 0.001 |
| **S7P** | 0.013 | 0.016 | 0.009 | 0.016 | 0.041 | 0.035 | 0.001 | 0.001 | 0.001 | 0.000 | 0.000 | 0.000 |
| **F6P** | 0.640 | 0.774 | 0.450 | 2.713 | 2.462 | 1.728 | 0.088 | 0.089 | 0.121 | 0.031 | 0.027 | 0.029 |
| **E4P** | 0.014 | 0.007 | 0.012 | 0.018 | 0.023 | 0.018 | 0.000 | 0.000 | 0.000 | 0.000 | 0.000 | 0.000 |
