## Supplemental Table 2 for "Dual-targeted NADK2 Links Mitochondrial Redox Homeostasis to Carbon Partitioning and Heterotrophic Growth in *Chlamydomonas reinhardtii*"

**Table S2. Complete list of metabolic reactions and carbon atom transitions utilized for flux modeling.** The table details the stoichiometric equations and carbon-mapping for all modeled pathways. Compartmentalization of metabolites is indicated by suffixes: .e (extracellular medium), .c (cytosol), .m (mitochondria), and .h (chloroplast). **Metabolites:** 3PG, 3-phosphoglycerate; ACALD, acetaldehyde; ACCOA, acetyl-CoA; Ace, acetate; AKG, alpha-ketoglutarate; Ala, alanine; Ara, arabinose; Asp, aspartate; chor, chorismate; CIT, citrate; CO_2_, carbon dioxide; E4P, erythrose 4-phosphate; F6P, fructose 6-phosphate; FUM, fumarate; G6P, glucose 6-phosphate; Gal, galactose; Glu, glutamate; GLX, glyoxylate; Gly, glycine; ICT, isocitrate; Leu, leucine; MAL, malate; OAA, oxaloacetate; PEP, phosphoenolpyruvate; Phe, phenylalanine; PYR, pyruvate; R5P, ribose 5-phosphate; Ru5P, ribulose 5-phosphate; RuBP, ribulose 1,5-bisphosphate; S7P, sedoheptulose 7-phosphate; FA, fatty acid; Ser, serine; Sta, starch; SUCC, succinate; T3P, triose 3-phosphate; thf, tetrahydrofolate; Thr, threonine; Tyr, tyrosine; Val, valine; Xu5P, xylulose 5-phosphate.

| Number | Reaction | Compartment |
| --- | --- | --- |
| 1 | Ace.e (ab) -> Ace.c (ab) | Transport |
| 2 | Ace.c (ab) -> ACCoA.c (ab) | Cytosol |
| 3 | ACCoA.c (ab) <-> ACCoA.m (ab) | Transport |
| 4 | ACCoA.c (ab) + OAA.c (cdef) -> CIT.c (caebfd) | Cytosol |
| 5 | CIT.c (abcdef) -> ICIT.c (afcbed) | Cytosol |
| 6 | ICIT.c (abcdef) -> GLX.c (df) + SUCC.c (baec) | Cytosol |
| 7 | ACCoA.c (ab) + GLX.c (cd) -> MAL.c (acbd) | Cytosol |
| 8 | MAL.c (abcd) <-> OAA.c (abcd) | Cytosol |
| 9 | SUCC.c (abcd) <-> SUCC.m (abcd) | Transport |
| 10 | MAL.c (abcd) <-> MAL.m (abcd) | Transport |
| 11 | ACCoA.m (ab) + OAA.m (cdef) -> CIT.m (caebfd) | Mitochondria |
| 12 | CIT.m (abcdef) -> ICIT.m (afcbed) | Mitochondria |
| 13 | ICIT.m (abcdef) -> AKG.m (badcf) + CO_2_.m (e) | Mitochondria |
| 14 | AKG.m (abcde) -> SUCC.m (badc) + CO_2_.m (e) | Mitochondria |
| 15 | SUCC.m (abcd) -> FUM.m (abcd) | Mitochondria |
| 16 | FUM.m (abcd) -> MAL.m (badc) | Mitochondria |
| 17 | MAL.m (abcd) <-> OAA.m (abcd) | Mitochondria |
| 18 | MAL.c (abcd) <-> PYR.c (abd) + CO_2_.c (c) | Cytosol |
| 19 | MAL.h (abcd) <-> PYR.h (abd) + CO_2_.h (c) | Plastid |
| 20 | PYR.m (abc) -> ACCoA.m (ab) + CO_2_.m (c) | Mitochondria |
| 21 | PYR.c (abc) -> ACCoA.c (ab) + CO_2_.c (c) | Cytosol |
| 22 | PEP.c (abc) -> PYR.c (abc) | Cytosol |
| 23 | PEP.c (abc) + CO_2_.c (d) <-> OAA.c (abdc) | Cytosol |
|  | PYR.c (abc) <-> PYR.m (abc) | Transport |
| 24 | PYR.h (abc) -> ACCoA.h (ab) + CO_2_.h (c) | Plastid |
| 25 | MAL.c (abcd) <-> MAL.h (abcd) | Transport |
| 26 | MAL.h (abcd) <-> OAA.h (abcd) | Plastid |
| 27 | G6P.c (abcdef) <-> G6P.h (abcdef) | Transport |
| 28 | G6P.h (abcdef) -> Ru5P.h (eadbc) + CO_2_.h (f) | Plastid |
| 29 | Ru5P.h (abcde) <-> X5P.h (abcde) | Plastid |
| 30 | Ru5P.h (abcde) <-> R5P.h (eadbc) | Plastid |
| 31 | S7P.h (abcdefg) + T3P.h (hij) <-> R5P.h (bdfge) + X5P.h (aicjh) | Plastid |
| 32 | S7P.h (abcdefg) + T3P.h (hij) <-> E4P.h (gbfd) + F6P.h (iajhec) | Plastid |
| 33 | F6P.h (abcdef) + T3P.h (ghi) <-> E4P.h (eadc) + X5P.h (bhfig) | Plastid |
| 34 | Ru5P.h (abcde) -> RuBP.h (badce) | Plastid |
| 35 | RuBP.h (abcde) + CO_2_.h (f) -> 3PG.h (ace) + 3PG.h (bdf) | Plastid |
| 36 | T3P.h (abc) + E4P.h (defg) <-> S7P.h (becgafd) | Plastid |
| 37 | G6P.h (abcdef) <-> F6P.h (afbcde) | Plastid |
| 38 | F6P.h (abcdef) <-> T3P.h (ebf) + T3P.h (dac) | Plastid |
| 39 | T3P.h (abc) <-> 3PG.h (bca) | Plastid |
| 40 | T3P.h (abc) <-> T3P.c (abc) | Transport |
| 41 | T3P.c (abc) <-> PEP.c (bca) | Cytosol |
| 42 | AKG.m (abcde) -> Glu.b (abcde) | Mitochondria |
| 43 | PYR.m (abc) -> Ala.b (abc) | Mitochondria |
| 44 | PYR.c (abc) -> Ala.b (abc) | Cytosol |
| 45 | OAA.m (abcd) <-> Asp.m (abcd) | Mitochondria |
| 46 | OAA.c (abcd) <-> Asp.c (abcd) | Cytosol |
| 47 | OAA.h (abcd) <-> Asp.h (abcd) | Plastid |
| 48 | Asp.m (abcd) -> Asp.b (abcd) | Mitochondria |
| 49 | Asp.c (abcd) -> Asp.b (abcd) | Cytosol |
| 50 | Asp.h (abcd) -> Asp.b (abcd) | Plastid |
| 51 | Asp.h (abcd) -> Thr.b (abcd) | Plastid |
| 52 | 3PG.h (abc) -> Ser.h (abc) | Plastid |
| 53 | Ser.h (abc) -> Gly.h (bc) + thf.h (a) | Plastid |
| 54 | Thr.h (abcd) <-> Gly.h (cd) + ACALD.h (ab) | Plastid |
| 55 | ACCoA.h (ab) -> ACALD.h (ab) | Plastid |
| 56 | Ser.h (abc) -> Ser.b (abc) | Plastid |
| 57 | Gly.h (ab) -> Gly.b (ab) | Plastid |
| 58 | GLX.c (ab) -> Gly.b (ab) | Cytosol |
| 59 | PYR.c (abc) + PYR.c (def) -> Val.b (daebc) + CO_2_.c (f) | Cytosol |
| 60 | PYR.c (abc) + PYR.c (def) + ACCoA.c (gh) -> Leu.b (adbech) + CO_2_.c (g) + CO_2_.c (f) | Cytosol |
| 61 | PEP.h (abc) + PEP.h (def) + E4P.h (ghij) -> chor.h (ahgibdjfce) | Plastid |
| 62 | chor.h (abcdefghij) -> Tyr.b (bdchafgei) + CO_2_.h (j) | Plastid |
| 63 | chor.h (abcdefghij) -> Phe.b (gchbdafei) + CO_2_.h (j) | Plastid |
| 64 | G6P.c (abcdef) -> Ara.b (dfbce) + CO_2_.c (a) | Cytosol |
| 65 | G6P.c (abcdef) -> Gal.b (abcdef) | Cytosol |
| 66 | G6P.h (abcdef) -> Sta.b (abcdef) | Plastid |
| 67 | ACCoA.h (ab) -> FA.b (ab) | Plastid |
| 68 | CO_2_.e (a) -> CO_2_.c (a) | Transport |
| 69 | CO_2_.c (a) <-> CO_2_.h (a) | Transport |
| 70 | CO_2_.c (a) <-> CO_2_.m (a) | Transport |
| 71 | 0*MAL.c (abcd) -> MAL.s (abcd) | Sampling Sink |
| 72 | 0*MAL.h (abcd) -> MAL.s (abcd) | Sampling Sink |
| 73 | 0*MAL.m (abcd) -> MAL.s (abcd) | Sampling Sink |
| 74 | 0*PYR.m (abc) -> PYR.s (abc) | Sampling Sink |
| 75 | 0*PYR.h (abc) -> PYR.s (abc) | Sampling Sink |
| 78 | 0*CIT.c (abcdef) -> CIT.s (abcdef) | Sampling Sink |
| 79 | 0*CIT.m (abcdef) -> CIT.s (abcdef) | Sampling Sink |
| 80 | 0*SUCC.c (abcd) -> SUCC.s (abcd) | Sampling Sink |
| 81 | 0*SUCC.m (abcd) -> SUCC.s (abcd) | Sampling Sink |
| 82 | 0.230*G6P.h + 0.678*Sta.b + 0.006*G6P.c + 0.185*Ara.b + 0.246*Gal.b + 0.116*F6P.h + 0.4*R5P.h + 0.435*T3P.h + 0.012*3PG.h + 0.136*PYR.c + 0.358*ACCoA.c + 0.023*ACCoA.h + 8.854*FA.b + 0.187*OAA.c + 0.203*SUCC.m + 0.355*Ala.b + 0.529*Gly.b + 0.748*Glu.b + 0.467*Asp.b + 0.347*Ser.b + 0.073*Phe.b + 0.037*Tyr.b + 0*Thr.b + 0.180*Val.b + 0.191*Leu.b -> Biomass | Biomass Reaction 21grD |
| 83 | 0.281*G6P.h + 0.742*Sta.b + 0.008*G6P.c + 0.202*Ara.b + 0.269*Gal.b + 0.127*F6P.h + 0.395*R5P.h + 0.532*T3P.h + 0.012*3PG.h + 0.099*PYR.c + 0.278*ACCoA.c + 0.005*ACCoA.h + 10.828*FA.b + 0.184*OAA.c + 0.203*SUCC.m + 0.268*Ala.b + 0.409*Gly.b + 0.588*Glu.b + 0.307*Asp.b + 0.154*Ser.b + 0.044*Phe.b + 0.015*Tyr.b + 0*Thr.b + 0.132*Val.b + 0.134*Leu.b -> Biomass | Biomass Reaction NK2D |
| 84 | 0.180*G6P.h + 0.325*Sta.b + 0.005*G6P.c + 0.089*Ara.b + 0.118*Gal.b + 0.056*F6P.h + 0.365*R5P.h + 0.341*T3P.h + 0.012*3PG.h + 0.221*PYR.c + 0.397*ACCoA.c + 0.100*ACCoA.h + 6.935*FA.b + 0.169*OAA.c + 0.203*SUCC.m + 0.509*Ala.b + 0.577*Gly.b + 0.878*Glu.b + 0.553*Asp.b + 0.646*Ser.b + 0.152*Phe.b + 0.084*Tyr.b + 0.485*Thr.b + 0.288*Val.b + 0.318*Leu.b -> Biomass | Biomass Reaction 21grLL |
| 85 | 0.200*G6P.h + 0.289*Sta.b + 0.005*G6P.c + 0.079*Ara.b + 0.105*Gal.b + 0.049*F6P.h + 0.473*R5P.h + 0.379*T3P.h + 0.012*3PG.h + 0.188*PYR.c + 0.464*ACCoA.c + 0.048*ACCoA.h + 7.703*FA.b + 0.223*OAA.c + 0.203*SUCC.m + 0.512*Ala.b + 0.667*Gly.b + 0.934*Glu.b + 0.763*Asp.b + 0.435*Ser.b + 0.119*Phe.b + 0.052*Tyr.b + 0.210*Thr.b + 0.261*Val.b + 0.291*Leu.b -> Biomass | Biomass Reaction NK2LL |
