## Supplemental Data 3 for "Dual-targeted NADK2 Links Mitochondrial Redox Homeostasis to Carbon Partitioning and Heterotrophic Growth in *Chlamydomonas reinhardtii*"

**pLM1-aadA-mt-roGFP2 sequence**

>pLM1-aadA-mt-roGFP2

gtgatccgtttaaactatcagtgtttgacaggatatattggcgggtaaacctaagagaaaagagcgtttattagaataatcggatatttaaaagggcgtgaaaaggtttatccgttcgtccatttgtatgtgccagccgcctttgcgacgctcaccgggctggttgccctcgccgctgggctggcggccgtctatggccctgcaaacgcgccagaaacgccgtcgaagccgtgtgcgagacaccgcggccgccggcgttgtggatacctcgcggaaaacttggccctcactgacagatgaggggcggacgttgacacttgaggggccgactcacccggcgcggcgttgacagatgaggggcaggctcgatttcggccggcgacgtggagctggccagcctcgcaaatcggcgaaaacgcctgattttacgcgagtttcccacagatgatgtggacaagcctggggataagtgccctgcggtattgacacttgaggggcgcgactactgacagatgaggggcgcgatccttgacacttgaggggcagagtgctgacagatgaggggcgcacctattgacatttgaggggctgtccacaggcagaaaatccagcatttgcaagggtttccgcccgtttttcggccaccgctaacctgtcttttaacctgcttttaaaccaatatttataaaccttgtttttaaccagggctgcgccctgtgcgcgtgaccgcgcacgccgaaggggggtgcccccccttctcgaaccctcccggcccgctaacgcgggcctcccatccccccaggggctgcgcccctcggccgcgaacggcctcaccccaaaaatggcagcgctggccaattcccgaggcacgaacccagtggacataagcctgttcggttcgtaagctgtaatgcaagtagcgtatgcgctcacgcaactggtccagaaccttgaccgaacgcagcggtggtaacggcgcagtggcggttttcatggcttgttatgactgtttttttggggtacagtctatgcctcgggcatccaagcagcaagcgcgttacgccgtgggtcgatgtttgatgttatggagcagcaacgatgttacgcagcagggcagtcgccctaaaacaaagttaaacatcatgggggaagcggtgatcgccgaagtatcgactcaactatcagaggtagttggcgtcatcgagcgccatctcgaaccgacgttgctggccgtacatttgtacggctccgcagtggatggcggcctgaagccacacagcgatattgatttgctggttacggtgaccgtaaggcttgatgaaacaacgcggcgagctttgatcaacgaccttttggaaacttcggcttcccctggagagagcgagattctccgcgctgtagaagtcaccattgttgtgcacgacgacatcattccgtggcgttatccagctaagcgcgaactgcaatttggagaatggcagcgcaatgacattcttgcaggtatcttcgagccagccacgatcgacattgatctggctatcttgctgacaaaagcaagagaacatagcgttgccttggtaggtccagcggcggaggaactctttgatccggttcctgaacaggatctatttgaggcgctaaatgaaaccttaacgctatggaactcgccgcccgactgggctggcgatgagcgaaatgtagtgcttacgttgtcccgcatttggtacagcgcagtaaccggcaaaatcgcgccgaaggatgtcgctgccgactgggcaatggagcgcctgccggcccagtatcagcccgtcatacttgaagctagacaggcttatcttggacaagaagaagatcgcttggcctcgcgcgcagatcagttggaagaatttgtccattacgtgaaaggcgagatcaccaaggtagtcggcaaataatgtctagctagaaattcgttcaagccgacgccgcttcgcggcgcggcttaactcaagcgttagatgcactaagcacataattgctcacagccaaactatcaggtcaagtctgcttttattatttttaagcgtgcataataagccctacacaaattgggagatatatcatgctgtcagaccaagtttactcatatatactttagattgatttaaaacttcatttttaatttaaaaggatctaggtgaagatcctttttgataatctcatgaccaaaatcccttaacgtgagttttcgttccactgagcgtcagaccccgtagaaaagatcaaaggatcttcttgagatcctttttttctgcgcgtaatctgctgcttgcaaacaaaaaaaccaccgctaccagcggtggtttgtttgccggatcaagagctaccaactctttttccgaaggtaactggcttcagcagagcgcagataccaaatactgtccttctagtgtagccgtagttaggccaccacttcaagaactctgtagcaccgcctacatacctcgctctgctaatcctgttaccagtggctgctgccagtggcgataagtcgtgtcttaccgggttggactcaagacgatagttaccggataaggcgcagcggtcgggctgaacggggggttcgtgcacacagcccagcttggagcgaacgacctacaccgaactgagatacctacagcgtgagctatgagaaagcgccacgcttcccgaagggagaaaggcggacaggtatccggtaagcggcagggtcggaacaggagagcgcacgagggagcttccagggggaaacgcctggtatctttatagtcctgtcgggtttcgccacctctgacttgagcgtcgatttttgtgatgctcgtcaggggggcggagcctatggaaaaacgccagcaacgcggcctttttacggttcctggcagatcctagatgtggcgcaacgatgccggcgacaagcaggagcgcaccgacttcttccgcatcaagtgttttggctctcaggccgaggcccacggcaagtatttgggcaaggggtcgctggtattcgtgcagggcaagattcggaataccaagtacgagaaggacggccagacggtctacgggaccgacttcattgccgataaggtggattatctggacaccaaggcaccaggcgggtcaaatcaggaataagggcacattgccccggcgtgagtcggggcaatcccgcaaggagggtgaatgaatcggacgtttgaccggaaggcatacaggcaagaactgatcgacgcggggttttccgccgaggatgccgaaaccatcgcaagccgcaccgtcatgcgtgcgccccgcgaaaccttccagtccgtcggctcgatggtccagcaagctacggccaagatcgagcgcgacagcgtgcaactggctccccctgccctgcccgcgccatcggccgccgtggagcgttcgcgtcgtcttgaacaggaggcggcaggtttggcgaagtcgatgaccatcgacacgcgaggaactatgacgaccaagaagcgaaaaaccgccggcgaggacctggcaaaacaggtcagcgaggccaagcaggccgcgttgctgaaacacacgaagcagcagatcaaggaaatgcagctttccttgttcgatattgcgccgtggccggacacgatgcgagcgatgccaaacgacacggcccgctctgccctgttcaccacgcgcaacaagaaaatcccgcgcgaggcgctgcaaaacaaggtcattttccacgtcaacaaggacgtgaagatcacctacaccggcgtcgagctgcgggccgacgatgacgaactggtgtggcagcaggtgttggagtacgcgaagcgcacccctatcggcgagccgatcaccttcacgttctacgagctttgccaggacctgggctggtcgatcaatggccggtattacacgaaggccgaggaatgcctgtcgcgcctacaggcgacggcgatgggcttcacgtccgaccgcgttgggcacctggaatcggtgtcgctgctgcaccgcttccgcgtcctggaccgtggcaagaaaacgtcccgttgccaggtcctgatcgacgaggaaatcgtcgtgctgtttgctggcgaccactacacgaaattcatatgggagaagtaccgcaagctgtcgccgacggcccgacggatgttcgactatttcagctcgcaccgggagccgtacccgctcaagctggaaaccttccgcctcatgtgcggatcggattccacccgcgtgaagaagtggcgcgagcaggtcggcgaagcctgcgaagagttgcgaggcagcggcctggtggaacacgcctgggtcaatgatgacctggtgcattgcaaacgctagggccttgtggggtcagttccggctgggggttcagcagcccctgctcggatctgttggaccggacagtagtcatggttgatgggctgcctgtatcgagtggtgattttgtgccgagctgccggtcggggagctgttggctggctggtggcaggatatattgtggtgtaaacaaattgacgcttagacaacttaataacacattgcggacgtttttaatgtactggggttgaacactctgtgggtctcatgccgaattcggatccggagCTGGCACTTTCTTGCGCTATGACACTTCCAGCAAAAGGTAGGGCGGGCTGCGAGACGGCTTCCCGGCGCTGCATGCAACACCGATGATGCTTCGACCCCCCGAAGCTCCTTCGGGGCTGCATGGGCGCTCCGATGCCGCTCCAGGGCGAGCGCTGTTTAAATAGCCAGGCCCCCGATTGCAAAGACATTATAGCGAGCTACCAAAGCCATATTCAAACACCTAGATCACTACCACTTCTACACAGGCCACTCGAGCTTGTGATCGCACTCCGCTAAGGGGGCGCCTCTTCCTCTTCGTTTCAGTCACAACCCGCAAAaatgGCCATGGCGGCCGTGATCGCCAAGTCGAGCGTGAGCGCCGCCGTGGCCCGCCCCGCCCGCTCCAGCGTGCGCCCCATGGCCGCCTTGAAGCCCGCCGTGAAGGCCGCGCCCGTGGCCGCGCCCGCCCAAGCCAACCAGCAATTGATGGCCATGCGCACCCCGGAGGAGCTGTCCAACCTGATTAAGGATCTGATCGAGCAGTACACTCCCGAGGTCAAGATGTCGATGGCTCGGGAGGCCGTGATTGCGGAGGgCTCGACCCAGCTGAGCGAAGTCGTGGGCGTCATCGAGCGCCACCTGGAGCCCACCCTGCTGGCCGTGCACCTGTACGGCTCCGCCGTGGACGGGGGCCTGAAGCCCCACTCGGACATCGACCTGCTCGTGACCGTGACCGTGCGCCTGGACGAGACTACTCGCCGGGCTCTCATCAACGACCTGCTGGAAACGAGCGCGTCGCCTGGCGAGTCGGAGATCCTGCGCGCCGTGGAAGTCACCATCGTCGTGCATGACGACATTATCCCCTGGCGCTACCCGGCCAAGCGCGAGCTGCAATTCGGCGAGTGGCAGCGCAACGACATCCTGGCCGGCATCTTCGAGCCCGCGACCATTGACATCGACCTGGCGATCCTCCTGACGAAGGCCCGCGAGCACTCCGTGGCGCTCGTCGGCCCGGCGGCGGAGGAGCTGTTTGACCCCGTGCCGGAGCAGGACCTGTTCGAGGCTCTGAACGAaACCCTGACGCTGTGGAACTCCCCTCCGGATTGGGCCGGCGACGAGCGGAACGTCGTGCTGACCCTGAGCCGCATTTGGTATTCGGCGGTCACCGGCAAGATCGCCCCCAAGGACGTGGCGGCGGACTGGGCCATGGAGCGGCTGCCGGCGCAATACCAGCCCGTGATCCTGGAGGCCCGGCAAGCCTACCTCGGGCAGGAGGAGGACCGCCTGGCGAGCCGGGCGGACCAGCTGGAGGAGTTCGTGCACTACGTCAAGGGCGAGATCACGAAGGTCGTGGGCAGTATCTAGgcttagcagctggaccgcctgtaccatggagaagagctttacttgccgggatggccgatttcgctgattgatacgggatcggagctcggaggctttcgcgctaggggctaggcgaagggcagtggtgaccagggtcggtgtggggtcggcccacggtcaattagccacaggaggatcagggggaggtaggcacgtcgacttggtttgcgaccccgcagttttggcggacgtgctgttgtagatgttagcgtgtgcgtgagccagtggccaacgtgccacacccattgagatgaccaaccaacttactggcaatatctgccaatgccatactgcatgtaatggccaggccatgtgagagtttgccgtgcctgcgcgcgccccgggggcgcagtttagctgaccagccgtgggatgatgcacgcatttgcaaggacagggtaatcacagcagcaacatggtgggcttaggacagctgtgggtcagtggacggacggcaggggagggacggcgcagctcgggagacagggggagacagcgtgactgtgcaatgccgctgcaagaattcaagcttggaggctgaggcttgacatgattggtgcgtatgtttgtatgaagctacaggactgatttggcgggctatgagggcgggggaagctctggaagggccgcgatggggcgcgcggcgtccagaaggcgccatacggcccgctggcggcacccatccggtataaaagcccgcgaccccgaacggtgacctccactttcagcgacaaacgagcacttatacatacgcgactattctgccgctatacataaccactcagctagcttaagatcccatcaagcttgcatgccgggcgcgccagaaggagcgcagccaaaccaggatgatgtttgatggggtatttgagcacttgcaacccttatccggaagccccctggcccacaaaggctaggcgccaatgcaagcagttcgcatgcagcccctggagcggtgccctcctgataaaccggccagggggcctatgttctttacttttttacaagagaagtcactcaacatcttaaaccatGCTGCAGCGCTCGGGAGGTCGCGTGGTGGCGGGGCTACTTCAGgtgcgactcaataactgatctgggcgagcgagtcagggggcctcgaaacctctgggcacgtccatcgtgtgggcgtcggcgcctcacgtctttgcaactcgcttgcgtcagctcacaccgcgtatcgatttgattggggctgcagATTGCCCGTGGATCCACTGCCAGCGATGTGGGCGCGCTTCGTGGAATCAGgtatgagcgtagcgtgttgcagctcatcggtgacgaccgagttagcacgcgtacattccggcagcgcgttacgctgctgtcgtcgttcggtgtaaacgaagccttctgctcatttggatgtttttgtccgacaacccgtatgcagTGCCTCGGCGGGCGCCCTTTCGGACGCaatgCGTGAGATCGTCCACATCCAGGTGCGTTGAAGCGCTTAGCGCATTGGCTGAGGGCTAGCGCAGTCAAGGGGCGCGGGGTCGTGGCTACACCCCCGCGGCTCAATTTCAAACCTGTTTCCGACTTCGAGGCTCATCGTCGCTCCGCCTGCTTGCGCCTTTACATCCACAGGGttcgGCGGCCGCGGAATTCAGCAAGGGCGAGGAGCTGTTCACCGGCGTGGTGCCCATCCTGGTGGAGCTGGACGGCGACGTGAACGGCCACAAGTTCAGCGTGAGCGGCGAGGGCGAGGGCGACGCCACCTACGGCAAGCTGACCCTGAAGTTCATCAGCACCACCGGCAAGCTGCCCGTGCCCTGGCCCACCCTGGTGACCACCCTGACCTACGGCGTGCAGTGCTTCAGCCGCTACCCCGACCACATGAAGCGCCACGACTTCTTCAAGAGCGCCATGCCCGAGGGCTACGTGCAGGAGCGCACCATCTTCTTCAAGGACGACGGCAACTACAAGACCCGCGCCGAGGTGAAGTTCGAGGGCGACACCCTGGTGAACCGCATCGAGCTGAAGGGCATCGACTTCAAGGAGGACGGCAACATCCTGGGCCACAAGCTGGAGTACAACTACAACTGCCACAACGTGTACATCATGGCCGACAAGCAGAAGAACGGCATCAAGGTGAACTTCAAGATCCGCCACAACATCGAGGACGGCAGCGTGCAGCTGGCCGACCACTACCAGCAGAACACCCCCATCGGCGACGGCCCCGTGCTGCTGCCCGACAACCACTACCTGAGCACCTGCAGCGCCCTGAGCAAGGACCCCAACGAGAAGCGCGACCACATGGTGCTGCTGGAGTTCGTGACCGCCGCCGGCATCACCCACGGCATGGACGAGCTGTACAAGTAAGCTTagcagctggaccgcctgtaccatggagaagagctttacttgccgggatggccgatttcgctgattgatacgggatcggagctcggaggctttcgcgctaggggctaggcgaagggcagtggtgaccagggtcggtgtggggtcggcccacggtcaattagccacaggaggatcagggggaggtaggcacgtcgacttggtttgcgaccccgcagttttggcggacgtgctgttgtagatgttagcgtgtgcgtgagccagtggccaacgtgccacacccattgagatgaccaaccaacttactggcaatatctgccaatgccatactgcatgtaatggccaggccatgtgagagtttgccgtgcctgcgcgcgccccgggggcgcagtttagctgaccagccgtgggatgatgcacgcatttgcaaggacagggtaatcacagcagcaacatggtgggcttaggacagctgtgggtcagtggacggacggcaggggagggacggcgcagctcgggagacagggggagacagcgtgactgtgcaatgccgctactagaggatgcacatgtgaccgagggacacgaa

SpecR

Ori

pBetaTub2

aadA

tPSAD

pAR

mtHSP70C

beta-Tub(i)

roGFP2

tPSAD
