## Supplemental Figures for "Dual-targeted NADK2 Links Mitochondrial Redox Homeostasis to Carbon Partitioning and Heterotrophic Growth in *Chlamydomonas reinhardtii*"

### Slide 1
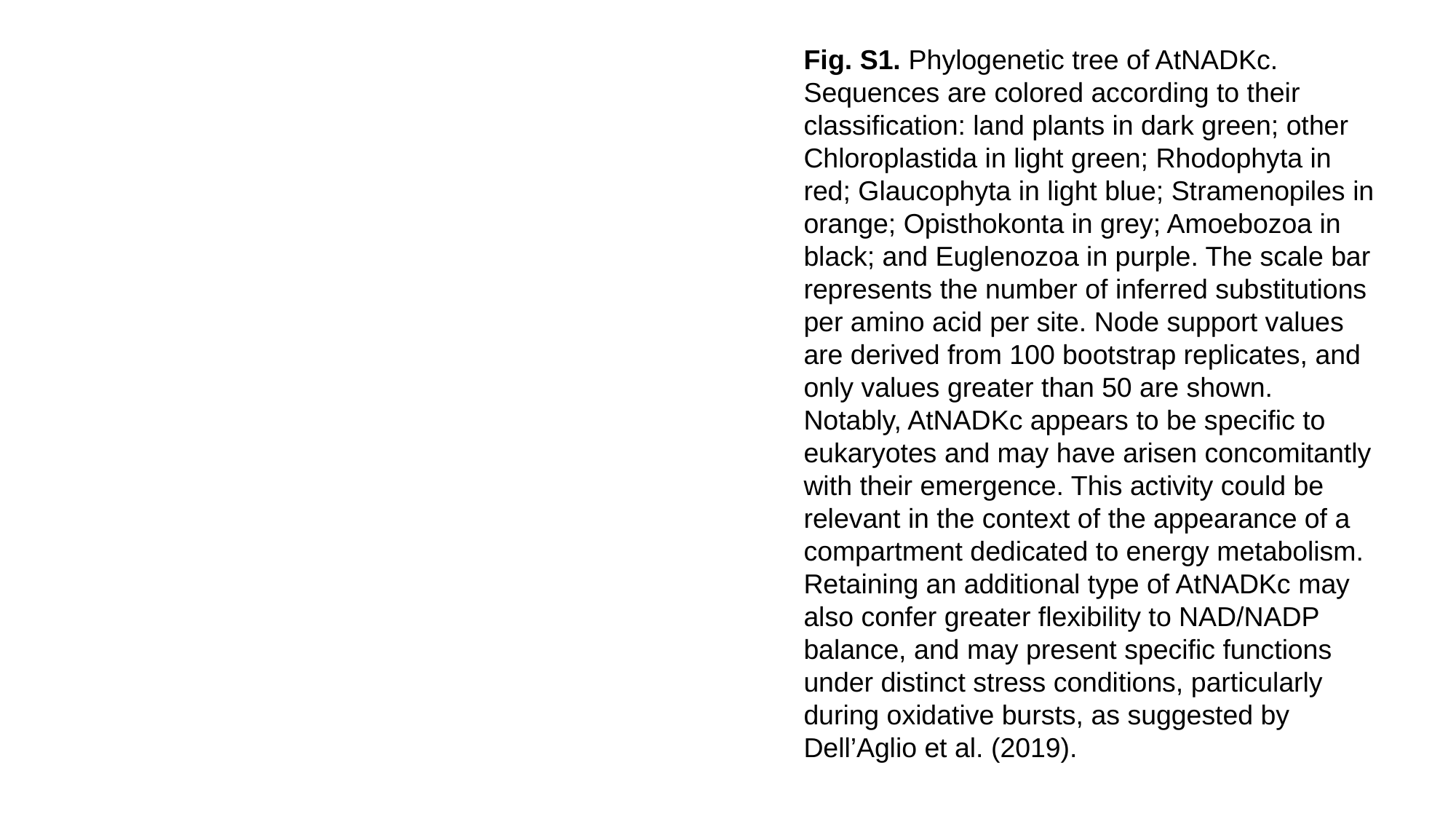

Fig. S1. Phylogenetic tree of AtNADKc. Sequences are colored according to their classification: land plants in dark green; other Chloroplastida in light green; Rhodophyta in red; Glaucophyta in light blue; Stramenopiles in orange; Opisthokonta in grey; Amoebozoa in black; and Euglenozoa in purple. The scale bar represents the number of inferred substitutions per amino acid per site. Node support values are derived from 100 bootstrap replicates, and only values greater than 50 are shown. Notably, AtNADKc appears to be specific to eukaryotes and may have arisen concomitantly with their emergence. This activity could be relevant in the context of the appearance of a compartment dedicated to energy metabolism. Retaining an additional type of AtNADKc may also confer greater flexibility to NAD/NADP balance, and may present specific functions under distinct stress conditions, particularly during oxidative bursts, as suggested by Dell’Aglio et al. (2019).

### Slide 2
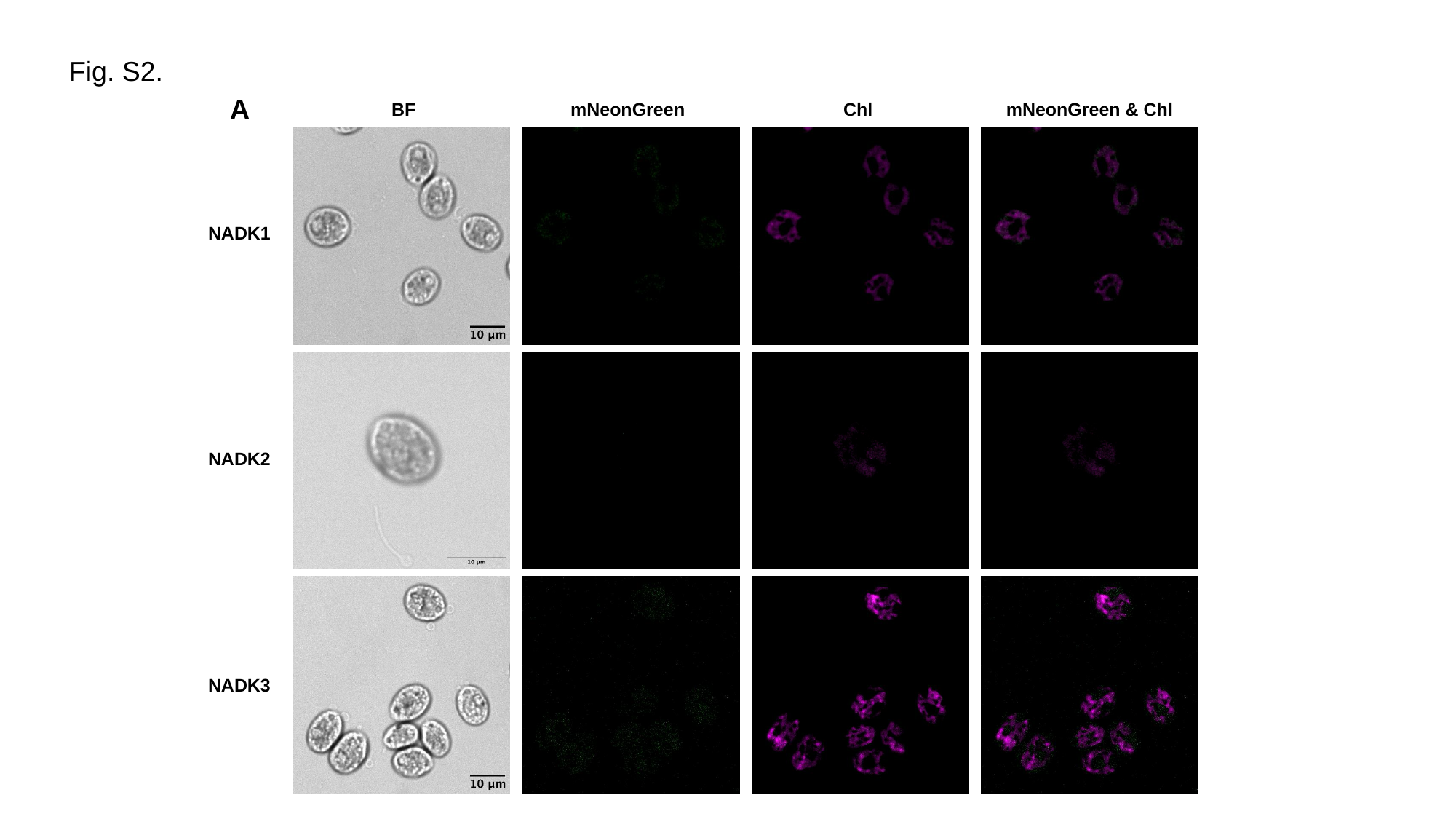

Fig. S2.
A
BF
mNeonGreen
Chl
mNeonGreen & Chl
NADK1
NADK2
NADK3

### Slide 3
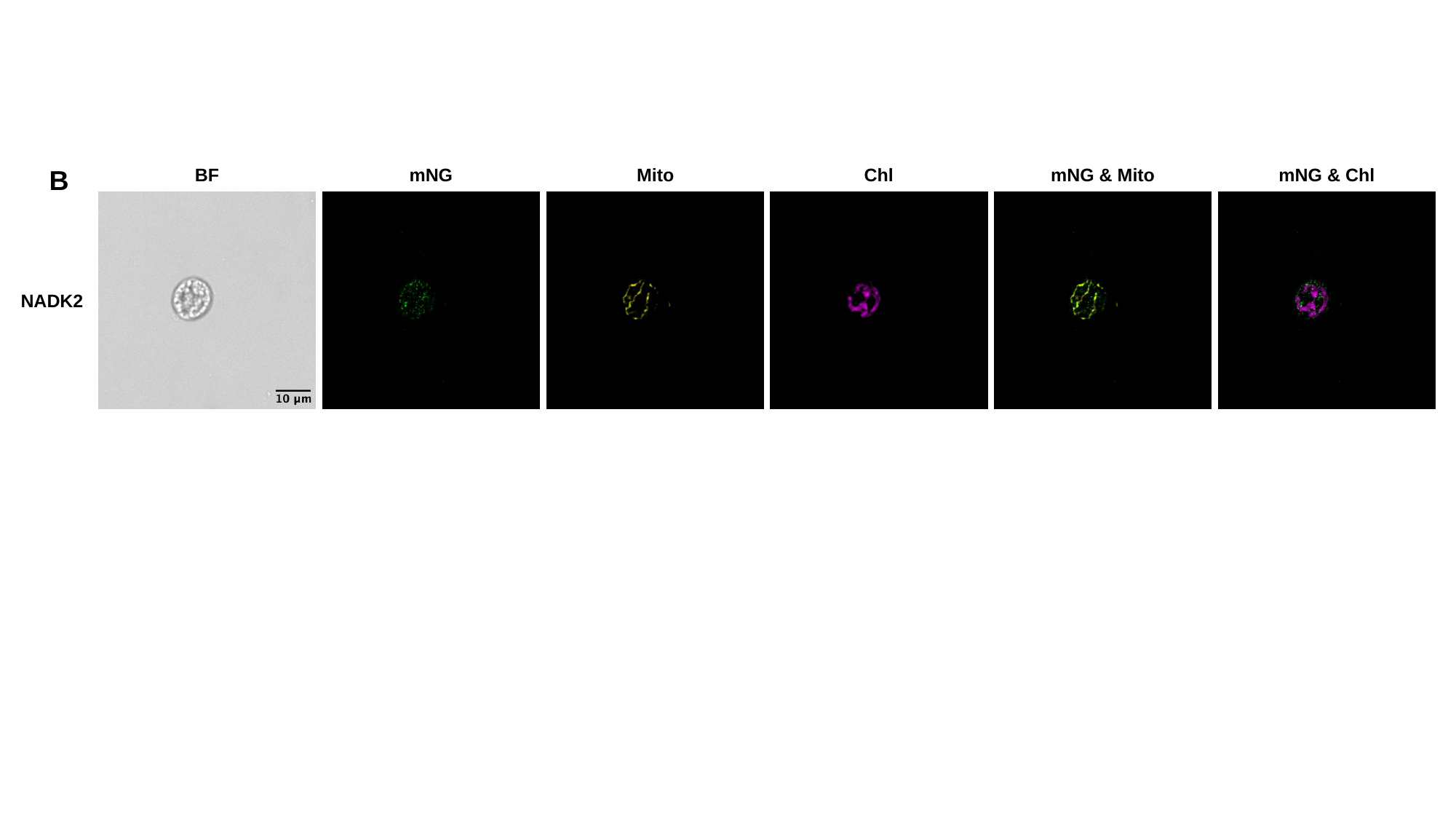

B
mNG & Mito
mNG & Chl
Chl
mNG
BF
Mito
NADK2

### Slide 4
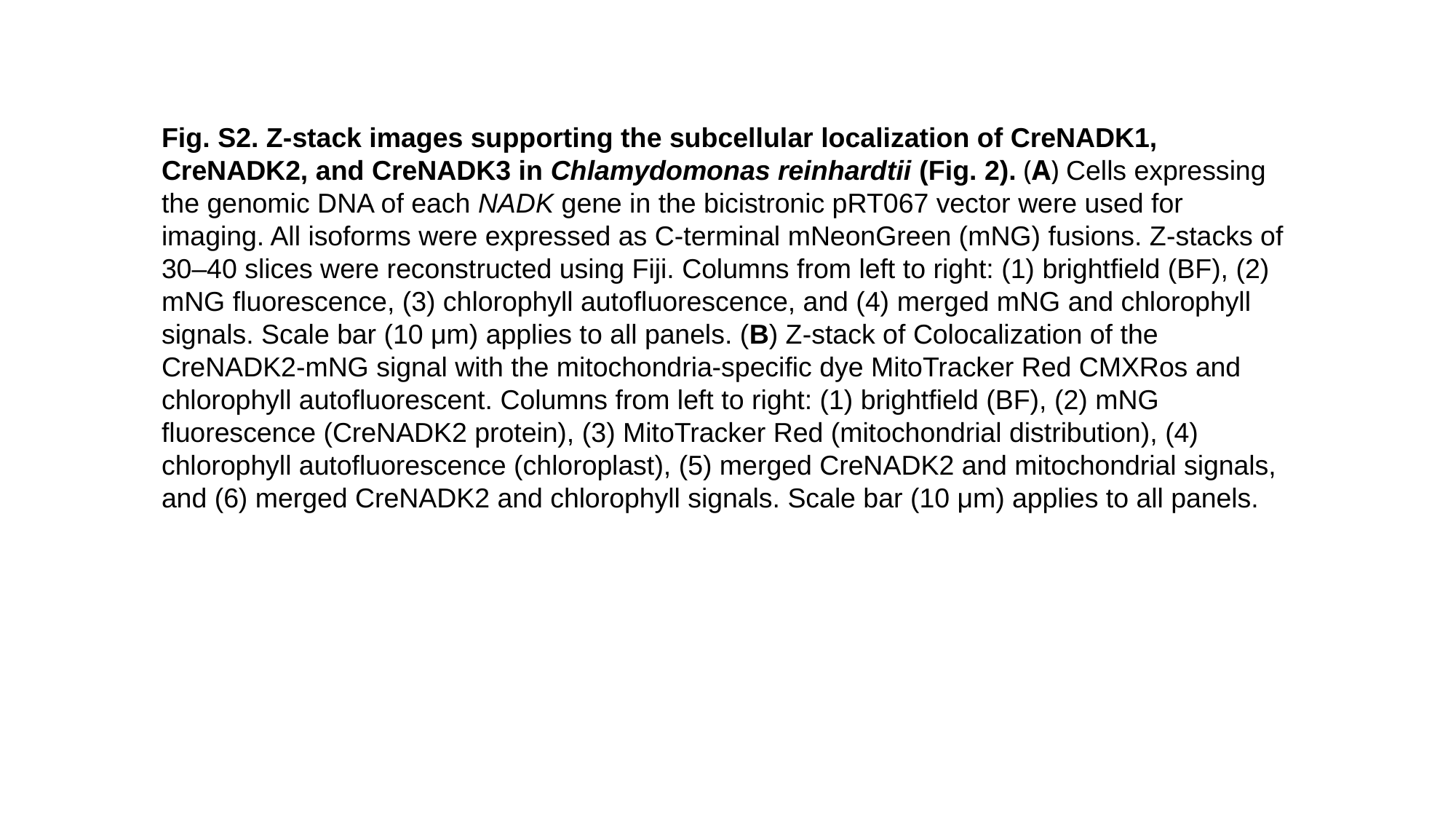

Fig. S2. Z-stack images supporting the subcellular localization of CreNADK1, CreNADK2, and CreNADK3 in Chlamydomonas reinhardtii (Fig. 2). (A) Cells expressing the genomic DNA of each NADK gene in the bicistronic pRT067 vector were used for imaging. All isoforms were expressed as C-terminal mNeonGreen (mNG) fusions. Z-stacks of 30–40 slices were reconstructed using Fiji. Columns from left to right: (1) brightfield (BF), (2) mNG fluorescence, (3) chlorophyll autofluorescence, and (4) merged mNG and chlorophyll signals. Scale bar (10 μm) applies to all panels. (B) Z-stack of Colocalization of the CreNADK2-mNG signal with the mitochondria-specific dye MitoTracker Red CMXRos and chlorophyll autofluorescent. Columns from left to right: (1) brightfield (BF), (2) mNG fluorescence (CreNADK2 protein), (3) MitoTracker Red (mitochondrial distribution), (4) chlorophyll autofluorescence (chloroplast), (5) merged CreNADK2 and mitochondrial signals, and (6) merged CreNADK2 and chlorophyll signals. Scale bar (10 μm) applies to all panels.

### Slide 5
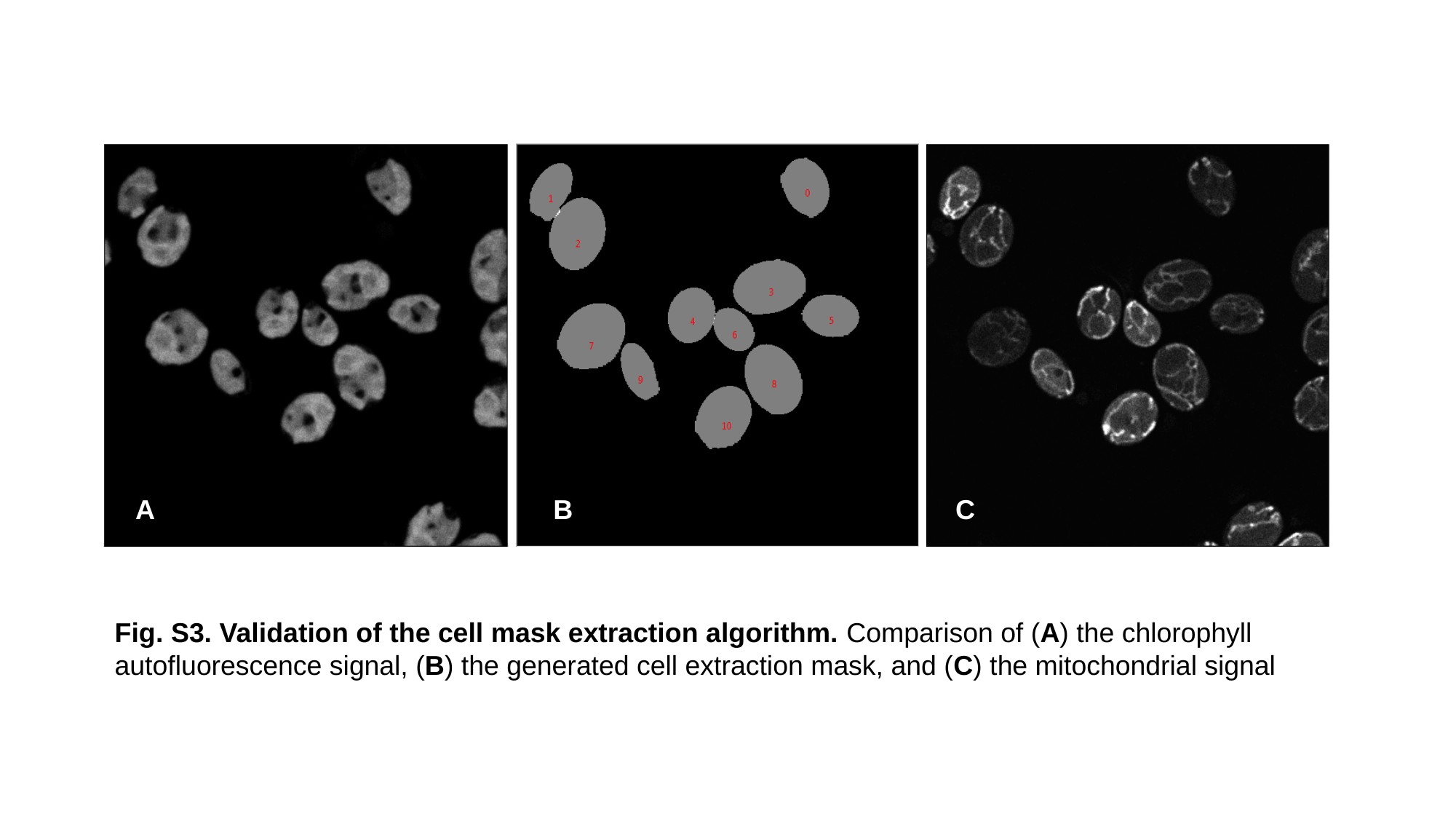

A
B
C
Fig. S3. Validation of the cell mask extraction algorithm. Comparison of (A) the chlorophyll autofluorescence signal, (B) the generated cell extraction mask, and (C) the mitochondrial signal

### Slide 6
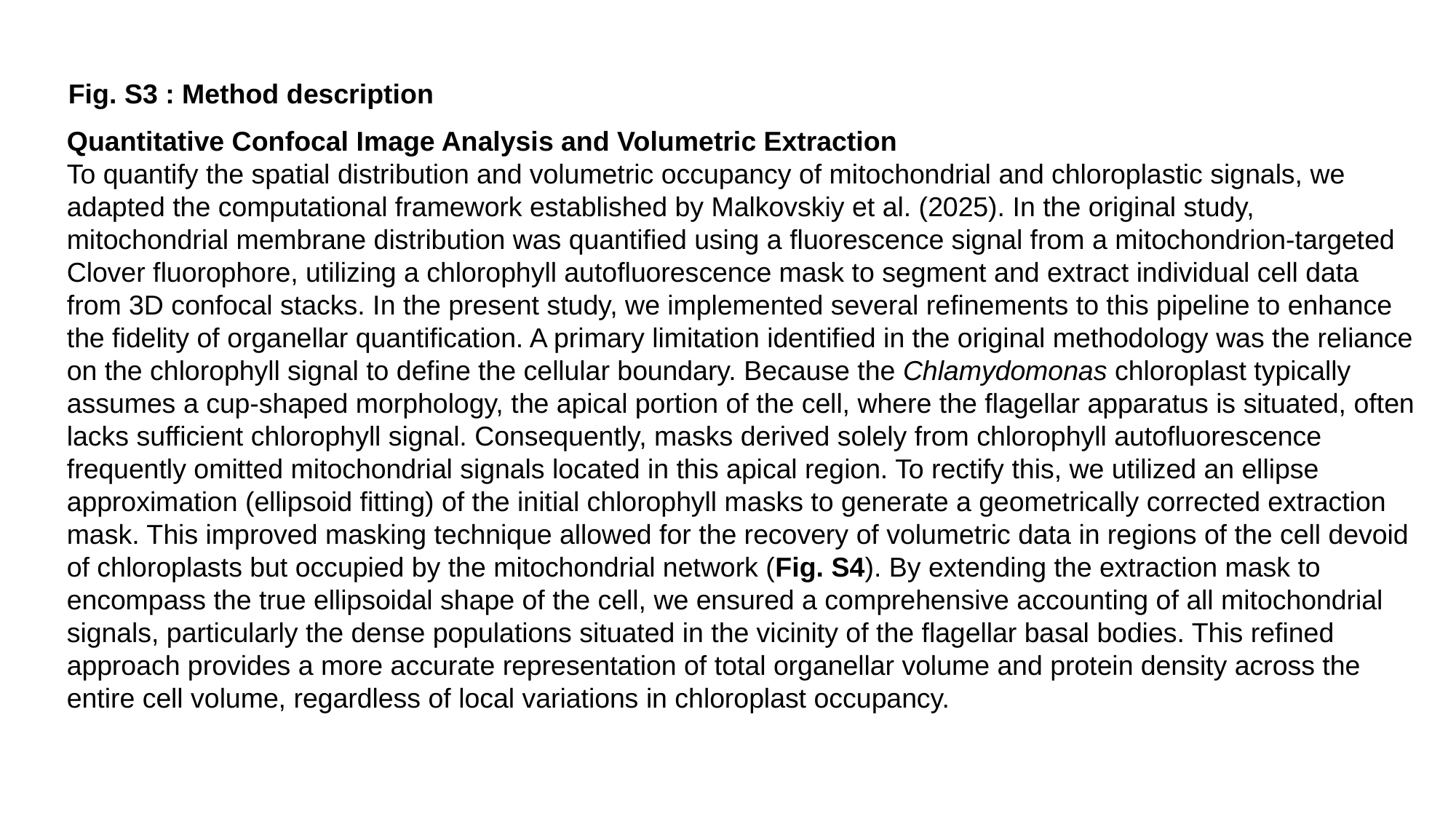

Fig. S3 : Method description
Quantitative Confocal Image Analysis and Volumetric Extraction
To quantify the spatial distribution and volumetric occupancy of mitochondrial and chloroplastic signals, we adapted the computational framework established by Malkovskiy et al. (2025). In the original study, mitochondrial membrane distribution was quantified using a fluorescence signal from a mitochondrion-targeted Clover fluorophore, utilizing a chlorophyll autofluorescence mask to segment and extract individual cell data from 3D confocal stacks. In the present study, we implemented several refinements to this pipeline to enhance the fidelity of organellar quantification. A primary limitation identified in the original methodology was the reliance on the chlorophyll signal to define the cellular boundary. Because the Chlamydomonas chloroplast typically assumes a cup-shaped morphology, the apical portion of the cell, where the flagellar apparatus is situated, often lacks sufficient chlorophyll signal. Consequently, masks derived solely from chlorophyll autofluorescence frequently omitted mitochondrial signals located in this apical region. To rectify this, we utilized an ellipse approximation (ellipsoid fitting) of the initial chlorophyll masks to generate a geometrically corrected extraction mask. This improved masking technique allowed for the recovery of volumetric data in regions of the cell devoid of chloroplasts but occupied by the mitochondrial network (Fig. S4). By extending the extraction mask to encompass the true ellipsoidal shape of the cell, we ensured a comprehensive accounting of all mitochondrial signals, particularly the dense populations situated in the vicinity of the flagellar basal bodies. This refined approach provides a more accurate representation of total organellar volume and protein density across the entire cell volume, regardless of local variations in chloroplast occupancy.

### Slide 7
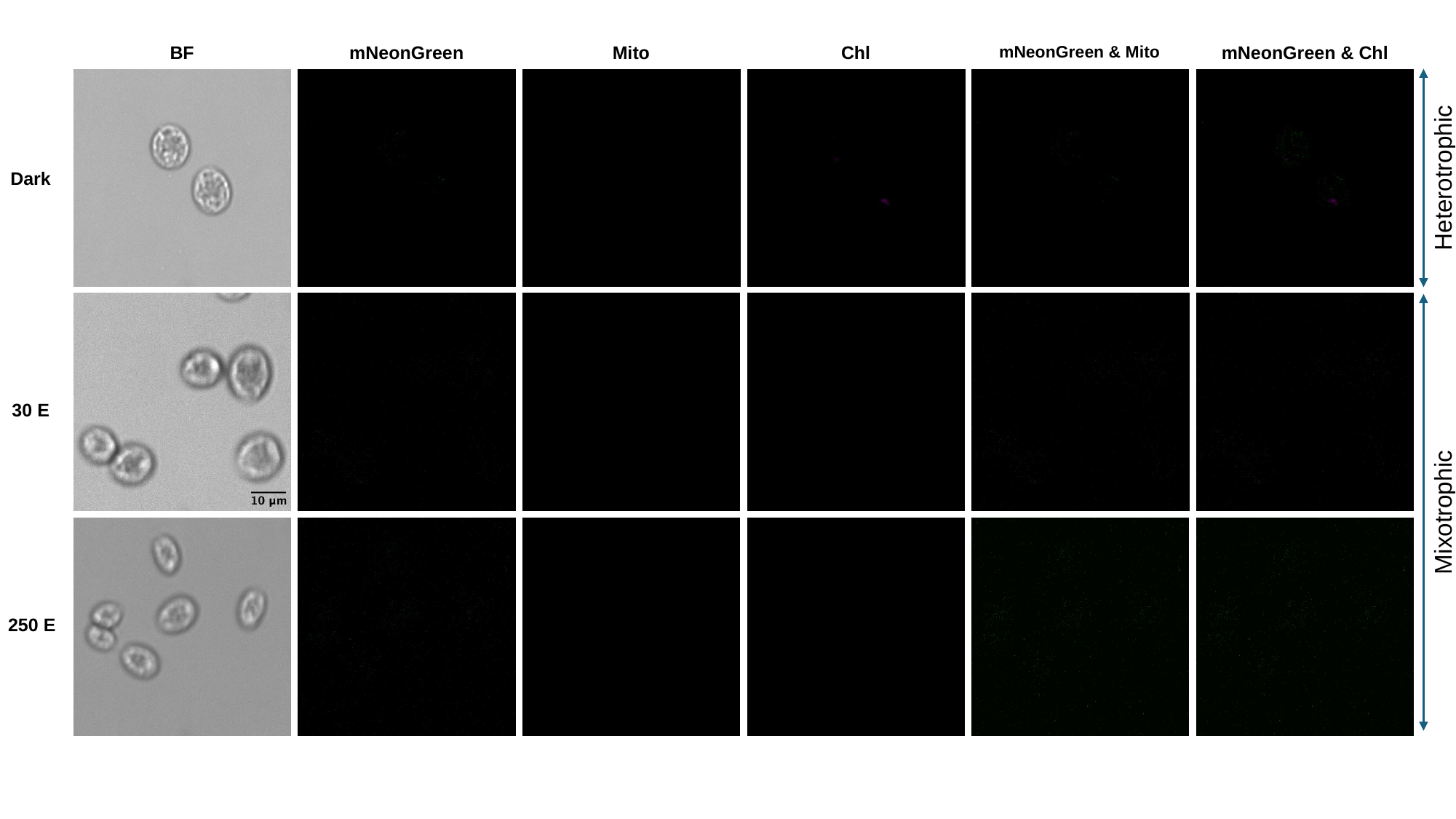

Mito
Chl
mNeonGreen & Mito
mNeonGreen & Chl
BF
mNeonGreen
Heterotrophic
Dark
Mixotrophic

### Slide 8
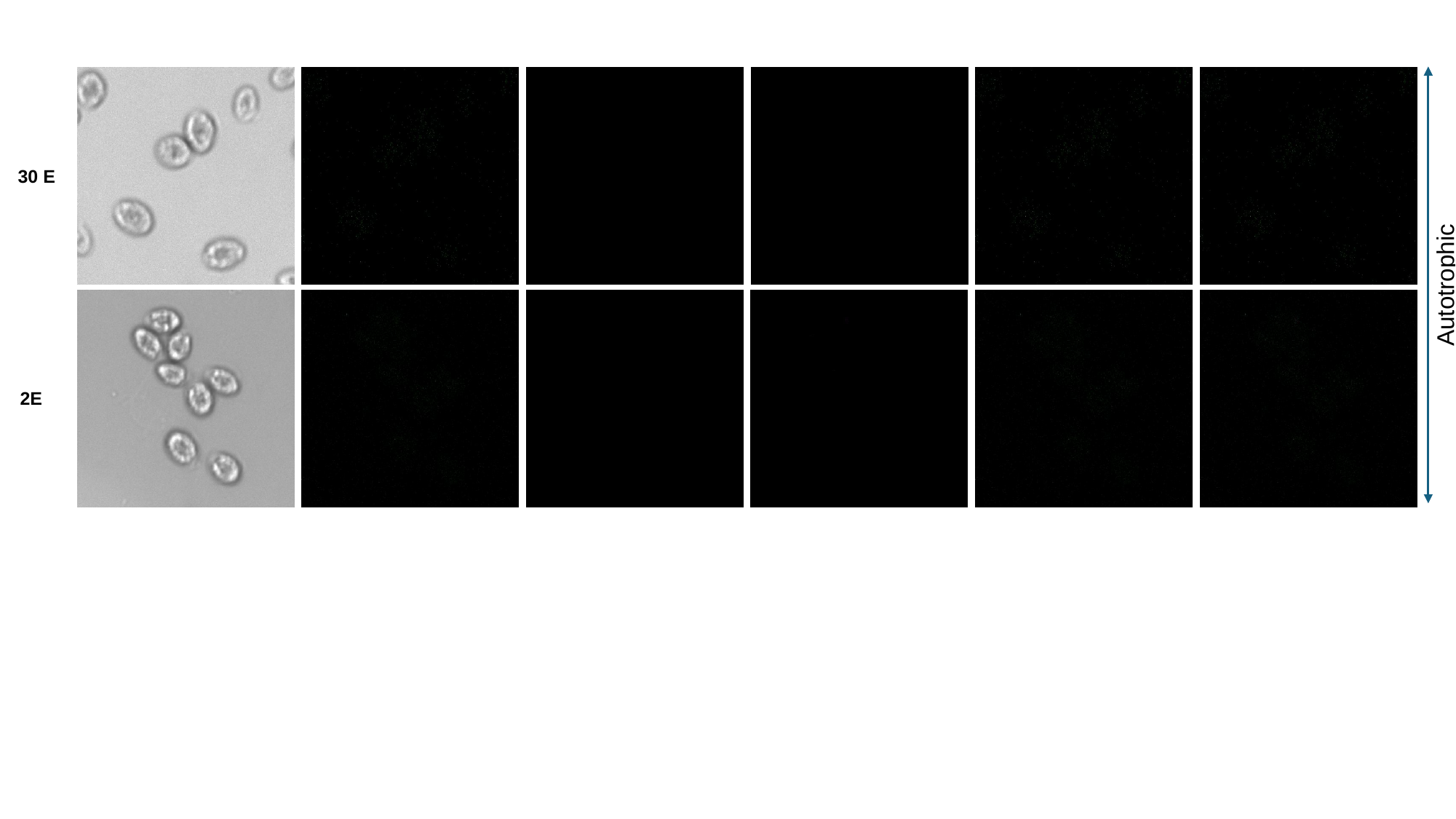

Autotrophic

### Slide 9
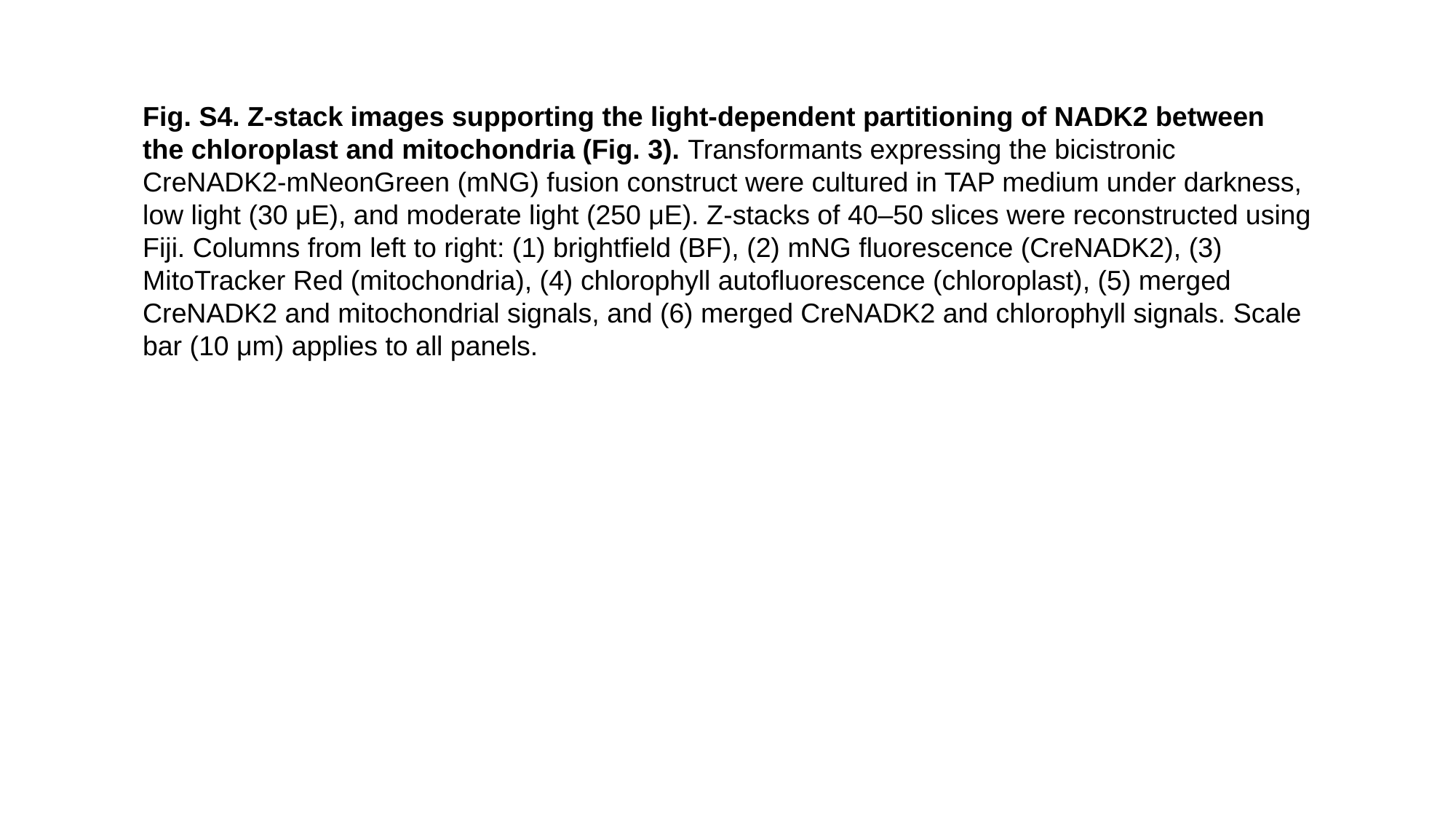

Fig. S4. Z-stack images supporting the light-dependent partitioning of NADK2 between the chloroplast and mitochondria (Fig. 3). Transformants expressing the bicistronic CreNADK2-mNeonGreen (mNG) fusion construct were cultured in TAP medium under darkness, low light (30 μE), and moderate light (250 μE). Z-stacks of 40–50 slices were reconstructed using Fiji. Columns from left to right: (1) brightfield (BF), (2) mNG fluorescence (CreNADK2), (3) MitoTracker Red (mitochondria), (4) chlorophyll autofluorescence (chloroplast), (5) merged CreNADK2 and mitochondrial signals, and (6) merged CreNADK2 and chlorophyll signals. Scale bar (10 μm) applies to all panels.

### Slide 10
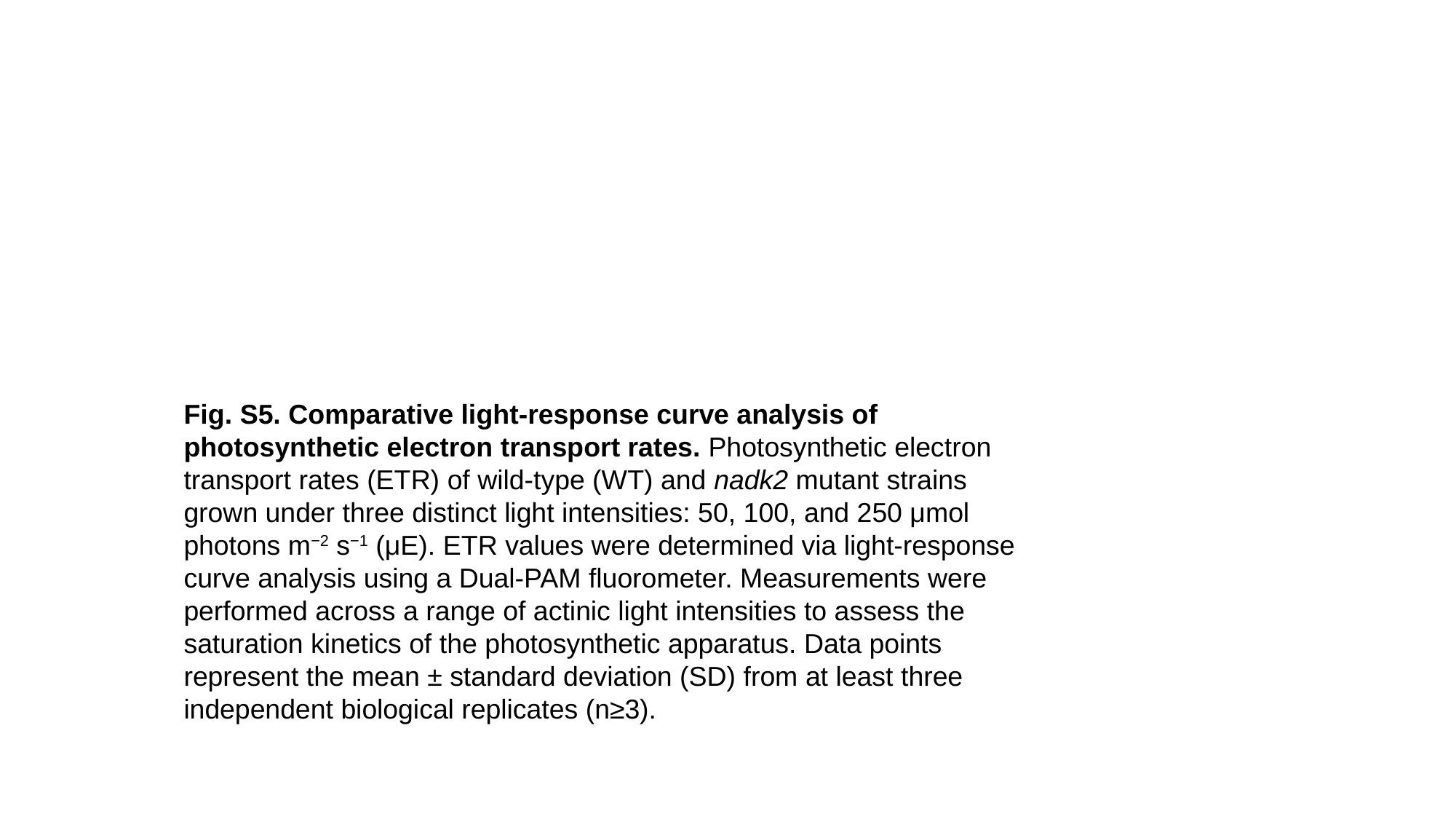

Fig. S5. Comparative light-response curve analysis of photosynthetic electron transport rates. Photosynthetic electron transport rates (ETR) of wild-type (WT) and nadk2 mutant strains grown under three distinct light intensities: 50, 100, and 250 μmol photons m^−2^ s^−1^ (μE). ETR values were determined via light-response curve analysis using a Dual-PAM fluorometer. Measurements were performed across a range of actinic light intensities to assess the saturation kinetics of the photosynthetic apparatus. Data points represent the mean ± standard deviation (SD) from at least three independent biological replicates (n≥3).

### Slide 11
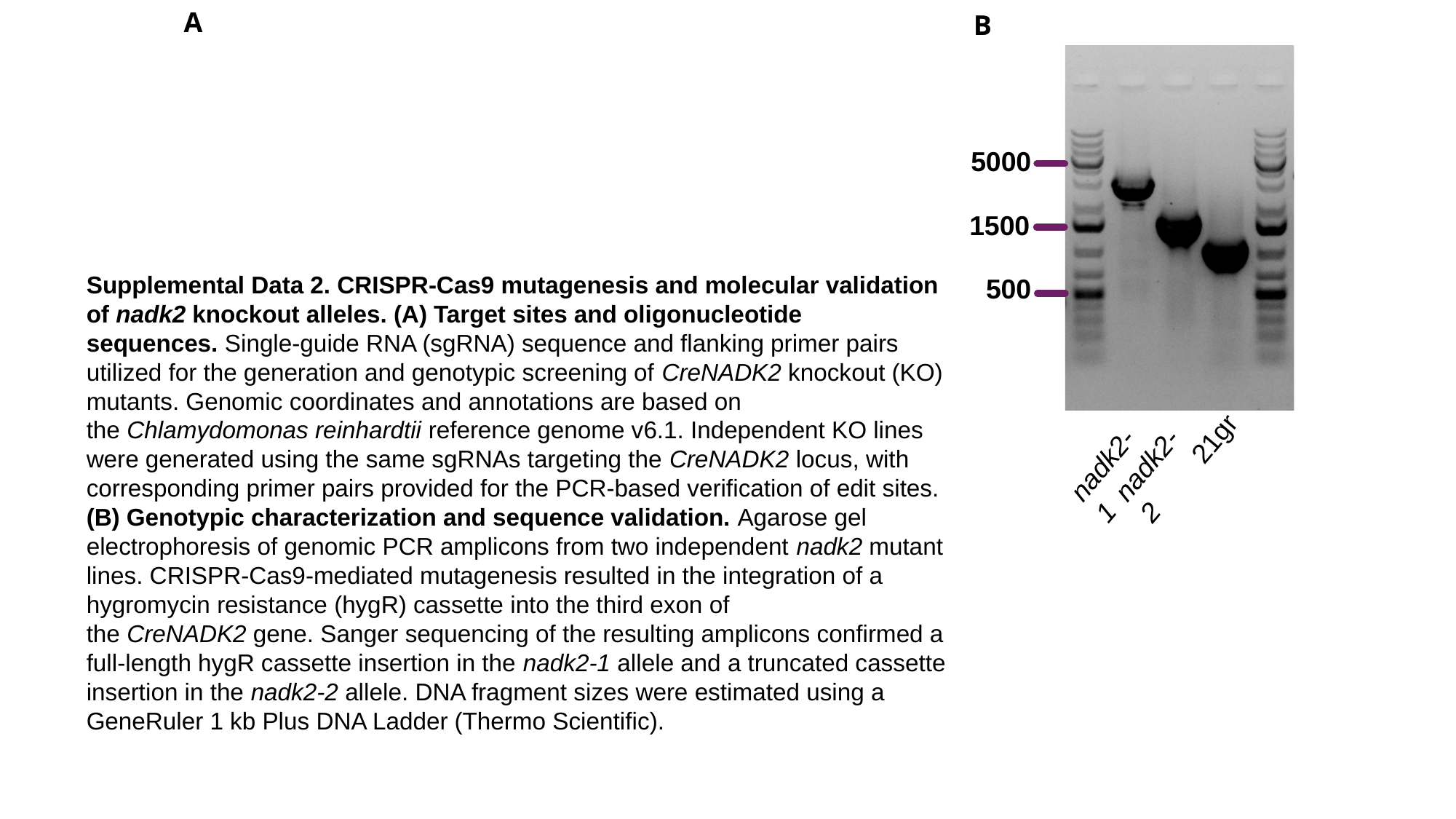

A
B
5000
1500
500
21gr
nadk2-1
nadk2-2
Supplemental Data 2. CRISPR-Cas9 mutagenesis and molecular validation of nadk2 knockout alleles. (A) Target sites and oligonucleotide sequences. Single-guide RNA (sgRNA) sequence and flanking primer pairs utilized for the generation and genotypic screening of CreNADK2 knockout (KO) mutants. Genomic coordinates and annotations are based on the Chlamydomonas reinhardtii reference genome v6.1. Independent KO lines were generated using the same sgRNAs targeting the CreNADK2 locus, with corresponding primer pairs provided for the PCR-based verification of edit sites. (B) Genotypic characterization and sequence validation. Agarose gel electrophoresis of genomic PCR amplicons from two independent nadk2 mutant lines. CRISPR-Cas9-mediated mutagenesis resulted in the integration of a hygromycin resistance (hygR) cassette into the third exon of the CreNADK2 gene. Sanger sequencing of the resulting amplicons confirmed a full-length hygR cassette insertion in the nadk2-1 allele and a truncated cassette insertion in the nadk2-2 allele. DNA fragment sizes were estimated using a GeneRuler 1 kb Plus DNA Ladder (Thermo Scientific).

### Slide 12
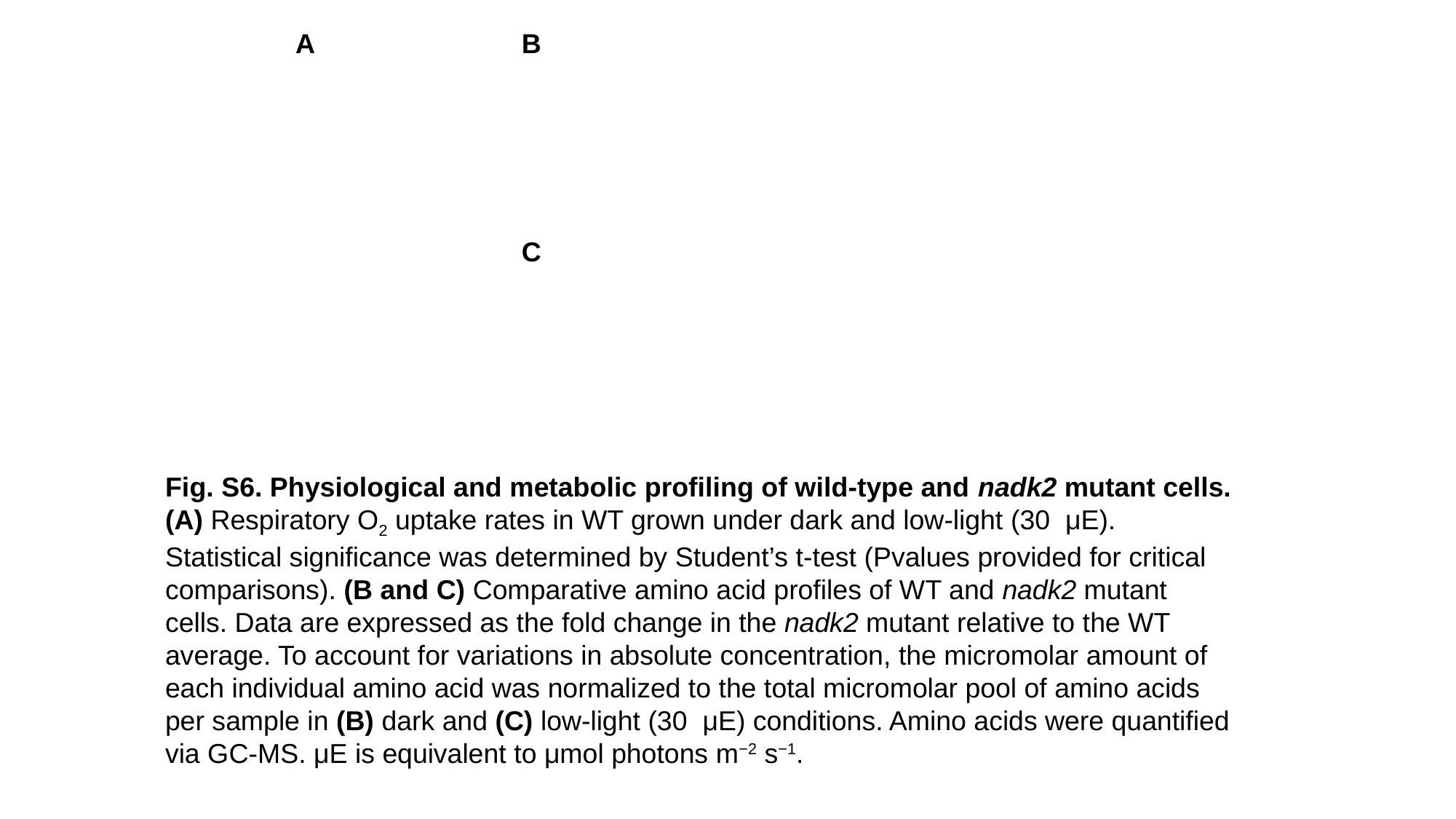

A
B
C
Fig. S6. Physiological and metabolic profiling of wild-type and nadk2 mutant cells. (A) Respiratory O_2_​ uptake rates in WT grown under dark and low-light (30  μE). Statistical significance was determined by Student’s t-test (Pvalues provided for critical comparisons). (B and C) Comparative amino acid profiles of WT and nadk2 mutant cells. Data are expressed as the fold change in the nadk2 mutant relative to the WT average. To account for variations in absolute concentration, the micromolar amount of each individual amino acid was normalized to the total micromolar pool of amino acids per sample in (B) dark and (C) low-light (30  μE) conditions. Amino acids were quantified via GC-MS. μE is equivalent to μmol photons m^−2^ s^−1^.

### Slide 13
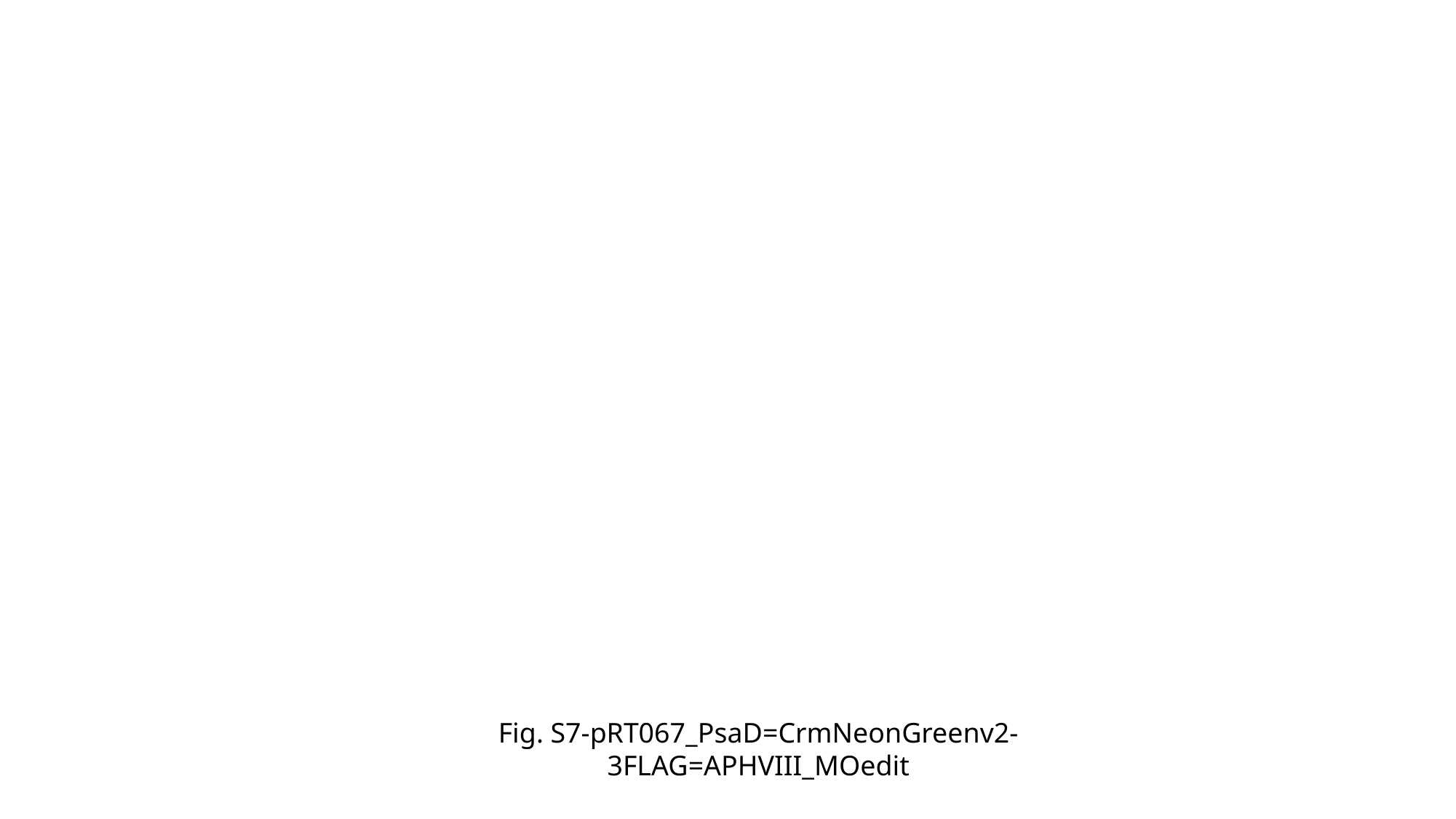

Fig. S7-pRT067_PsaD=CrmNeonGreenv2-3FLAG=APHVIII_MOedit

### Slide 14
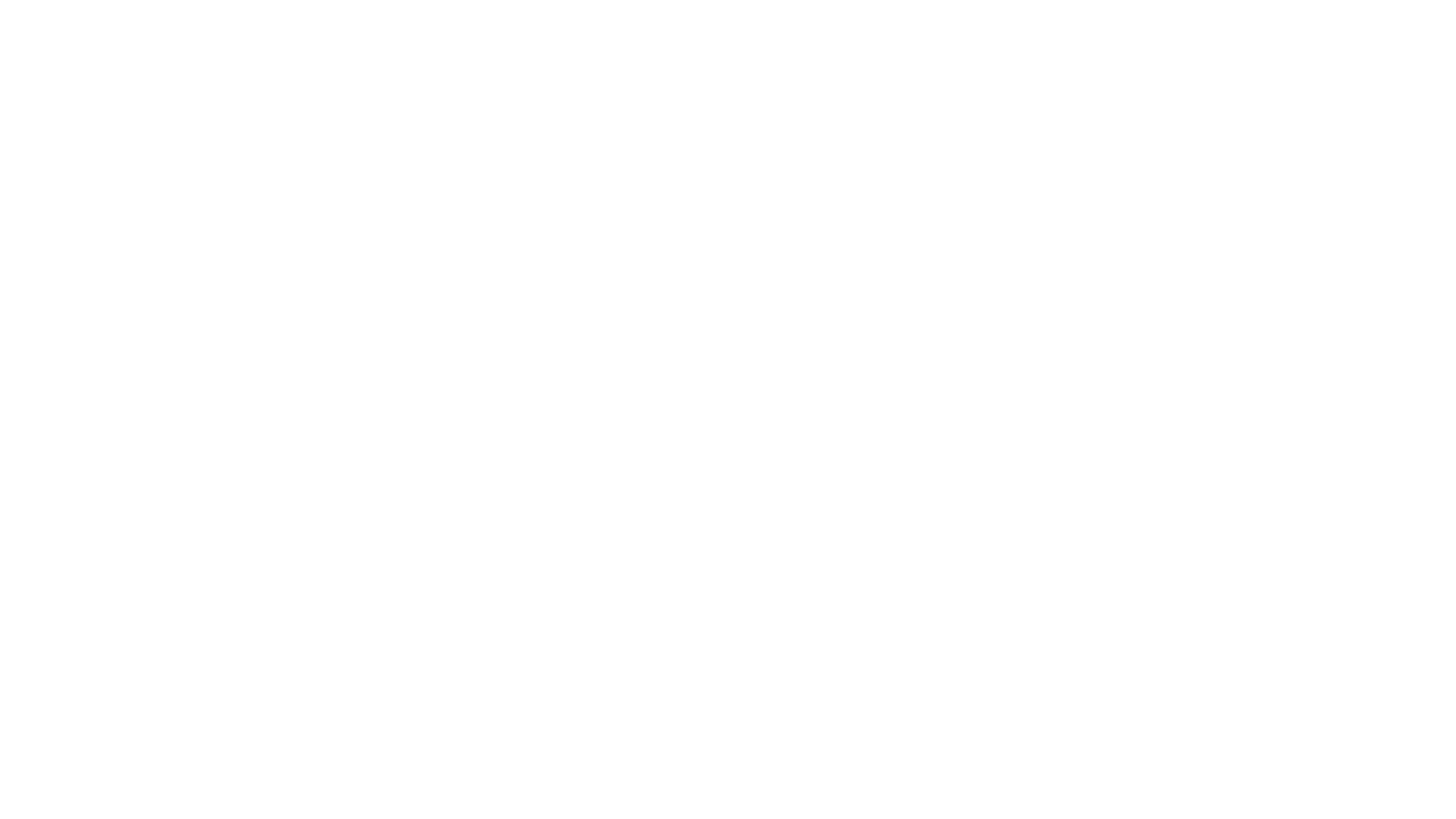

### Slide 15
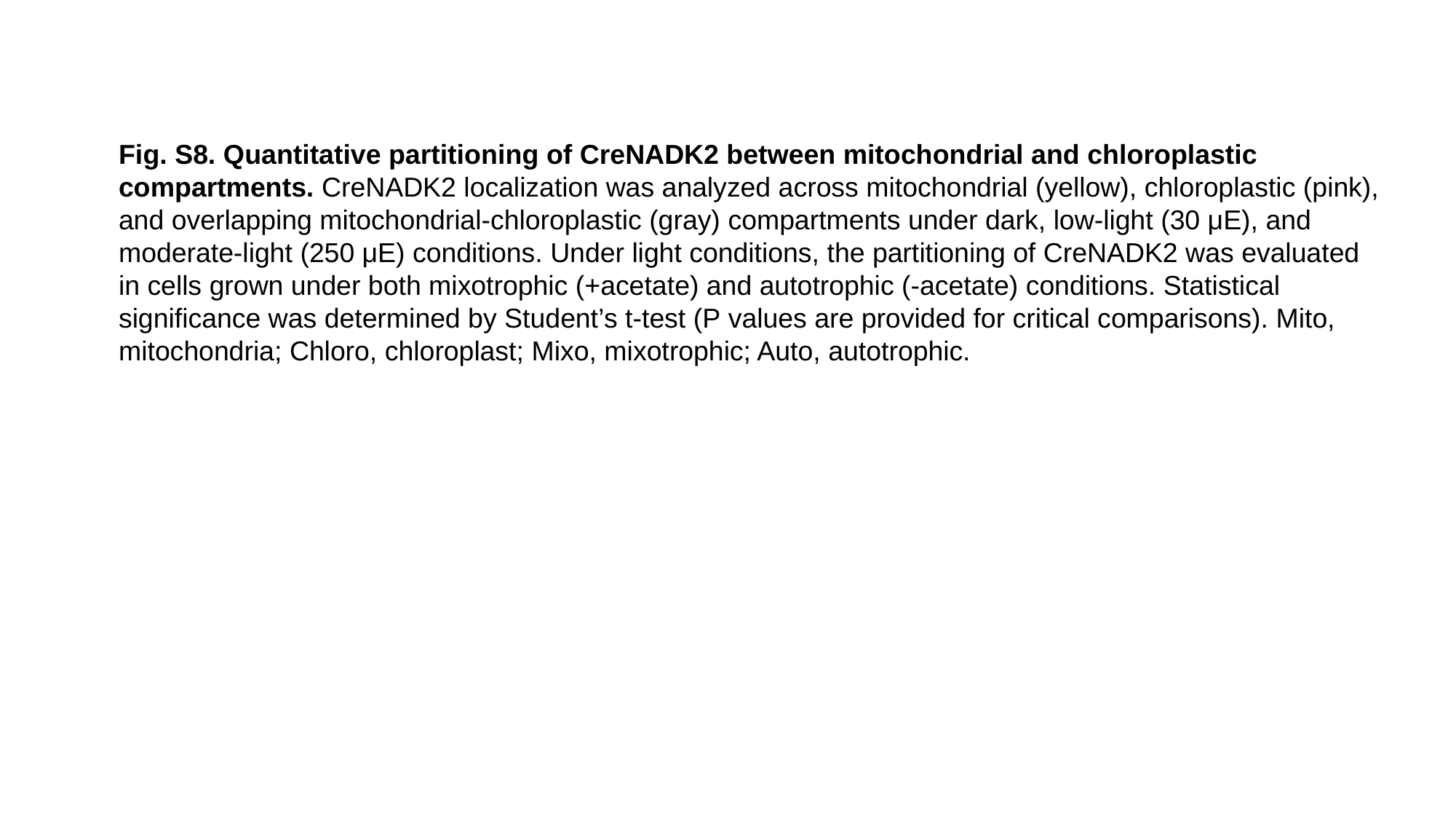

Fig. S8. Quantitative partitioning of CreNADK2 between mitochondrial and chloroplastic compartments. CreNADK2 localization was analyzed across mitochondrial (yellow), chloroplastic (pink), and overlapping mitochondrial-chloroplastic (gray) compartments under dark, low-light (30 μE), and moderate-light (250 μE) conditions. Under light conditions, the partitioning of CreNADK2 was evaluated in cells grown under both mixotrophic (+acetate) and autotrophic (-acetate) conditions. Statistical significance was determined by Student’s t-test (P values are provided for critical comparisons). Mito, mitochondria; Chloro, chloroplast; Mixo, mixotrophic; Auto, autotrophic.

### Slide 16
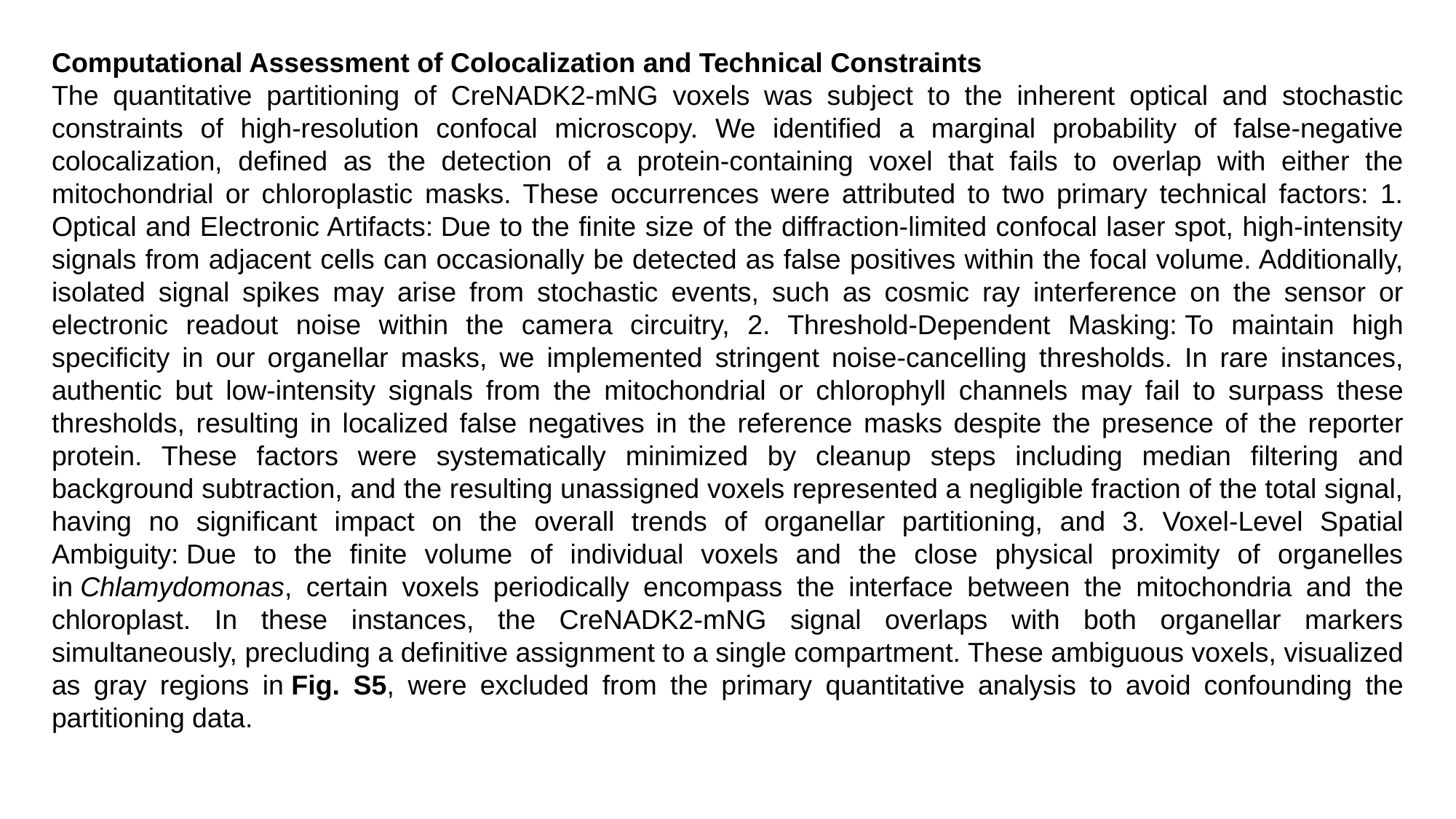

Computational Assessment of Colocalization and Technical Constraints
The quantitative partitioning of CreNADK2-mNG voxels was subject to the inherent optical and stochastic constraints of high-resolution confocal microscopy. We identified a marginal probability of false-negative colocalization, defined as the detection of a protein-containing voxel that fails to overlap with either the mitochondrial or chloroplastic masks. These occurrences were attributed to two primary technical factors: 1. Optical and Electronic Artifacts: Due to the finite size of the diffraction-limited confocal laser spot, high-intensity signals from adjacent cells can occasionally be detected as false positives within the focal volume. Additionally, isolated signal spikes may arise from stochastic events, such as cosmic ray interference on the sensor or electronic readout noise within the camera circuitry, 2. Threshold-Dependent Masking: To maintain high specificity in our organellar masks, we implemented stringent noise-cancelling thresholds. In rare instances, authentic but low-intensity signals from the mitochondrial or chlorophyll channels may fail to surpass these thresholds, resulting in localized false negatives in the reference masks despite the presence of the reporter protein. These factors were systematically minimized by cleanup steps including median filtering and background subtraction, and the resulting unassigned voxels represented a negligible fraction of the total signal, having no significant impact on the overall trends of organellar partitioning, and 3. Voxel-Level Spatial Ambiguity: Due to the finite volume of individual voxels and the close physical proximity of organelles in Chlamydomonas, certain voxels periodically encompass the interface between the mitochondria and the chloroplast. In these instances, the CreNADK2-mNG signal overlaps with both organellar markers simultaneously, precluding a definitive assignment to a single compartment. These ambiguous voxels, visualized as gray regions in Fig. S5, were excluded from the primary quantitative analysis to avoid confounding the partitioning data.
